# An atlas of neural crest lineages along the posterior developing zebrafish at single-cell resolution

**DOI:** 10.1101/2020.06.14.150938

**Authors:** Aubrey G.A. Howard, Phillip A. Baker, Rodrigo Ibarra-García-Padilla, Joshua A. Moore, Lucia J. Rivas, James J. Tallman, Eileen W. Singleton, Jessa L. Westheimer, Julia A. Corteguera, Rosa A. Uribe

## Abstract

Neural crest cells (NCCs) are vertebrate stem cells that give rise to various cell types throughout the developing body in early life. Here, we utilized single-cell transcriptomic analyses to delineate NCC-derivatives along the posterior developing vertebrate, zebrafish, during the late embryonic to early larval stage, a period when NCCs are actively differentiating into distinct cellular lineages. We identified several major NCC/NCC-derived cell-types including mesenchyme, neural crest, neural, neuronal, glial, and pigment, from which we resolved over three dozen cellular subtypes. We dissected gene expression signatures of pigment progenitors delineating into chromatophore lineages, mesenchyme subtypes, and enteric NCCs transforming into enteric neurons. Global analysis of NCC derivatives revealed they were demarcated by combinatorial *hox* gene codes, with distinct profiles within neuronal cells. From these analyses, we present a comprehensive cell-type atlas that can be utilized as a valuable resource for further mechanistic and evolutionary investigations of NCC differentiation.

## INTRODUCTION

Unique to vertebrates, neural crest cells (NCC) are an embryonic stem cell population characterized as transient, highly migratory, and multipotent. Following their birth from the dorsal neural tube, NCCs migrate extensively, dorsolaterally or ventrally along the main axial levels of the embryo; the cranial, vagal, trunk, and sacral regions (Graham et al., 2004; Le Douarin and Teillet et al., 1974). Depending on the axial level of their origination, NCCs give rise to different cell types within many critical tissues; such as the cornea, craniofacial cartilage and bone, mesenchyme, pigment cells in the skin, as well as neurons and glia that comprise peripheral ganglia (Hutchins et al., 2018; Epstein et al., 1994; Kuo and Erickson, 2011; Hall and Hörstadius, 1988; Le Douarin and Kalcheim, 1999; Theveneau and Mayor, 2012; Williams and Bohnsack, 2015; Yntema and Hammond, 1954).

During their development, NCCs undergo dramatic transcriptional changes which lead to diverse cellular lineages, making their transcriptomic profiles highly dynamic (Simões-Costa et al., 2014; Martik et al., 2017; Soldatov et al., 2019; Williams et al., 2019). In support of the model that complex transcriptional programs govern NCC ontogenesis, gene regulatory networks involved in early development of NCCs into broad cell types has been studied at a high level using a combination of transcriptomics, chromatin profiling and enhancer studies, especially during pre-migratory and early migratory NCC specification along cranial axial regions, across amniotes (Martik et al., 2017; Simoes-Costa and Bronner, 2016; Green et al. 2016; Lumb et al., 2017; Williams et al., 2019; Hockman et al., 2019). For example, during pre-migratory stages the transcription factors FoxD3, Tfap2a and Sox9 are important for NCC fate specification and in turn regulate the expression of Sox10, a conserved transcription factor that is expressed along all axial levels by early migrating NCCs and within many differentiating lineages (Sauka-Spengler and Bronner-Fraser, 2008; Martik et al., 2017). Gene regulatory networks that are important for select NCC cell fates, like melanocytes and chondrocytes, have been well characterized (reviewed in Martik and Bronner, 2017). Recently, the regulatory circuitry behind glial, neuronal, and mesenchymal fates of vagal NCC was described (Ling and Spengler, 2019) where *Prrx1* and *Twist1* have been described as key differentiation genes for mesenchymal fate. Despite this progress, however, comprehensive knowledge of the genes that are expressed and participate in NCC lineage differentiation programs during later phases of embryogenesis remains to be fully characterized, particularly for posterior tissues (reviewed in Hutchins et al., 2018). Indeed, altered gene expression during NCC differentiation can cause several neurocristopathies, such as DiGeorge syndrome, neuroblastoma, Hirschsprung disease, Auriculo-condylar syndrome, and Klein-Waardenburg syndrome (Barlow, 1984; Bolande, 1997; Brosens et al., 2016; Escot et al., 2016; Vega-Lopez et al., 2017; Wang et al., 2014), further highlighting the need to understand NCC spatiotemporal gene expression patterns during their differentiation into diverse cellular types.

Previous single-cell transcriptomic studies in zebrafish have laid a strong foundation to globally map early lineages of a majority of cell types through early to middle embryonic development (Wagner et al. 2018; Tambalo et al. 2020), and recently this has been extended into the larval stage (Farnsworth et al., 2020). With respect to the posterior NCC fates, however, many of these cells undergo differentiation programs during the embryonic to larval transition, a developmental stage that emerges between ∼48 hpf to 72 hours post fertilization (hpf). Transcriptomic analysis during this transitional phase therefore would enhance our understanding of the dynamic shifts in cell states that may regulate cellular differentiation programs.

In this study, we leverage the power of single-cell transcriptomics and curate the cellular identities of *sox10*-expressing and *sox10*-derived populations along the posterior zebrafish during development. We have utilized the Tg(*-4.9sox10:EGFP*) (hereafter referred to as *sox10*:GFP) transgenic fish line to identify NCCs and their recent derivatives (Carney et al., 2006). Using *sox10*:GFP^+^ 48-50 hpf embryos and 68-70 hpf larvae, we identified eight major classes of cells: mesenchyme, NCC, neural, neuronal, glial, pigment, muscle, and otic. Among the major cell types, we annotated over 40 cellular subtypes. By leveraging in depth analysis of each time point separately, we remarkably captured the dynamic transition of several NCC fates, most notably we discovered over a dozen mesenchymal subtypes and captured the progressive differentiation of enteric neural progenitors into maturing enteric neurons. Using Hybridization Chain Reaction (HCR) and *in situ* hybridization, we validated the spatiotemporal expression patterns of various subtypes. By computationally merging our 48-50 hpf and 68-70 hpf datasets, we generated a comprehensive atlas of *sox10^+^* cell types spanning the embryonic to larval transition, which can also be used as a tool to identify novel genes and mechanistically test their roles in the developmental progression of posterior NCCs. Using the atlas, we characterized a *hox* signature for each cell type, detecting novel combinatorial expression of *hox* genes within specific cell types. Our intention is that this careful analysis of posterior NCC fates and resulting atlas will aid the cell and developmental biology communities by advancing our fundamental understanding of the diverging transcriptional landscape during the NCC’s extensive cell fate acquisition.

## RESULTS

### Single-cell profiling of sox10:GFP^+^ cells along the posterior zebrafish during the embryonic and larval stage transition

To identify *sox10*-expressing and *sox10*-derived cells along the posterior zebrafish during the embryonic to larval transition, we utilized the transgenic line *sox10:*GFP (**Figure 1A**) (Carney et al., 2006; Kwak et al., 2013). Tissue posterior to the otic vesicle, encompassing the vagal and trunk axial region (**Figure 1B**), was dissected from 100 embryonic zebrafish at 48-50 hpf and 100 larval zebrafishes at 68-70 hpf. Dissected tissues were dissociated and immediately subjected to fluorescence-activated cell sorting (FACS) to isolate *sox10*:GFP^+^ cells (**Figure 1B**; **Figure 1-figure supplement 1A,B**). Isolated cells were then input into 10X Genomics Chromium scRNA-seq assays and captured at a depth of 2300 cells from the 48-50 hpf time point and 2580 cells from the 68-70 hpf time point (**Figure 1C**; **Figure 1-figure supplement 1C**). We performed cell filtering and clustering (**Figure 1-figure supplement 1D-I**) of the scRNA-seq datasets using Seurat (Butler et al., 2018; Stuart et al., 2019) to computationally identify cell populations based on shared transcriptomes, yielding 1608 cells from the 48-50 hpf time point and 2410 cells from the 68-70 hpf time point, totaling 4018 cells for final analysis (**Figure 1-figure supplement 1C**). We detected cell population clusters with transcriptionally unique signatures, as shown in heatmap summaries that revealed the top enriched gene signatures per cluster, with 19 clusters (0-18) from the 48-50 hpf time point (**Figure 1-figure supplement 2A**) and 23 clusters (0-22) from the 68-70 hpf time point (**Figure 1-figure supplement 2B**), totaling 42 clusters across both time points. Datasets were visualized with the t-Distributed Stochastic Neighbor Embedding (tSNE) method, which spatially grouped cells in each cluster, for both time points examined (**Figure 1D,E).** The top significantly enriched markers for each cluster at 48-50 and 68-70 hpf are provided in a table in **Figure 1-source data 1**.

**Figure 1.**
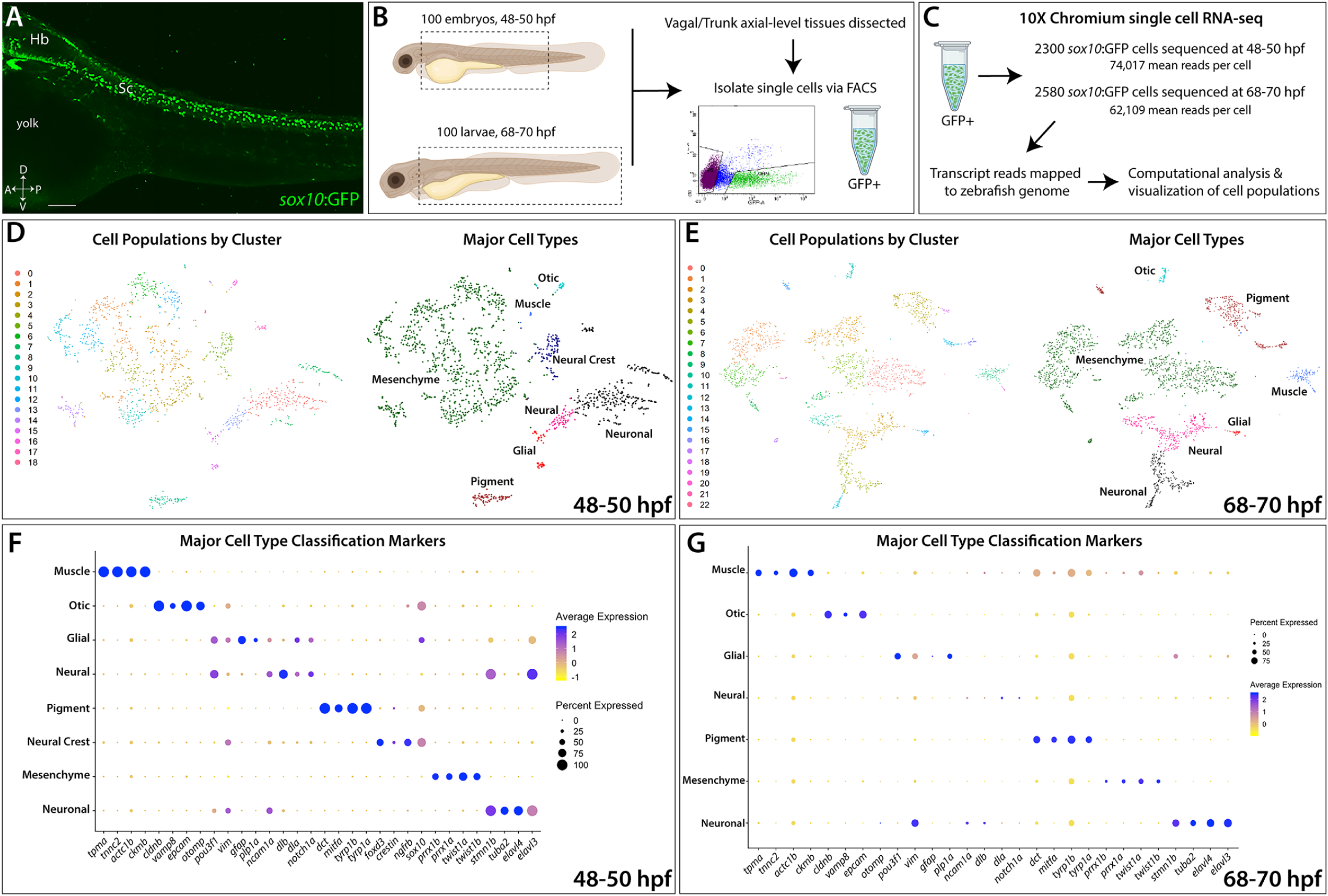
Single-Cell profiling strategy and cell population composition of posterior *sox10*:GFP^+^ cells from the posterior zebrafish during the embryonic to larval stage transition. **(A)** Confocal image of a *sox10*:GFP^+^ embryo at 48 hpf; Hb: Hindbrain; Sc: Spinal cord. A: Anterior, P: Posterior, D: Dorsal, V: Ventral. Scale bar: 50 μM **(B)** Cartoon illustrations of a zebrafish embryo at 48-50 hpf and an early larval fish at 68-70 hpf depicted laterally to summarize the dissection workflow used to collect posterior *sox10*:GFP^+^ cells. **(C)** Schematic of the 10X Genomics Chromium and data analysis pipeline. **(D)** tSNE plots showing the arrangement of Clusters 0-18 and where the major cell types identified among *sox10*:GFP^+^ cells arrange in the 48-50 hpf dataset. **(E)** tSNE plots showing the arrangement of Clusters 0-22 and where the major cell types identified among *sox10*:GFP^+^ cells arrange in the 68-70 hpf dataset. (**F,G**) Dot plots of the identifying gene markers for each major cell type classification in the 48-50 hpf and 68-70 hpf datasets, respectively. Dot size depicts the cell percentage for each marker within the dataset and the color summarizes the average expression levels for each gene.

### Major classification of sox10:GFP^+^ cell states

To assess the proliferative state of *sox10:*GFP***^+^*** cells, we determined their G1, S or G2/M phase occupancy, based on expression of proliferative cell cycle marker genes (**Figure 1-figure supplement 3I**). At 48-50 hpf, 52% of *sox10*:GFP^+^ cells were in G1 phase, 31% were in the S phase and 17% in G2/M phase (**Figure 1-figure supplement 3G**), collectively indicating that 48% of the cells in the 48-50 hpf time point were proliferative. At 68-70 hpf, 64% of cells were in G1 phase, 24% of cells were in the S phase and 12% in G2/M phase (**Figure 1-figure supplement 3G**), indicating that 36% of the cells were proliferative. The cell cycle occupancy distributions were visualized in tSNE plots, revealing congregations of proliferative and non-proliferative *sox10*:GFP^+^ cells (**Figure 1-figure supplement 3A,B**); *aurkb* and *mcm3* confirmed general occupancy in the G2/M and S phase (**Figure 1-figure supplement 3C-F**). Together, these data of cell cycle state reflect a general decrease in proliferative cells among *sox10*:GFP^+^ populations between 48 and 70 hpf, in agreement with prior observations (Rajan et al., 2018).

Using a combination of gene expression searches of the literature and bioinformatics sources, examination of the scRNA-seq transcriptomes indicated that *sox10*:GFP^+^ cells exist in several major cell type categories. Major cell type categories were based on the presence of signature marker genes (**Figure 1F,G; Figure 1-figure supplement 2E,F**). These major cell type categories included: neural, neuronal, glial, mesenchyme, pigment cell, NCC, otic, and muscle; their respective fraction of the datasets were also calculated (**Figure 1D-G; Figure 1-figure supplement 2C,D; Figure 1-figure supplement 3H**). Neuronal refers to cells predominantly expressing neuron markers, such as *elavl3/4*, while neural cells are defined by a multipotent state with potential towards fates of either glial or neuron identity, and marked by expression of factors such as *sox10*, *dla* and/or *ncam1a*.

Notably, mesenchyme tissue identity among clusters represented the largest proportion of the datasets at 61% and 53% of the cells at 48-50 and 68-70 hpf, respectively (**Figure 1-figure supplement 3H**). Mesenchyme clusters were identified by a combination of mesenchymal gene markers, including: *snai1a/b*, *twist1a/b*, *prrx1a/b*, *meox1*, *foxc1a/b*, *cdh11, sparc, colec12*, and/or *pdgfra*, as recently described in amniotes (Soldatov et al., 2019). Pigment cell development markers *tyrp1a/b*, *dct*, and *mitfa* were expressed in 5% of the 48-50 hpf dataset and increased to 14% at 68-70 hpf (**Figure 1-figure supplement 3H**), during which time the pigment cells diverged and expanded into distinct chromatophore lineages, as discussed further in **Figure 2**. A NCC cluster, Cluster 5 at 48-50 hpf, which we also discuss further in **Figure 4**, was identified by the combined expression of *sox10*, *crestin*, *foxd3, ngfrb*, and *tfap2a*. Cells with an otic vesicle and muscle signature were also detected (**Figure 1D-G**; **Figure 1-figure supplement 3H**; **Figure 1-figure supplement 5**), as has previously been described in the *sox10*:GFP line (Carney et al., 2006; Rajan et al., 2018; Rodrigues et al., 2012; Kwak et al., 2013). Overall, major cell type cluster identities and top signature marker genes are summarized in a table in **Figure 1-figure supplement 4**.

**Figure 2.**
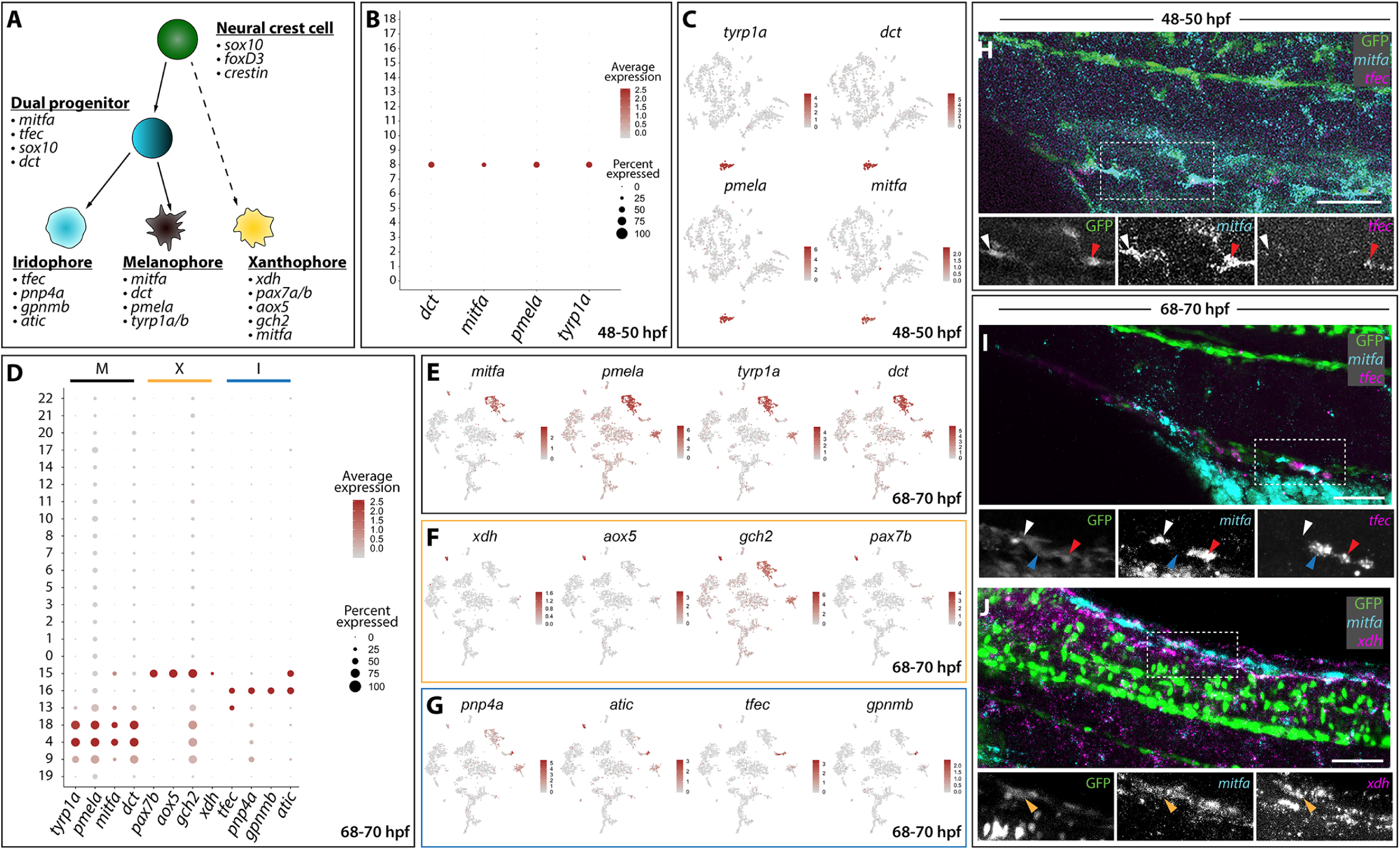
Distinct pigment cell populations are present among *sox10*:GFP^+^ cells during embryonic to larval transition. **(A)** Cartoon schematic depicting the model for neural crest delineation into pigment cell lineages and the genes that were used to identify each pigment cell population. **(B)** Dot plot identifying melanophore markers within the 48-50 hpf dataset. Dot size depicts the cell percentage for each marker within the dataset and the color summarizes the average expression levels for each gene. **(C)** tSNE plots depicting melanophore signature in the 48-50 hpf dataset. Relative expression levels are summarized within the color keys, where color intensity is proportional to expression level of each gene depicted. **(D)** Dot plot showing distinct pigment chromatophore markers within the 68-70 hpf dataset. Dot size depicts the cell percentage for each marker within the dataset and the color summarizes the average expression levels for each gene. M: melanophore markers; X: xanthophore markers; I: iridophore markers. **(E-G)** tSNE plots revealing the location of melanophores **(E)**, xanthophores **(F)**, and iridophores **(G)** in the 68-70 hpf dataset. Relative expression levels are summarized within the color keys, where color intensity is proportional to expression level of each gene depicted. **(H)** HCR against *mitfa* and *tfec* at 48-50 hpf reveals *mitfa*^+^ melanophores (white arrowhead) and *mitfa*^+^/*tfec*^+^ pigment progenitors (red arrowhead). Cropped panels show individual fluorescent channels. **(I)** HCR against *mitfa* and *tfec* at 68-70 hpf presents *mitfa*^+^ melanophores (white arrowhead), *tfec*^+^ iridophores (blue arrowhead), and *mitfa*^+^/*tfec*^+^ pigment progenitors (red arrowhead). Cropped panels show individual fluorescent channels. **(J)** HCR against *mitfa* and *xdh* at 68-70 hpf shows *mitfa*^+^/*xdh*^+^ xanthophores (orange arrowhead). Cropped panels show individual fluorescent channels. Scale bar in H-J: 50 µm.

### Annotation of posterior sox10:GFP^+^ cell subtypes

Analysis of the 42 cluster gene signatures among the two time points revealed distinct subpopulations of posterior *sox10*:GFP^+^ cells and their transcriptomes in the developing zebrafish. Indeed, we identified previously described *sox10*-derived cell subtypes. The *sox10*:GFP line has been shown to transiently label sensory dorsal root ganglion (DRG) progenitors between the 1^st^ and 2^nd^ day of development (McGraw et al., 2008; Rajan et al., 2018). We observed sensory neuronal/DRG gene expression in Cluster 17 at 48-50 hpf (**Figure 1-figure supplement 4**; **Figure1-figure supplement 6**) by the markers *neurod1*, *neurod4, neurog1, six1a*/*b*, *elavl4* (Carney et al 2006; Delfino-Machín et al., 2017). In addition, the scRNA-seq transcriptome datasets at both time points exhibited gene expression indicative of previously described NCC-derived lineages (summarized in Hutchins et al., 2018) including mesenchymal cells (Le Lievre and Le Douarin, 1975; Kague et al., 2012; Soldatov et al., 2019; Ling and Sauka-Spengler, 2019), pigment cells (Reedy et al., 1998; Higdon et al., 2013), and enteric neurons (Kelsh and Eisen, 2000; Kuo and Erickson, 2011; Lasrado et al., 2017), which we describe in further detail for both time points in Figures 2-5.

### Identification of pigment cell chromatophore lineages

With robust genetic lineage details published on pigment cell differentiation in zebrafish (Kelsh, 2004; Lister, 2002; Quigley and Parichy, 2002), we sought to validate our scRNA-seq analysis by assessing if we could resolve pigment cell gene expression states. Pigment cell development has been broadly studied in the developing zebrafish, where NCCs give rise to three distinct chromatophore populations: melanophores, xanthophores, and iridophores (**Figure 2A**). Melanophores, the best characterized population, express a combination of markers throughout their development including *mitfa*, *dct*, *tyrp1a/b*, and *pmela* (Du et al., 2003; Lister et al., 1999; Ludwig et al., 2004; Quigley and Parichy, 2002). Similarly, the genes *pnp4a*, *tfec*, *gpnmb*, and *atic* are all enriched in iridophores and are critical for their maturation (Higdon et al., 2013; Lister et al., 2011; Petratou et al., 2018, 2019). Finally, differentiating xanthophores express *gch2*, *pax7a/b*, *xdh*, *mitfa*, and *aox5* (Nord et al., 2016; Parichy et al., 2000; Saunders et al., 2019; Minchin and Hughes, 2008; Lister et al., 1999).

Our cluster analysis of *sox10*:GFP^+^ single cell datasets revealed the robust presence of pigment cell lineages (**Figure 2B-J**; **Figure 1-figure supplement 2**). At 48-50 hpf, melanophores were detected in Cluster 8 based on expression of *mitfa*, *dct*, *tyrp1b*, and *pmela* (**Figure 1F**; **Figure 2A**), reflected by the dot and tSNE plots (**Figure 2B,C**). At 68-70 hpf, we resolved discrete pigment cell populations that included xanthophore, iridophore, and two distinct melanophore clusters (**Figure 2D-G**, **Figure 1-figure supplement 4**). The xanthophores mapped to Cluster 15 and were enriched with *xdh*, *aox5*, *pax7b*, *mitfa*, and *gch2* (**Figure 2A,D,F**). Cluster 16 was identified as iridophores, which presented the well characterized markers: *tfec*, *pnp4a*, *gpnmb*, and *atic* (**Figure 2A,D,G**; **Figure 1-figure supplement 4**).

The use of cell cycle markers revealed that two different melanophore clusters at 68-70 hpf (Clusters 4 and 18) were present in different proliferative states. While the majority of cells in Cluster 4 were in G1, Cluster 18 contained the presence of S and G2/M markers, such as *pcna* and *aurkb*, suggesting that this population is proliferating melanophores (**Figure 1-figure supplement 3B,E,F**; **Figure 2-source data 1**). These data corroborate and extend previous observations where melanophores in distinct proliferative and differentiating states have been described during the larval stage at 5 days post fertilization (dpf) (Saunders et al., 2019). Overall, genes shared between Cluster 4 and Cluster 18, as well as their unique genes, are summarized in **Figure 2-source data 1**.

At 68-70 hpf, we were able to identify a pigment progenitor population, where iridophore and melanophore markers were co-expressed in Cluster 13: an irido-melano progenitor (**Figure 2A,D**). These undifferentiated pigment progenitor cells express *tfec* in combination with *mitfa* and have been described recently at 24, 30, and 48 hpf (Petratou et al. 2018). Additionally, Cluster 13 expressed *tfap2e*, *gpx3*, and *trpm1b* (**Figure 1-figure supplement 4**) whose expression patterns have been previously reported in pigment progenitors (Saunders et al., 2019). Finally, a population of pigmented muscle (Cluster 9) was also found with a weak melanophore signature coupled with expression of the muscle markers *ckmb*, *tpma*, *tnnc2* and *tnnt3b* (**Figure 1-figure supplement 4**; **Figure 1-figure supplement 5**).

We next performed whole mount HCR to assess the spatial expression of *mitfa*, *tfec* and *xdh,* and to validate the *sox10*:GFP datasets. When examining *mitfa* and *tfec* at 48-50 hpf (**Figure 2H**), we detected GFP^+^ cells that expressed *mitfa*, identifying the melanophores (**Figure 2H**; white arrowhead), and cells that expressed both *mitfa* and *tfec*, defining the pigment progenitors (**Figure 2H**; red arrowhead). At 68-70 hpf, we confirmed the four distinct pigment populations we identified through Seurat (**Figure 2B-G**): GFP^+^ melanophores expressing *mitfa* only (**Figure 2I**; white arrowhead), iridophores only expressing *tfec* (**Figure 2I**; blue arrowhead), and pigment progenitors expressing both *mitfa* and *tfec* (**Figure 2I**; red arrowhead) were detected. When examining *xdh* and *mitfa* expression patterns, GFP^+^ xanthophores were found to be expressing both markers (**Figure 2J**; orange arrowhead). Taken together, these results show that the scRNA-seq datasets effectively identify discrete subpopulations, and coupled with our HCR analysis, effectively shows we are able to validate these cell populations *in vivo*.

### Mesenchyme in the posterior embryo and larvae exists in various subpopulations

Heatmap analysis of gene expression groups depicted that mesenchyme cells clustered together globally within the datasets (**Figure 1-figure supplement 2C,D; Figure 3A,B**), with *twist1a* expression broadly labeling all mesenchyme cells (**Figure 1-figure supplement 2E,F**). In addition to *twist1a*, mesenchyme cells also expressed *prrx1a/b, twist1b, foxc1a/b*, *snai1a/b*, *cdh11*, *sparc*, *colec12*, *meox1*, *pdgfra* (**Figure 3A,B**), and other known mesenchymal markers such as *mmp2* (**Figure 1-figure supplement 4**) (Janssens et al., 2013; Theodore et al., 2017). In whole mount embryos at 48 hpf, we observed broad expression of *foxc1a* and *mmp2* along the posterior pharyngeal arches and ventral regions of the embryo via *in situ* hybridization (**Figure 3-figure supplement 1C,D**; arrowheads), confirming their expression territories within posterior-ventral mesenchymal tissues.

**Figure 3.**
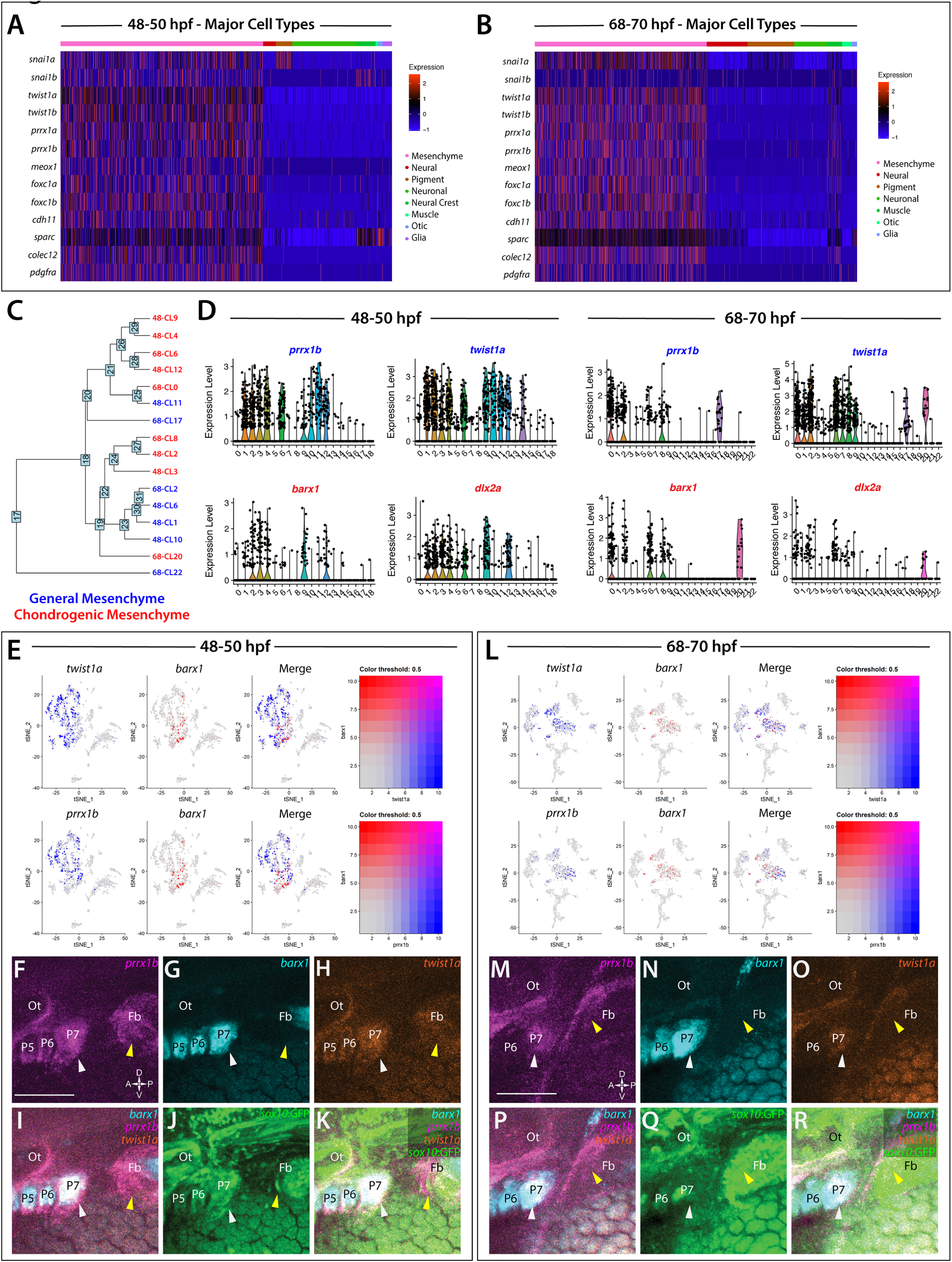
Global analysis of mesenchyme cell signatures among *sox10*:GFP^+^ cells. **(A,B)** A heatmap of signature mesenchyme identity genes within the major cell type classified cells at 48-50 and 68-70 hpf, respectively. Relative expression levels within each cluster is summarized within the color key, where red to blue color indicates high to low gene expression levels. **(C)** A cluster tree depicting the relationship between general and chondrogenic mesenchyme cellular subtypes. **(D)** Violin plots summarizing the expression levels for select mesenchyme identity markers within individual clusters at the 48-50 and 68-70 hpf time points, respectively. Data points depicted in each cluster represent single cells expressing each gene shown. **(E,L)** tSNE plots depicting the expression of *prrx1b*, *barx1* and *twist1a* in the 48-50 and 68-70 hpf datasets, respectively. Relative expression levels are summarized within the color keys, where color intensity is proportional to expression level of each gene depicted. **(F-K)** Whole mount HCR analysis reveals the spatiotemporal expression of *prrx1b* **(F)**, *barx1* **(G)**, *twist1a* **(H)**, *sox:*GFP **(J)** in 48 hpf embryos. **(I)** A merge of *barx1*, *prrx1b* and *twist1a* is shown. **(K)** A merge of *barx1*, *prrx1b*, *twist1a* and *sox10*:GFP is shown. White arrowheads denote expression in posterior pharyngeal arch, while yellow arrowheads highlight fin bud expression. **(M-R)** Whole mount HCR analysis reveals the spatiotemporal expression of *prrx1b* **(M)**, *barx1* **(N)**, *twist1a* **(O)**, *sox:*GFP **(Q)** in 68 hpf embryos. **(P)** A merge of *barx1*, *prrx1b* and *twist1a* is shown. **(R)** A merge of *barx1*, *prrx1b*, *twist1a* and *sox10*:GFP is shown. White arrowheads denote expression in posterior pharyngeal arch, while yellow arrowheads highlight fin bud expression. Ot: otic; Fb: Fin bud. Scale bar: 100 µm.

Analysis of the mesenchyme clusters revealed various subtypes were present in the scRNA-seq datasets. Among these, we detected 9 chondrogenic cell subtypes—identified by expression of mesenchymal signature genes, as well as the chondrogenic markers *barx1* and/or *dlx2a* (Sperber et al., 2008; Sperber and Dawid, 2008) (**Figure 3C,D**). All chondrogenic clusters expressed *barx1*, regardless of the time point, which is expressed in developing mesenchymal tissues and required for development of osteochondral progenitor cells and their tissue condensation therein (Sperber et al., 2008; Sperber and Dawid, 2008; Ding et al., 2013; Barske et al., 2016). Within the 9 chondrogenic subtypes, we discovered genes indicative of heterogeneous cell states; ranging from proliferative, progenitor/stem-like, migratory to differentiating signatures (**Figure 1-figure supplement 4**).

All other mesenchyme subtypes were classified into various progenitor and differentiation categories. Among these categories, 7 clusters expressed either proliferative progenitor markers, differentiation signatures, or general migratory mesenchymal markers (**Figure 1-figure supplement 4**). Additionally, Cluster 14 at 48 hpf and Clusters 1 and 7 at 68-70 hpf exhibited a general mesenchymal signature, but also expressed fin bud marker genes (*hand2, tbx5a, hoxd13a, prrx1a*) (**Figure 1-figure supplement 6**) (Yelon et al., 2000; Lu et al., 2019, Nakamura et al., 2016; Feregrino et al., 2019). The fin bud cells formed distinct groupings, as depicted in tSNE analysis (**Figure 1-figure supplement 6)**.

Visualization of clusters with general mesenchyme and chondrogenic identities using a cluster tree further highlighted potential similar subtypes between the time points (**Figure 3C**). For example, the cluster tree showed proximal location of Cluster 8 at 68-70 hpf and Cluster 2 at 48-50 hpf, which we noted contained clear proliferative chondrogenic gene signatures (**Figure 3D; Figure 1-figure supplement 4**). Further, the tree depicted the closeness of Cluster 1 and 6 at 48-50 hpf with Cluster 2 at 68-70 hpf. These clusters present with varying proliferative/migratory/differentiation states, suggesting that transcriptionally-related mesenchymal cells captured in our datasets were in various stages of dynamic differentiation, proliferation, and migration.

Feature plot exports confirmed that a sub-population of chondrogenic cells (*barx1*^+^) were present in relation to all other mesenchyme (*prrx1b*^+^, *twist1a*^+^) cells in the datasets (**Figure 3E,L**). To confirm the spatial co-expression of *prrx1b*, *twist1a*, and *barx1* at 48-50 hpf and 68-70 hpf, we utilized HCR analysis (**Figure 3F-K,M-R**). The expression of *prrx1b*, *twist1a*, and *barx1* was seen within *sox10*:GFP^+^ cells along the posterior pharyngeal arches (white arrowheads) and fin bud mesenchyme (yellow arrowheads) at both time points (**Figure 3F-K,M-R**). While *prrx1b* and *twist1a* labeled the arches and fin buds (**Figure 3F,H,M,O**), *barx1* was observed in the arches, but not the fin buds (**Figure 3I,G,P,N**).

Overall, these analyses reveal great diversity among the mesenchyme and suggest that mesenchymal cells in the posterior zebrafish exist in various subpopulations and exhibit dynamic transcriptional states during their development. Furthermore, these results show that our dataset analysis can pinpoint previously unknown discrete *sox10* subpopulations.

### sox10-derived cells during the embryonic to early larval transition reveal enteric progenitor to enteric neuron progression

At 48-50 hpf, cells with NCC identity gene signatures were notably detected in Cluster 5, defined by expression of the core markers *sox10, foxd3, crestin*, and *tfap2a* (**Figure 1F; Figure 4A, Figure 1-figure supplement 4**) (Dutton et al., 2001; Luo et al., 2001; Knight et al., 2003; Stewart et. al., 2006). In addition to the core genes, Cluster 5 cells also contained genes previously shown to be expressed in zebrafish NCCs; including *vim, snai1b, sox9b*, *zeb2a, mych*, *and mmp17b* (**Figure 4D**) (Cerdà et al., 1998; Heffer et al., 2017; Hong et al., 2008; Leigh et al., 2013; van Otterloo et al., 2012; Wang et al., 2011; Rocha et al., 2020). We reasoned that many of the NCCs had started their respective differentiation programs and were beginning to assume specified genetic profiles. Therefore, we sought to determine if the NCC cluster also contained gene expression profiles of known differentiating NCC types along the posterior body, such as enteric progenitors, also known as enteric neural crest cells (ENCCs).

**Figure 4.**
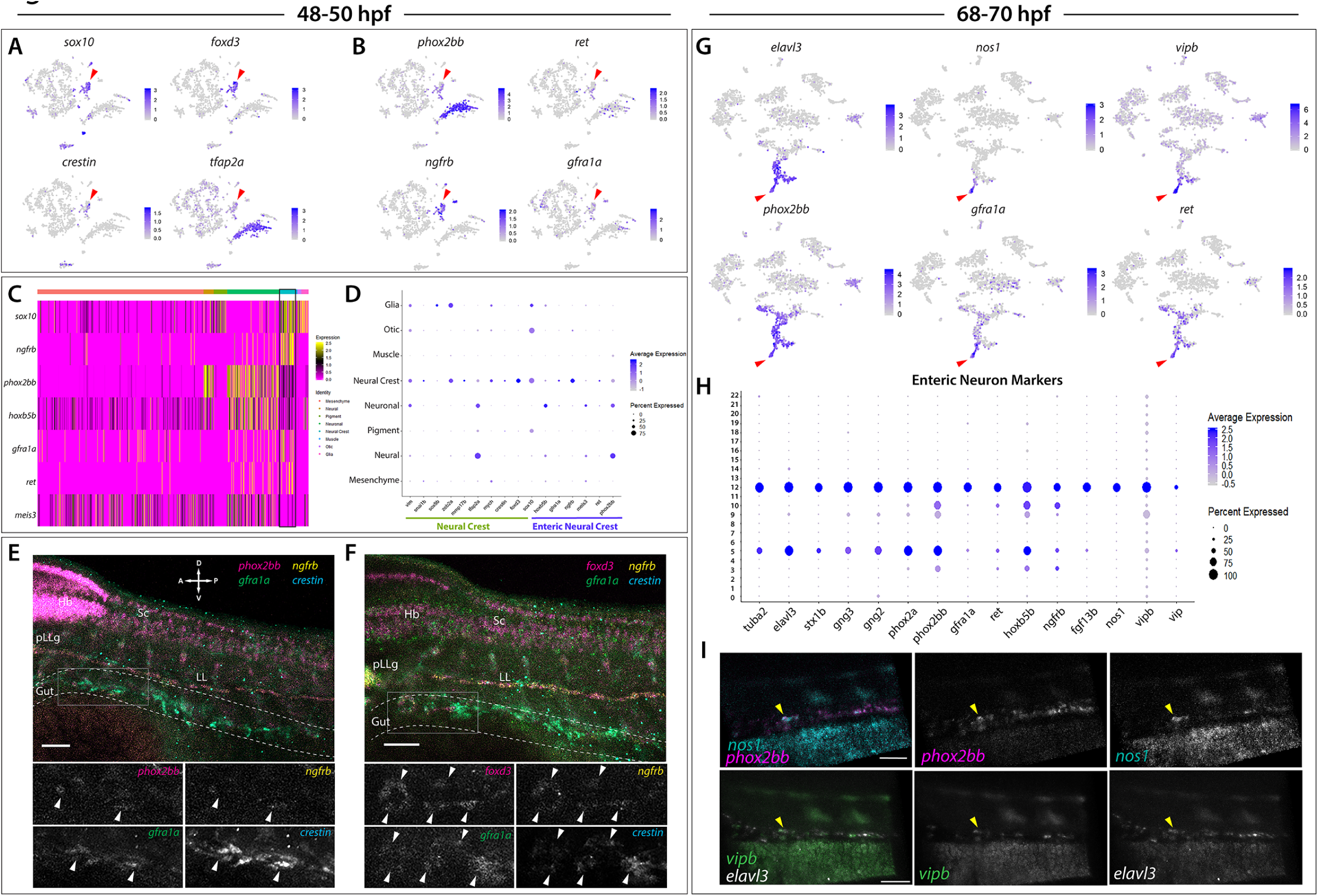
Enteric neural crest cells and differentiating enteric neurons are present among posterior *sox10*:GFP^+^ cell populations. (A) tSNE feature plots reveal expression of core neural crest cell markers *sox10*, *foxd3*, *crestin* and *tfap2a* mapping to the neural crest cell cluster (red arrow). (B) tSNE feature plots depict expression of the enteric neural crest cell markers *phox2bb*, *ret*, *ngfrb* and *gfra1a* within the neural crest cell cluster (red arrow). Relative expression levels are summarized within the color keys in (**A**) and (**B**), where color intensity is proportional to expression level of each gene depicted. (C) A heatmap reveals expression levels of enteric neural crest cell markers across the 8 major cell populations captured in the 48-50 hpf data set (color key denotes cells types represented in color bar on top of heatmap). Neural crest cell cluster highlighted in black rectangle. Relative expression levels within each major cell type cluster is summarized within the color key, where yellow to magenta color indicates high to low gene expression levels. (D) Dot plot of expanded list of neural crest (green line) and enteric neural crest (purple line) cell markers across each major cell type within 48-50 hpf data set. Dot size depicts the cell percentage for each marker within the data set and the color summarizes the average expression levels for each gene. (**E,F**) Whole mount HCR analysis of 48 hpf embryos reveals co-expression of enteric neural crest cell markers within the developing gut (dashed outline). Top panels depict merged images of color channels for each HCR probe. Lower panels represent grey-scale images of each separated channel corresponding to the magnified region of foregut (grey rectangle). Arrows depict regions where all markers are found to be co-expressed. Hb: Hindbrain, Sc: Spinal cord, pLLg: posterior Lateral Line ganglia, LL: Lateral Line. A: Anterior, P: Posterior, D: Dorsal, V: Ventral. Scale bar: 50 μM. (G) tSNE feature plots reveal expression levels of enteric neuron markers *elavl3*, *phox2bb*, *gfra1a*, *nos1*, *vipb* and *ret*, within a common region of a neuronal cluster (red arrow). Relative expression levels are summarized within the color keys, where color intensity is proportional to expression level of each gene depicted. (H) Dot plot depicts expression levels of pan-neuronal and enteric neuron specific markers across individual clusters generated within the original 68-70 hpf tSNE. Pan-neuronal markers found throughout clusters 5 and 12, with enteric neuron markers most prominently expressed within cluster 12. Dot size depicts the cell percentage for each marker within the data set and the color summarizes the average expression levels for each gene. (I) Whole mount HCR analysis depicts differentiating enteric neurons within the foregut region at 69 hpf co-expressing *nos1*, *phox2bb*, *vipb*, and *elavl3* (yellow arrow). Anterior: Left, Posterior: Right. Scale bar: 50 μM.

During NCC diversification, ENCCs fated to give rise to the enteric nervous system (ENS) express a combination of NCC and enteric progenitor marker genes over developmental time (reviewed in Nagy and Goldstein, 2017; Rao and Gershon, 2018), which occurs between 32 to 72 hpf in zebrafish (reviewed in Ganz, 2018). Enteric markers in zebrafish include *sox10*, *phox2bb*, *ret*, *gfra1a*, *meis3*, and *zeb2a* (Dutton et al., 2001; Shepherd et al., 2004; Elworthy et al., 2005; Delalande et al., 2008; Heanue and Pachnis, 2008; Uribe and Bronner, 2015). Given the developmental timing of early ENS formation in zebrafish as occurring between 32 to 72 hpf, we expected to capture a population of ENCCs within our 48-50 hpf dataset. Indeed, within Cluster 5 we observed expression of the enteric markers *phox2bb, ret, gfra1a, meis3, sox10*, and *zeb2a* (**Figure 4B-D**). Using whole mount *in situ* hybridization, we confirmed the expression of *sox10* and *phox2bb* within ENCCs along the foregut at 48 hpf (**Figure 3-figure supplement 1A,A’,B,B’**; arrowheads). Furthermore, gene orthologs known to be expressed in ENCC in amniotes were detected within Cluster 5, such as *ngfrb* (orthologue to p75) (Anderson et al., 2006; Wilson et al., 2004) and *hoxb5b* (orthologous to *Hoxb5*) (Kam and Lui, 2015) (**Figure 4B-D**).

HCR analysis of 48 hpf embryos validated the co-expression profiles of several ENCC markers along the foregut (**Figure 4E-F**; foregut in grey box). Co-expression analysis demonstrated that a chain of *crestin*^+^ cells localized in the foregut contained a subpopulation of cells expressing *ngfrb*, *phox2bb*, and *gfra1a* (**Figure 4E**; white arrowheads), or expressing *foxd3*, *ngfrb*, and *gfra1a* (**Figure 4F**; white arrowheads). Together, these HCR data reveal a subpopulation of ENCCs within the zebrafish gut, confirming ENCC markers are co-expressed.

We next asked if we could resolve discrete differentiating enteric neurons over time. In zebrafish, by 72 hpf ENCCs have migrated throughout the length of the gut and begun early neuron differentiation and neural patterning (Elworthy et al., 2005; Olden et al., 2008; Harrison et al., 2014; Uribe and Bronner, 2015; Taylor et al., 2016). During early neuronal differentiation, ENCCs display differential enteric progenitor gene expression patterns (Taylor et al., 2016) and neurochemical signatures representative of varying stages of neuronal differentiation and subtype diversification (Poon et al., 2003; Holmqvist et al., 2004; Uyttebroek et al., 2010). Zebrafish early differentiating enteric neurons have been characterized by the RNA expression of *sox10*, *phox2bb*, *gfra1a*, *fgf13b*, and *ret,* as well as the immunoreactivity of Elavl3/4 (Shepherd et al., 2004; Heanue and Pachnis, 2008; Uyttebroek et al., 2010; Taylor et al., 2016). In addition, at this time, enteric neurons express multiple neurochemical markers, with Nos1 being most prominent (Olden et al., 2008; Uyttebroek et al., 2010), a finding consistent with studies performed within the amniote ENS (Hao and Young, 2009; Matini et al., 1995; Qu et al., 2008; Heanue et al., 2016). In light of these previous observations, our 68-70 hpf dataset was expected to contain the transcriptomes of ENCCs captured at various stages of their progressive differentiation into the diverse subtypes of the ENS.

tSNE analyses identified differentiating enteric neurons within the 68-70 hpf dataset based off of the combinatorial expression of *elavl3, phox2bb, ret,* and *gfra1a* (**Figure 4G**), which mapped to the neural/neuronal major cell type regions of the dataset (**Figure 1E**), comprising Clusters 5 and 12 (**Figure 1E**). Transcripts that encode for the neurochemical marker *nos1,* and the neuropeptides *vip* and *vipb,* a paralogue to *vip* (Gaudet et al., 2011), were found in a subpopulation of enteric neurons localized to a distal group of the neuronal cluster, likely indicative of a differentiating enteric neuron subtype (**Figure 4G**; red arrows). We then queried for the presence of a combination of pan-neuronal and enteric neuron markers (**Figure 4H**). The pan-neuronal markers *tuba2, elavl3, stx1b,* and *gng2/3,* as well as the autonomic neuron markers, *phox2a* and *phox2bb* (Gou et al., 2018; Hans et al., 2013), were present in both Clusters 5 and 12 (**Figure 4H**). However, the enteric neuron markers, *gfra1a, ret, hoxb5b, ngfrb, fgf13b, nos1, vipb,* and *vip* were mostly confined to Cluster 12, suggesting that this cluster contained differentiating enteric neurons (**Figure 4H**). Indeed, whole mount HCR analysis validated the spatiotemporal expression of *phox2bb, nos1, vipb,* and *elavl3* transcripts throughout the foregut of the zebrafish embryo by 69 hpf (**Figure 4I**; yellow arrowheads). These results suggest that *elavl3*^+^/*phox2bb*^+^ early differentiating enteric neurons in the foregut display an inhibitory neurochemical signature, consistent with prior observations in zebrafish and mammalian ENS (Olden et al., 2008; Hao and Young, 2009).

In an effort to examine the enteric neuron populations with finer resolution, Clusters 5 and 12 were subset from the main dataset in Seurat, re-clustered and visualized using a tSNE plot, producing 5 new clusters (**Figure 5A**). The gene markers from each new Sub-Cluster are provided in **Figure 5-source data 1**. Following this, the previously mentioned enteric neuron markers, with the addition of *etv1,* a recently identified marker of enteric intrinsic primary afferent neurons (IPANs) in mouse (Morarach et al., 2020), were queried and visualized using dot and feature plots allowing the identification of Sub-Cluster 3 as a differentiated enteric neuron cluster (**Figure 5A-C**). *nos1*, *vipb* and *vipb* were enriched in Sub-Cluster 3 (**Figure 5B,C; Figure 5-figure supplement 1**). Interestingly, while expressed at lower average levels than in Sub-Cluster 3, the enteric combination markers were also present in Sub-Cluster 1 (**Figure 5B-C**). Sub-Cluster 1 formed a central point from which Sub-Cluster 3 could be seen emanating as a distal population (**Figure 5A**). Given the developmental timing, this is likely depicting enteric neurons captured at different stages along their progressive differentiation, which would suggest that the distal most population represents the more mature enteric neurons.

**Figure 5.**
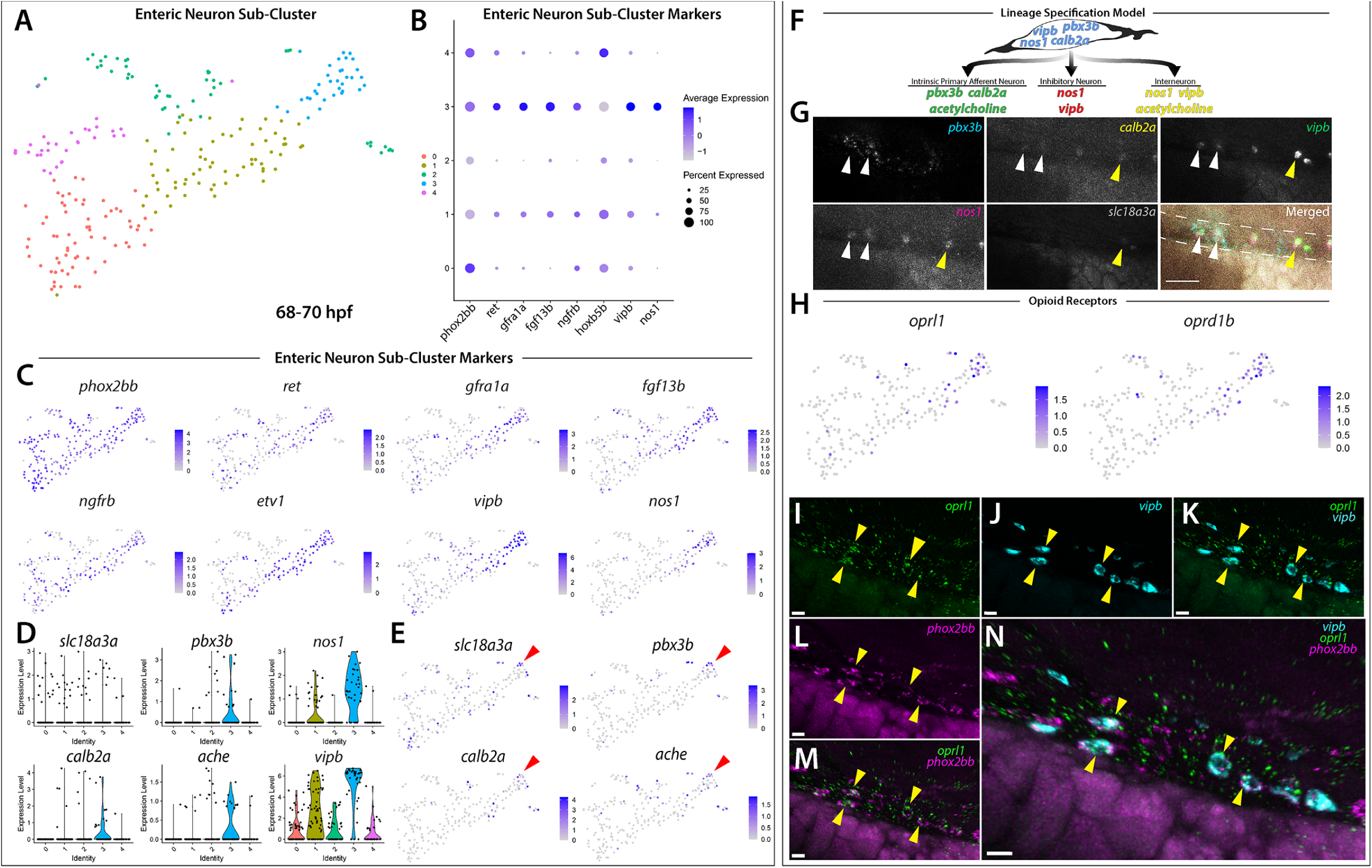
Differentiating enteric neurons captured during key transitional stage of subtype diversification within 68-70 hpf *sox10*:GFP^+^ larval cells. **(A)** tSNE plot reveals 5 distinct sub-clusters following the subset analysis and re-clustering of clusters 5 and 12 from the 68-70 hpf data set. **(B)** Dot plot depicts expression levels of enteric neuron markers across resulting Sub-clusters. Each marker was expressed at low levels in Sub-cluster 1 and were found to be expressed at higher levels within Sub-cluster 3. **(C)** tSNE feature plots further depict the expression of enteric neuron markers by illustrating the levels and localization of expression within the Sub-cluster architecture. Feature plots supplement dot plot and demonstrate the prominent expression of enteric neuron markers within Sub-cluster 3, which appears to emanate from Sub-cluster 1. **(D,E)** Violin and feature plots reveal expression levels of acetylcholine-associated and excitatory neuron markers reported to distinguish enteric IPANs. These markers were found in a discrete pocket of cells forming the distal-most region of Sub-cluster 3 (red arrow). Violin data points depicted in each Sub-cluster represent single cells expressing each gene shown. **(F)** Graphical model summarizes expression patterns observed in 68-70 hpf data set and HCR validation. Common enteric neuroblast capable of diverging into subsequent lineages, IPAN, inhibitory neuron, and interneuron through lineage restricted gene expression. *pbx3b* promotes assumption of IPAN role through loss of *nos1* and *vipb* and begins expressing *calb2a*, *ache* and *slc18a3a*. **(G)** Whole mount HCR analysis reveals co-expression of IPAN markers, *pbx3b* and *calb2a*, and inhibitory neurochemical markers, *vipb* and *nos1* (white arrows), within the foregut (dashed white line) at 68 hpf. Vesicular acetylcholine transferase, *slc18a3a,* was not observed in tandem with *pbx3b* but was co-expressed with *calb2a, vipb,* and *nos1* (yellow arrow). Scale bar: 50 μM. **(H)** Feature plots reveal expression of opioid receptors, *oprl1* and *oprd1b* within the differentiated enteric neuron Sub-Cluster 3. **(I-N)** Whole mount HCR analysis validates expression of *oprl1* in combination with vipb and *phox2bb* (yellow arrows) in enteric neurons localized to the foregut region of a 68 hpf embryo. Scale bar: 10 μM.

Given our hypothesis that the enteric neurons further along a differentiation program were localized to the distal tip of Sub-Cluster 3, we asked whether this population of cells contained additional neurochemical or neuron subtype specific differentiation genes. As expected, a small pocket of these cells was found to contain the two acetylcholine associated genes, *acetylcholine esterase* (*ache)* (Bertrand et al., 2001; Huang et al., 2019) and *vesicular acetylcholine transferase* (*slc18a3a*) (Hong et al., 2013; Zoli, 2000) (**Figure 5E**; red arrowheads). Within Sub-Cluster 3, we detected the expression of *calb2a* and *pbx3b* (**Figure 5D-E; Figure 5-figure supplement 1**), two genes that have previously been shown to denote myenteric IPANs in mammals (Furness et al., 2004; Memic et al., 2018). Corroborating our single-cell findings, HCR analysis revealed the co-expression of *pbx3b, calb2a, vipb, nos1,* and *slc18a3a* in discrete differentiating enteric neurons within the foregut region of the zebrafish gut at 68 hpf (**Figure 5G**). In particular, *calb2a, vipb, nos1,* and *slc18a3a* were all found to be co-expressed (**Figure 5G**; yellow arrowheads). While *pbx3b* expression was found in combination with *calb2a, vipb, and nos1* (**Figure 5G**; white arrowheads), we were unable to observe detectable levels of *slc18a3a* within the *pbx3b* expressing cells. Taken together, these results regarding enteric neuron subpopulation gene expression patterns suggest a model (**Figure 5F**), whereby *nos1, vipb, calb2a,* and *pbx3b* are co-expressed within early enteric neurons, and that through lineage-restricted gene expression, *pbx3b* expression may promote the assumption of an IPAN signature characterized by the presence of *calb2a, ache,* and *slc18a3a* and the loss of inhibitor markers *nos1* and *vipb*. Therefore, our single-cell analysis in zebrafish suggests that the transcriptional emergence of specific enteric neuron subtypes may be conserved between vertebrate species.

In order to identify the presence of novel gene pathways within the developing enteric neuron population, the significantly enriched gene list from Sub-Cluster 3 was processed using gene ontology (GO) pathway enrichment analysis. We found that three opioid signaling pathways were among the top 10 highest fold enriched pathways (**Figure 5-source data 2**) (Mi et al., 2019). Further investigation into these pathways led to the identification of G-protein-coupled receptors, *oprl1* and *oprd1b,* respectively representing nociception/orphan FQ (N/OFQ) peptide (NOP)-receptor and delta-opioid receptor (DOR) subtype members within the opioid receptor superfamily (**Figure 5H**) (Donica et al., 2013; Sobczak et al., 2014). Specifically, feature plots showed that the expression of the opioid receptors was tightly confined within the pocket of enteric neuron progenitors we identified as undergoing sensory lineage-specification, suggesting the expression of opioid receptor genes within both excitatory and inhibitory neurons at the early stages of enteric neuron differentiation within the zebrafish ENS (**Figure 5E and H**). Confirming this suggestion using HCR analysis, we observed the combinatorial expression of *oprl1*, *vipb* and *phox2bb* within migrating enteric neuron progenitors at 68 hpf along the foregut (**Figure 5I-N**). The presence of opioid receptors within immature enteric neurons undergoing lineage-specification helps us to better understand the complexity of early ENS signaling and highlights an area that requires further investigation.

Based on our observation that the enteric neuron population that comprised Sub-Cluster 3 only made up one of five *phox2bb^+^* Sub-Clusters (**Figure 5A-C**), we suspected that the remaining Sub-Clusters were made up of closely related autonomic neurons. In order to better visualize specific differences between the Sub-Clusters, we viewed them using UMAP (Becht et al., 2019; Mcinnes et al., 2018) (**Figure 5-figure supplement 2**). While the identity of the previous tSNE Sub-Clusters were maintained, UMAP analysis allowed us to better visualize the separation between the Sub-Clusters, which we were able to broadly classify as autonomic neurons based on their shared expression of *ascl1a, hand2, phox2a* and *phox2bb* (**Figure 5-figure supplement 2A,B,D**). Within the population of autonomic neurons, we were able to distinguish a population of sympathetic neurons within Sub-Cluster 2 based on their combinatorial expression of *th, dbh, lmo1* and *insm1a* (**Figure 5-figure supplement 2B,E**), which could be clearly distinguished from the enteric neuron Sub-Cluster 3 (**Figure 5-figure supplement 2B,F**). Using a cluster tree, we were able to confirm the distinction between enteric neuron Sub-Cluster 3 and sympathetic neuron Sub-Cluster 2 (**Figure 5-figure supplement 2C**). Overall, these results suggest that Sub-Cluster 0 cells may represent a pool of immature sympatho-enteric neurons, and that Sub-Clusters 1 and 4 both represent better resolved, yet still immature, pools of enteric and sympathetic neurons.

### Atlas of sox10:GFP^+^ cell types encompassing the embryonic-to-larval transition

To describe the dynamic relationship between *sox10*:GFP^+^ cells across both time points, we merged the 48-50 hpf and 68-70 hpf datasets using Seurat’s dataset and Integration and Label Transfer utility (Stuart et al., 2019). The merged datasets were visualized via UMAP, which allowed us to describe the transition between cell types by making both inter- and intra-cluster comparisons. From the merged datasets, we detected 27 clusters (**Figure 6-figure supplement 1A**). We observed that every cluster identified in the 48-50 hpf dataset mapped proximally to clusters at 68-70 hpf (**Figure 6-figure supplement 2A**). We labeled each cell using the major cell type categories: neural, neuronal, glial, NCC, pigment, mesenchyme, otic, and muscle, forming a major cell type atlas (**Figure 6-figure supplement 1B**). Further refinement of the cell identities based on our previous curation (**Figure 1-figure supplement 2**) allowed us to form a higher resolution atlas for each cell type (**Figure 6A**). The genes for each major cell type category in the atlas is provided in **Figure 6-source data 1**.

**Figure 6.**
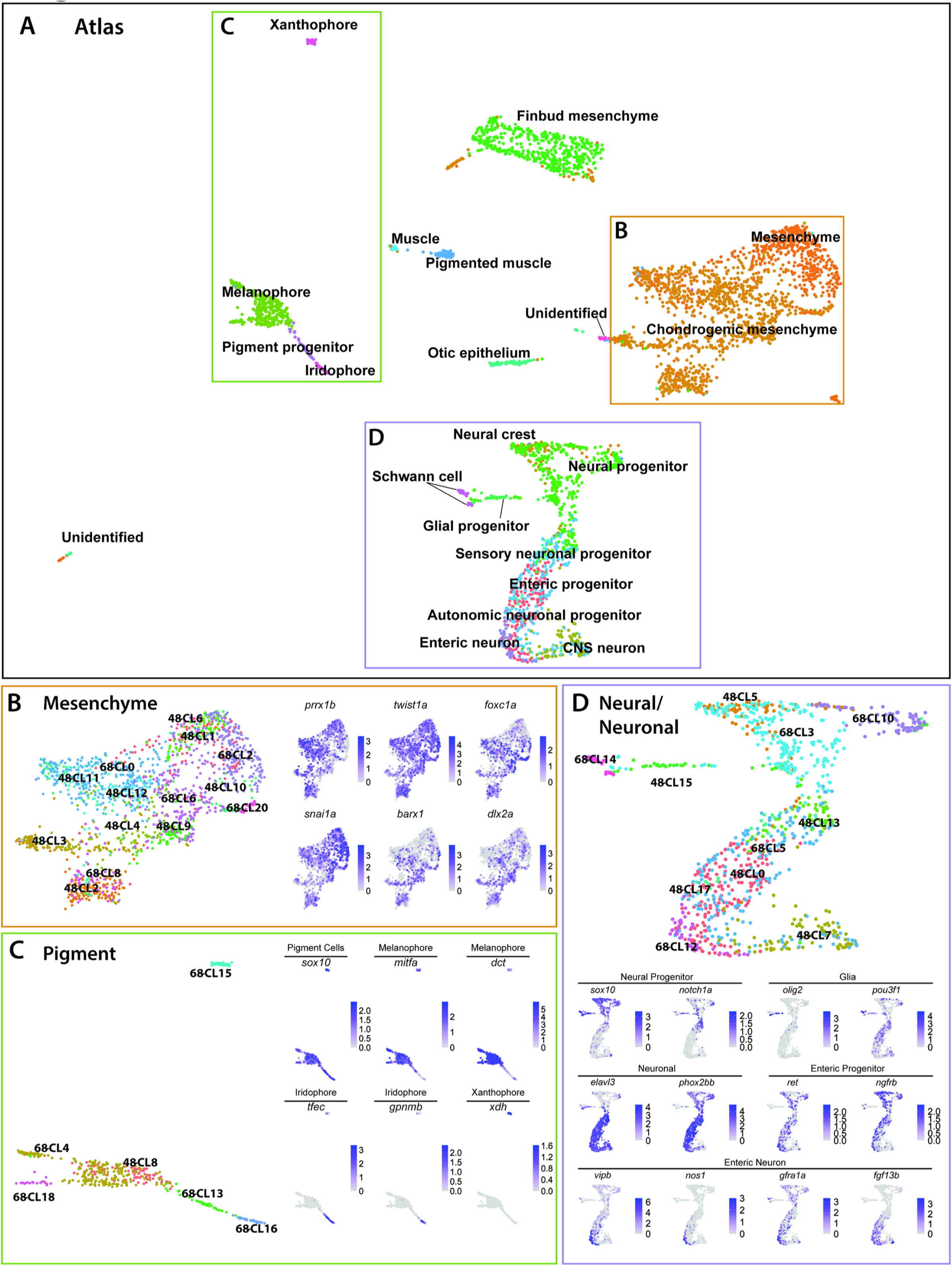
Integrated atlas of posterior *sox10*:GFP^+^ cell types spanning the embryonic to larval transition. **(A)** Global UMAP embedding demonstrating the clustering of cell types across 48-50 hpf and 68-70 hpf. Cell labels were transferred from the original curation to the new atlas after its creation, allowing for unbiased assessment of cell type organization. **(B)** Previously identified mesenchyme clusters form a large discernible cluster marked by *prrx1b*, *twist1a*, *foxc1a*, and *snai1a*, which was separated into both chondrogenic and general mesenchyme, as denoted by its differential expression of *barx1* and *dlx2a*. Importantly, nearly every 48-50 hpf cell type nests with a cluster at 68-70 hpf. **(C)** Pigment cells clusters reflect differentiation paths described in Figure 4A. Melanophores at 48-50 hpf group near to the 68-70 hpf melanophore cluster, bipotent pigment progenitors bridges both the iridophores and melanophores. Xanthophores cluster separately, reflecting their distinct lineage of origin at this developmental window. **(D)** Detailed analysis of the larger neural/neuronal cluster shows clear progression of cell fates from progenitor to differentiating glia or neuron. We confirm the presence of a clear enteric neuronal population, which is distinct from other subtypes at this dataset.

The overall architecture of the UMAP revealed that the cells of each major cell type congregated together, strongly supporting that our formerly described characterizations of each time point are accurate and predictive. For example, cells with mesenchyme identity constellated together, forming three large mesenchyme regions in the atlas: general, chondrogenic, and fin bud (**Figure 6A,B**). NCCs mapped precisely to the top end of a neural/neuronal region of the atlas, while differentiating neurons positioned at the base (**Figure 6A,D**). Additionally, pigment cell types aligned adjacent to one another in the atlas (**Figure 6A,C**).

Comparison of individual subsets of clusters provided deeper analysis of each cell type, with respect to changing cell states across time. We examined three of the largest regions of the atlas in detail: mesenchyme, pigment, and neural/neuronal cell types. We observed consistency across the larger mesenchyme population, which excluded the fin bud mesenchyme, in both the cluster identity shown in the UMAP and gene expression profiles. For example, 48h-Cluster 2 and 68h-Cluster 8 both showed a very high degree of similarity, as well as consistent *barx1, dlx2a*, and *twist1a* expression (**Figure 6B**), consistent with our prior analysis (**Figure 3C,D**). The central node of the pigment region within the atlas was marked by the 48h-Cluster 8, which resolved into respective pigment chromatophore clusters at 68-70 hpf (**Figure 6C**). We were able to globally discern that pigment cell types displayed expression of *sox10*, regardless of time point. As expected from our previous analysis, we observed the early specified melanophore population at 48h-Cluster 8 branched into later stage melanophore populations (68h-Clusters 4 and 18), both of which expressed *mitfa* and *dct*. Further, we observed that the common bi-potent pigment progenitor population (68h-Cluster 13) bridged both melanophore clusters and the iridophore 68h-Cluster 16. This nested positioning of 68h-Cluster 13 supports its dual progenitor identity. Xanthophores, marked by *xdh*, segregated tightly away from the remaining pigment populations, reflective of their earlier and distinct lineage (**Figure 2A**).

Cells within the neural/neuronal clusters of the atlas assembled such that progenitor cells bridged into differentiating neurons spatially from the top to the bottom of the neural/neuronal region of the atlas (**Figure 6D**). 6 clusters (Clusters 0, 5, 7, 13, 15, 17) were represented from 48-50 hpf, and 5 clusters (Clusters 3, 5, 10, 12, 14) from 68-70 hpf. The 68-70 hpf neural progenitor populations (Clusters 3 and 10) shared common gene expression with the 48-50 hpf NCC population (48h-Cluster 5), reflected largely by their co-expression of *sox10*, *notch1a*, *dla*, and *foxd3* (**Figure 6D; Figure 6-figure supplement 1D**). We confirmed the spatiotemporal expression domains of *notch1a* and *dla* along the hindbrain, spinal cord, and in NCC populations along the post-otic vagal domain at 48 hpf (**Figure 3-figure supplement 1E,F**; arrowheads), in particular with *dla* in the ENCCs along the foregut (**Figure 3-figure supplement 1F**; arrow), a pattern similar to the ENCC makers *sox10* and *phox2bb* (**Figure 3-figure supplement 1A,B**). Delineated from the neural progenitor cells, we observed a bifurcation in cell states; with one moving towards a Schwann/glial cell fate, while the other branched towards neuronal. The glial arm followed a temporal progression of earlier cell fates at 48-50 hpf (48h-Cluster 15) toward the more mature fates at 68-70 hpf (68h-Cluster 14), both denoted by expression of *olig2* and *pou3f1*, respectively (**Figure 6D**). From 48h-Cluster 13, we observed the beginning of the neuronal populations, namely 48h-Clusters 0, 7, 13, and 17 and 68h-Clusters 5 and 12. Cells in these clusters patterned in the atlas UMAP such that the progenitor clusters (48h-Clusters 0, 13, and 17; 68h-Cluster 5) formed a spectrum of cell states leading toward the neuronal populations (48h-Cluster 7; 68h-Cluster 12). Among the neuronal populations, we observed a clear autonomic signature, indicated by *phox2a* and *phox2bb* (**Figure 6D; Figure 6-figure supplement 1D**). We also detected a large fraction of enteric progenitors, indicated by *ret, ngfrb,* and *hoxb5b* expression, (**Figure 6D; Figure 6-figure supplement 1D**) supporting our previous observations (**Figure 4,5**). The enteric progenitors culminated into a pool of enteric neurons, with the specific neural signature: *vipb, nos1, gfra1a, fgf13b,* and *etv1* (**Figure 6D; Figure 6-figure supplement 1D**).

Closer inspection of the pigment, mesenchymal, and neural/neuronal clusters separated by time highlighted both predicted and novel changes in gene expression patterns (**Figure 6 - figure supplement 2**). Analysis of each of these sub-setted cell type lineages revealed temporal differences between the two stages, as clearly exemplified in the pigment subcluster (**Figure 6 - figure supplement 2B**). For example, the xanthophore differentiation marker *xdh* demonstrated restricted expression to 68-70hpf. Further, we identified genes with no known roles in pigment cell development differentially expressed between the two sages, such as *rgs16* and *SMIM18*. Within the mesenchyme lineages, both *barx1* and *snai1b* followed expected temporal expression trends, while *abracl and id1* both demonstrated novel differential gene expression profiles within the mesenchyme (**Figure 6 - figure supplement 2C**). Lastly, the neural/neuronal lineage showed expected differential gene expression of genes such as *etv1* and *vipb*, while revealing novel expression of *nova2* and *zgc:162730,* which have previously uncharacterized roles in *sox10*-derived cells (**Figure 6 - figure supplement 2D**). Together, these findings demonstrate the power of the *sox10* atlas not only at cataloguing lineages, but also at characterizing novel gene expression changes across developmental stages.

### A hox gene signature within sox10-derived cells in the posterior zebrafish

A common theme examined by many recent and insightful single cell profile studies of the NCC (Dash and Trainor, 2020; Soldatov et al., 2019) is that patterns of Homeobox transcription factors, known as *hox* genes, display discrete expression between various cell lineages, such as in the cranial NCC. We wondered whether specific *hox* signatures were expressed within posterior NCC and their recent derivatives. To analyze if we could detect *hox* gene patterns within the atlas, we queried all the known canonical *hox* genes within zebrafish as listed on zfin.org (Ruzicka et al., 2019). We detected broad expression of 45 of the 49 zebrafish *hox* genes across the atlas, with 85% of the cells in the atlas expressing at least one *hox* gene (**Figure 7-figure supplement 1A,J**). The four undetected hox genes (*hoxc1a*, *hoxc12b*, *hoxa11a*, and *hoxa3a*) were not examined further.

A dot plot revealed that specific *hox* gene expression patterns demarcated distinct tissues, with specific robustness in the neural fated cells (**Figure 7A)**. Common to the neural lineages, we observed a core *hox* profile which included *hoxb1b, hoxc1a, hoxb2a, hoxb3a, hoxc3a, hoxd3a, hoxa4a*, *hoxd4a, hoxb5a, hoxb5b, hoxc5a, hoxb6a, hoxb6b,* and *hoxb8a* (**Figure 7A**). One of the top expressed constituents of the core signature, *hoxa4a,* was also widely expressed in several other lineages (**Figure 7A; Figure 7 - figure supplement 1O**). The *hox* signature applied to the NCC, neural progenitor, enteric progenitor, enteric neuron, glial progenitor, autonomic neuronal progenitor, and CNS lineages described in the atlas. Clustering of all the atlas lineages relying only on *hox* gene expression highlighted the robustness of the core *hox* signature to distinguish the neural lineage fates, grouping the differentiating (autonomic neuronal progenitors, enteric neurons, and enteric progenitors, and CNS neurons) and progenitor (neural progenitors and glial progenitors) lineages into neighboring clades (**Figure 7B**). Building on the core neural signature unifying the neural fates, slight variations in *hox* expression between autonomic and enteric lineages distinguished them from one another, which are summarized in **Figure 7E.** Most notably, considering the lineages in increasing specificity of cell fate, there was a detectable shift in *hox* expression among the autonomic neural progenitors to the enteric neuronal lineage, demarcated by the increase in *hoxb2a, hoxd4a, hoxa5a, hoxb5a,* and *hoxb5b,* accompanied by diminished expression of *hoxc3a, hoxc5a, hoxb6a* and *hoxb8a*, which formed a distinctive enteric hox signature (**Figure 7 - figure supplement 1 A, K-N**).

**Figure 7.**
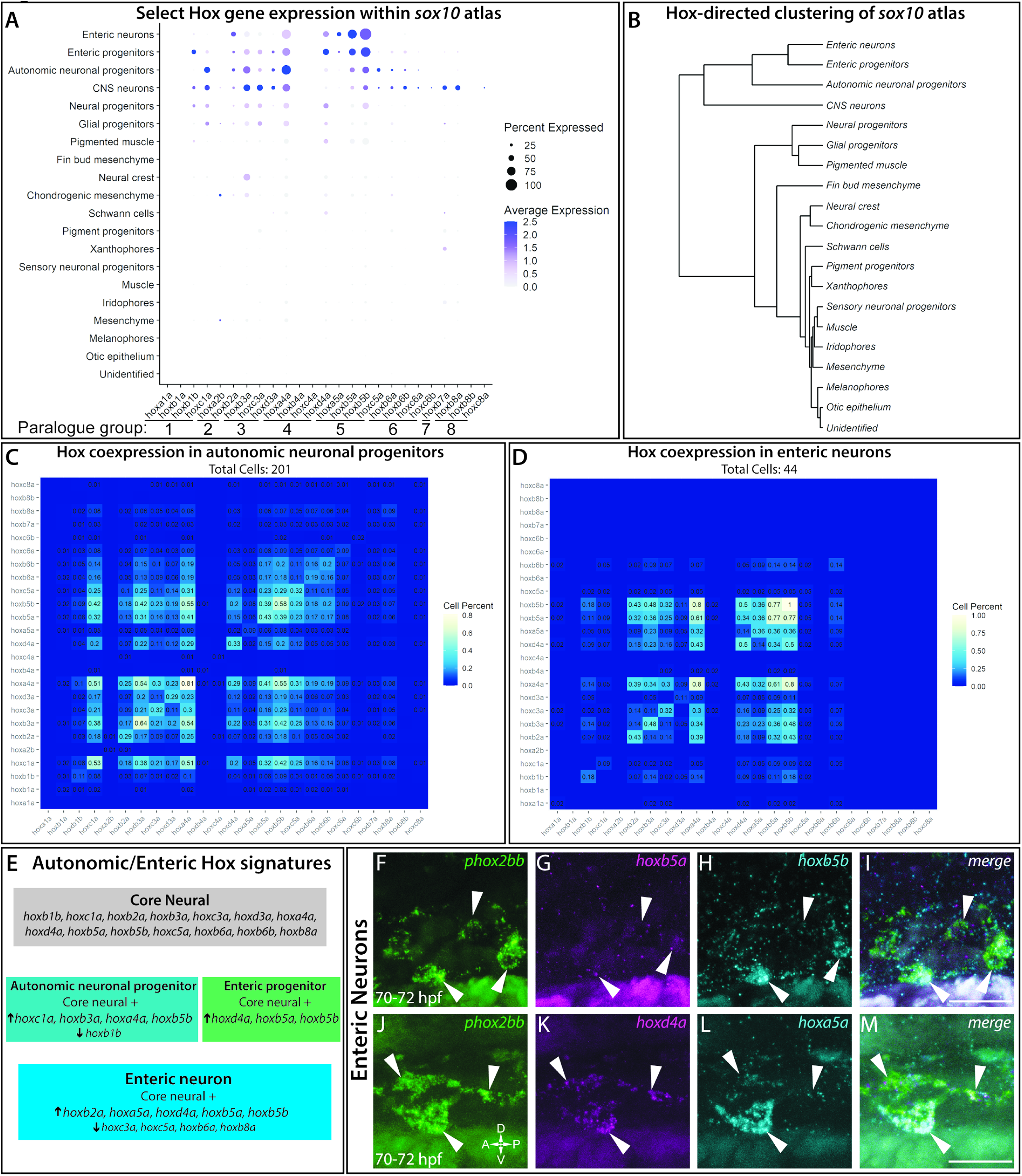
Description of Hox genes expressed across cell lineages within the *sox10*:GFP^+^ atlas. **(A)** Dot plot shows both the mean expression (color) as well as percent of cells (size) per lineage for zebrafish *hox* genes in the first 8 paralogy groups (PG). The full list of *hox* gene expression profiles per lineage can be found in Figure 7 - figure supplement 1. Discrete *hox* profiles discern specific cell types, which is particularly evident in the enteric neuronal cluster. **(B)** Clustering of atlas lineages based *hox* expression profiles groups highlights robust core neural signature, which distinguishes the neural lineages from the remainder of the clades. Neural and glial progenitors formed an intermediate clade between the low-*hox* expressing lineages and the main neural branch. Additionally, the fin bud mesenchyme, which also has a highly distinctive *hox* profile, also forms a distinct clade. Subtle variations in *hox* expression by remaining lineages further are further reflected in the remaining portion of the dendrogram, however these are far less distinct. **(C-D)** Pairwise comparison of the fraction of cells in either the autonomic neural progenitor lineage **(C)** or the enteric neurons **(D)** for the first eight parology groups. Intersection of the gene pairs reflect the fraction of cells with expression for both genes with a log2 Fold change values > 0, with the identical gene intersections along the primary diagon representing the total number of cells which express that gene in the lineage. Enteric neural hox signature was not only specific to this cell population, but also was abundantly coexpressed. **(E)** Summary panel describing the specific autonomic and enteric *hox* signatures detected. We found a common *hox* expression profile, which we refer to as the core signature, that is then modified across the specific lineages. **(F-M)** In situ validation of the chief enteric neural *hox* signature via hybridization chain reaction. *phox2bb* (**F-J)** labels enterically fated neurons at the level of the midgut in larval stage embryos fixed at 70-72hpf. White arrows highlight specific cells of interest. Key hox signature constituents *hoxb5a* **(G)** and *hoxb5b* **(H)** or *hoxd4a* **(K)** and *hoxa5a* **(L)** were found to be co-expressed within *phox2bb* expressing cells (White Arrows). Scale bars in (**I,M)**:50 μm.

In order to better understand the complexities of *hox* codes within specific lineages, we performed a pairwise comparison of each *hox* gene for autonomic and enteric lineages, counting the number of cells which coexpressed each *hox* pair. Examining the autonomic neuronal progenitors (**Figure 7C)** and the enteric neuron populations (**Figure 7D**), both lineages demonstrated pervasive fractions of co-positive cells for combinations of the core *hox* signature. For example, autonomic neuronal cells were enriched with a high fraction of pairwise combinations for *hoxc1a*, *hoxa4a*, *hoxb3a*, and/or *hoxb5b* (**Figure 7C**). The enteric signature was highly enriched in the unique expression of *hoxa5a*, with co-expression for *hoxb5a* (36%) and *hoxb5b* (36%), as well as strong co-expression of *hoxd4a* or *hoxb5a* with *hoxb5b* (**Figure 7D**).

To confirm that *hox* core genes were co-expressed within enteric neurons, we sought to validate their co-expression patterns using HCR probes. As previously described, *hoxd4a*, *hoxa5a*, *hoxb5a,* and *hoxb5b* all exhibited strong hindbrain expression (**Figure 7 - figure supplement 1 B-I**) (Barsh et al., 2017), confirming the specificity of our probes. At 70-72 hpf, as predicted by our analysis, enteric neurons along the level of the midgut, marked by *phox2bb* expression (**Figure 7 F,J**), were *hoxb5a*^+^*/hoxb5b^+^* (**Figure 7 G-I**) and *hoxd4a^+^*/*hoxa5a^+^* (**Figure 7 K-M**). These data confirm that enteric neurons co-express enteric *hox* code genes during their early development.

With respect to the remaining cluster identities (**Figure 7A**), many of the populations showed varied *hox* expression profiles. Both the chondrogenic and general mesenchyme clusters demonstrated *hoxa2b* expression, as well as weak expression for *hoxb2a, hoxb3a*, and *hoxd4a*. Our detection of these *hox* expression profiles was consistent with prior reports that they are expressed within NCC targets toward the posterior pharyngeal arches, as well as migrating NCC (Minoux and Rijli, 2010; Parker et al., 2018, 2019). We detected the distinct identity of the fin bud mesenchyme (Ahn and Ho, 2008; Nakamura et al., 2016) through the expression of *hoxa9b, hoxa10b, hoxa11b, hoxa13b*, *hoxd9a,* and *hoxd12a* (**Figure 7A; Figure 7 - figure supplement 1 P-Q**). The pigment populations, including the pigment progenitors, melanophores, iridophores, and xanthophores, contained generally low levels of *hox* gene expression. Despite this, we still observed a slight variation of *hox* expression among the pigment populations. For example, low levels of *hoxa4a*, *hoxb7a*, *hoxb8a*, *hoxc3a*, and *hoxd4a* were detected among the iridophore population, while only *hoxb7a* was detected within a high fraction of xanthophores (**Figure 7 - figure supplement 1A**). Interestingly, these expression profiles are not shared by the melanophore population, which displayed uniformly very low levels of detectable *hox* expression. Lastly, the muscle, otic, and unidentified cells showed almost no *hox* expression profile, which serves a foil for the specificity of the signatures outlined. We noted that the “pigmented muscle” cluster weakly mirrored the general neural *hox* signature, likely a shared signature more reflective of the axial position of the muscle cells rather than a shared genetic profile, as corroborated by their distinct separation of the clusters on the atlas UMAP (**Figure 6A**).

Overall, these above described *hox* signatures detected within our scRNA-seq atlas indicates that distinct cell types express unique *hox* combinations during their delineation. Description of the *hox* signatures within the *sox10* atlas provides further tools to identify these discrete cell populations, as well as exciting new avenues for further mechanistic investigation.

## DISCUSSION

We present a single cell transcriptomic atlas resource capturing diversity of posterior-residing *sox10*-derived cells during the embryonic (48-50 hpf) to early larval transition (68-70 hpf) in zebrafish. From our analysis, we identified a large number of cell types; including pigment progenitor cells delineating into distinct chromatophores, as well as NCC, glial, neural, neuronal, and mesenchymal cells, extending prior whole embryo-based zebrafish single cell studies (Farnsworth et al., 2020) and expanding the resolution at which these cells have been described to date. We discovered that distinct *hox* transcriptional codes demarcate differentiating neural and neuronal populations, highlighting their potential roles during cell subtype specification. We also uncovered evolutionarily-conserved and novel transcriptional signatures of differentiating enteric neuron cell types, thereby expanding our knowledge of ENS development. Corroborating our transcriptomic characterizations, we validated the spatiotemporal expression of several key cell type markers using HCR. Furthermore, our datasets captured otic vesicle and muscle cells, populations which the *sox10:*GFP line has been characterized as marking, and may be useful for investigating these cell types in the future. Collectively, this comprehensive cell type atlas can be used by the wider scientific community as a valuable resource for further mechanistic and evolutionary investigation of posterior *sox10*- expressing and NCC-derived cells during development and the ontogenesis of neurocristopathies. The atlas is available via an interactive cell browser (https://zebrafish-neural-crest-atlas.cells.ucsc.edu/).

The study of NCC-derived posterior cell types has recently gained increased attention due to their complex and essential roles in vertebrate development (Gandhi et al., 2020; Hutchins et al., 2018; Soldatov et al., 2019; Ling and Sauka-Spengler, 2019). Characterizing the differentiation of NCC-derived cells is important to understand as it will enhance our concept of human health, especially to fields such as stem cell therapeutics and regenerative medicine. Prior studies provide incredible insight into their own respective research systems. Our paper is the first single-cell transcriptomic analysis covering detailed description of the early development of ENS in zebrafish, in addition to analysis of the *sox10*^+^ mesenchyme and pigment cells present during the late embryonic to larval phase. The developmental window we examined, the embryonic to larval transition, is regarded as an ephemeral phase (Singleman and Holtzman, 2014) and as such is expected to contain the dynamic cell differentiation states that we observed within our atlas.

Analysis of pigment populations demonstrated the accuracy and specificity of our transcriptome datasets and identified distinct NCC-derived differentiating chromatophore lineages during the embryonic to larval transition, extending on previous descriptions of pigment cell lineage development performed in older larval and juvenile zebrafish at single cell resolution (Saunders et al., 2019). Specifically, melanophores were identified in the 48-50 hpf dataset (**Figure 2B,C**), while at 68-70 hpf we identified iridophore, xanthophore, pigment progenitor, and two distinct melanophore populations (**Figure 2D-G**). We employed the robust characterization of these pigment populations to validate technical aspects of integrating both time points into a single, cohesive atlas (**Figure 6A**). The atlas shows a common progenitor population branching into both the iridophores and the melanophores, which are composed of the melanophore progenitor cluster from 48-50 hpf and the two melanophore clusters at 68-70 hpf (**Figure 6C**). We validated the results regarding pigment population gene signatures using whole mount HCR on 48-50 hpf and 68-70 hpf embryos (**Figure 2H-J**). Thus, our validation of pigment populations highlights that the *sox10* atlas can be used to identify cell lineages and discover new information regarding their development in zebrafish.

Our datasets captured the transition from enteric neural progenitor to differentiating enteric neuron subtype (**Figure 4,5,6**). We found that in both 48-50 and 68-70 hpf datasets, the expression of *elavl3*, *phox2bb*, *ret*, and *gfra1a* transcripts were present (**Figure 4,5**); however, enteric progenitor populations at 48-50 hpf still retained a NCC signature, marked by *crestin* and *foxd3*, among others (**Figure 4**). The combined atlas revealed broad transcriptional states captured within enteric neural progenitors and enteric neurons (**Figure 6D**), whereby *elavl3*, *phox2bb*, *ret*, *ngfrb*, and *gfra1a* could be seen extending throughout the neural/neuronal regions of the atlas. A similar enteric progenitor population consisting of *Sox10*, *Ret*, *Phox2b*, and *Elavl4* was identified by scRNA-seq in the mouse (Lasrado et al., 2017), indicating zebrafish express conserved enteric programs. Notably, genes that encode for neurochemicals within enteric neurons were detected in the enteric clusters, with *nos1* and *vipb* being most prominent (**Figure 6D**) and co-expressed in a subset of cells among the enteric neuron population along the foregut (**Fig 5A-C**). Collectively, these results regarding enteric populations suggest that the atlas likely reflects cells captured across a spectrum of differentiation states, with immature neurons reflecting the onset of *elavl3* expression, while others, such as the *nos1^+^*/*vipb^+^* subpopulation, representing cells further along a differentiation trajectory.

A recent scRNA-seq study performed using E15.5 mice, a time point further along in ENS development when compared to our zebrafish study described here, suggests that *Nos1^+^*/*Vip^+^* cells represent a post-mitotic immature neuron population capable of branching into excitatory and inhibitory neurons via subsequent differentiation mediated by lineage-restricted gene expression (Morarach et al., 2020). Their model posits that *Nos1^+^*/*Vip^+^*/*Gal^+^* enteric neurons are capable of assuming an intrinsic primary afferents neuron (IPAN) signature, characterized by the loss of *Vip* and *Nos1*, and the gain of *Calb*, *Slc18a2*/*3*, and *Ntng1*; a process regulated by transcription factors, *Pbx3* and *Etv1*. This model of IPAN formation appears congruent with a previous birth dating study performed in mice, where researchers demonstrated the transient expression of *Nos1* in enteric neurons (Bergner et al., 2014). We wondered if the IPAN gene expression signature was evolutionarily conserved in zebrafish. Testing this spatiotemporal gene signature model in our own datasets, we asked if the *nos1^+^*/*vipb^+^* population represented a snapshot of immature enteric neurons. We found that *pbx3b*, *etv1*, *calb2a*, *slc18a3a*, *ache*, *vipb*, and *nos1* were all expressed in differentiating enteric neuron clusters (**Figure 5D-E; Figure 5-figure supplement 1A**), likely reflecting their transition to an IPAN fate in our 68-70 hpf dataset. Intriguingly, we discovered that the markers tightly mapped to a subpopulation of cells in an enteric neuron sub-cluster (**Figure 7E**, red arrows) and that *nos1* was either absent or expressed at lower levels than other enteric subpopulations (**Figure 5C; Figure 5-figure supplement 1**), a finding that corroborates the proposed mammalian model. Our observations in zebrafish suggest that we captured a transitional time point where subsequent differentiation is just being initiated and suggests an evolutionarily-conserved mechanism of ENS formation across vertebrate species.

Within the enteric neuron subpopulation in our dataset, we discovered the enrichment of opioid pathway members. Opioids have been known effectors of gastrointestinal function for centuries based on their use as an anti-diarrheal medicine (De Luca and Coupart, 1996). Modern research has allowed us to understand the mechanism by which opioids exert their effect on gut function, namely, through the signaling mediated by the superfamily of G-protein coupled opioid receptors that serve to regulate neurotransmission within the adult ENS (Holzer, 2004; De Luca and Coupart, 1996; Wood and Galligan, 2004). This superfamily of opioid receptors is comprised of mu- (MOR), delta- (DOR), kappa- (KOR), and nociception/orphan FQ (N/OFQ) peptide (NOP)-receptors (Donica et al., 2013; Holzer, 2004). The expression of opioid receptors has previously been shown in inhibitory interneurons found within the adult ENS and are believed to inhibit gut peristalsis and secretion by inducing hyperpolarization of inhibitory neurons, resulting in the deregulation of the excitatory circuits that drive smooth muscle contraction within the gut (DiCello et al., 2020; Lay et al., 2016; Wood and Galligan, 2004).

According to the CDC, licit and illicit opioid abuse has risen 400% in pregnant women between 1999-2014, which has in turn led to a 400% increase in the number of infants born with neonatal abstinence syndrome (NAS), a multisystemic clinical condition characterized by a wide array of symptoms including autonomic nervous system and gastrointestinal dysfunction (Raffaeli et al., 2017).

While the presence and inhibitory effect of opioid receptors is well characterized within the adult ENS, the role of opioid signaling during the earliest stages of enteric neuron maturation and ENS formation has yet to be investigated. Indeed, a recent study performed in zebrafish found that the NOP-receptor, *oprl1* was expressed within 7 dpf enteric neurons following bulk RNA sequencing (Roy-Carson et al., 2017). However, this data represents a developmental stage where zebrafish are characterized as free swimming larva that display feeding behavior and digestive capability representative of a functioning and more mature ENS (Cassar et al., 2018). As such, our 68-70 hpf dataset, in which we detected the expression of the two receptors, *oprl1* and *oprd1b,* respectively representing (NOP) and (DOR) class opioid receptors (**Figure 5H**), represents the earliest stage in which these opioid receptors have been shown to be expressed within the early developing ENS. The presence of opioid receptors within the ENS at 68-70 hpf, a time when immature enteric neurons are continuing to migrate and pattern within the developing embryo, highlights an important area of research focusing on the interplay between opioid use and fetal ENS development, a reality that has been on the rise in recent decades as opioid abuse continues to increase.

Finally, we have elucidated a comprehensive, combinatorial code of Hox transcription factor expression which define specific cell lineages within the context of the *sox10* atlas. The fin bud mesenchyme, pigment populations, and neural fates presented the most specific *hox* codes (**Figure 7**). The previously well characterized signature identified in the fin bud served as validation of our dataset’s curation for further analysis of the *hox* signatures. Among the neural lineages, we identified a previously undescribed *hox* signature demarcating the developing enteric neuron population in zebrafish: high relative expression of *hoxb2a*, *hoxa5a*, *hoxd4a*, *hoxb5a* and *hoxb5b,* while also exhibiting low expression of *hoxc3a, hoxc5a, hoxb6a* and *hoxb8a* (**Figure 7**). While previous bulk microarray studies of differentiating enteric neurons from humans and mice have been shown to express orthologs to the former signature (Heanue and Pachnis, 2006; Memic et al., 2018), our analysis extends knowledge to comprehensively show for the first time specific combinatorial *hox* expression at a single cell resolution, thereby providing a heretofore unknown readout of *hox* heterogeneity among nascent enteric cells. The conservation of an enteric *hox* signature between zebrafish and other systems point toward larger conservation of function, which may facilitate translation of future findings between models. These findings imply interesting potential models wherein combinations of *hox* codes may represent a molecular address designating the axial site of origination for migrating NCC or indicate dynamic expression profiles which are modified during the NCC to enteric neuron developmental course. Further work is required to test these possible models as well as the functional requirement of the constituent members of the *hox* code the developing zebrafish ENS. Considered collectively, these data lend support to a model in which overlapping expression domains of *hox* genes may facilitate enteric neural subtype differentiation, similar to their function in hindbrain and spinal neurons (Philippidou and Dasen, 2013).

In summary, our study greatly increases our foundational understanding of NCC-derived cell fates, as well as other *sox10*^+^ posterior cell types in zebrafish, thereby complementing ongoing studies in mammalian models and expanding fundamental knowledge of how cells diversify in developing organisms. The spatiotemporal information contained within our zebrafish atlas will serve as a resource for the developmental biology, stem cell, evolutionary biology and organogenesis communities.

## METHODS & MATERIALS

### KEY RESOURCE TABLE

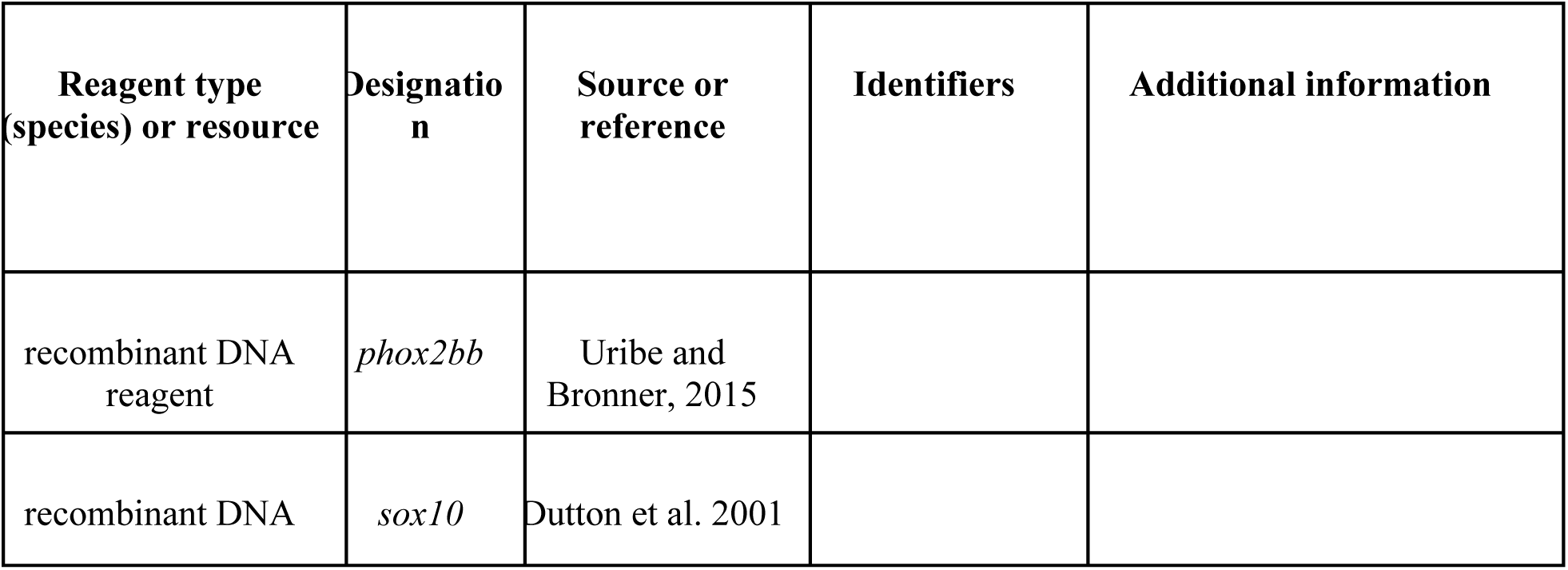
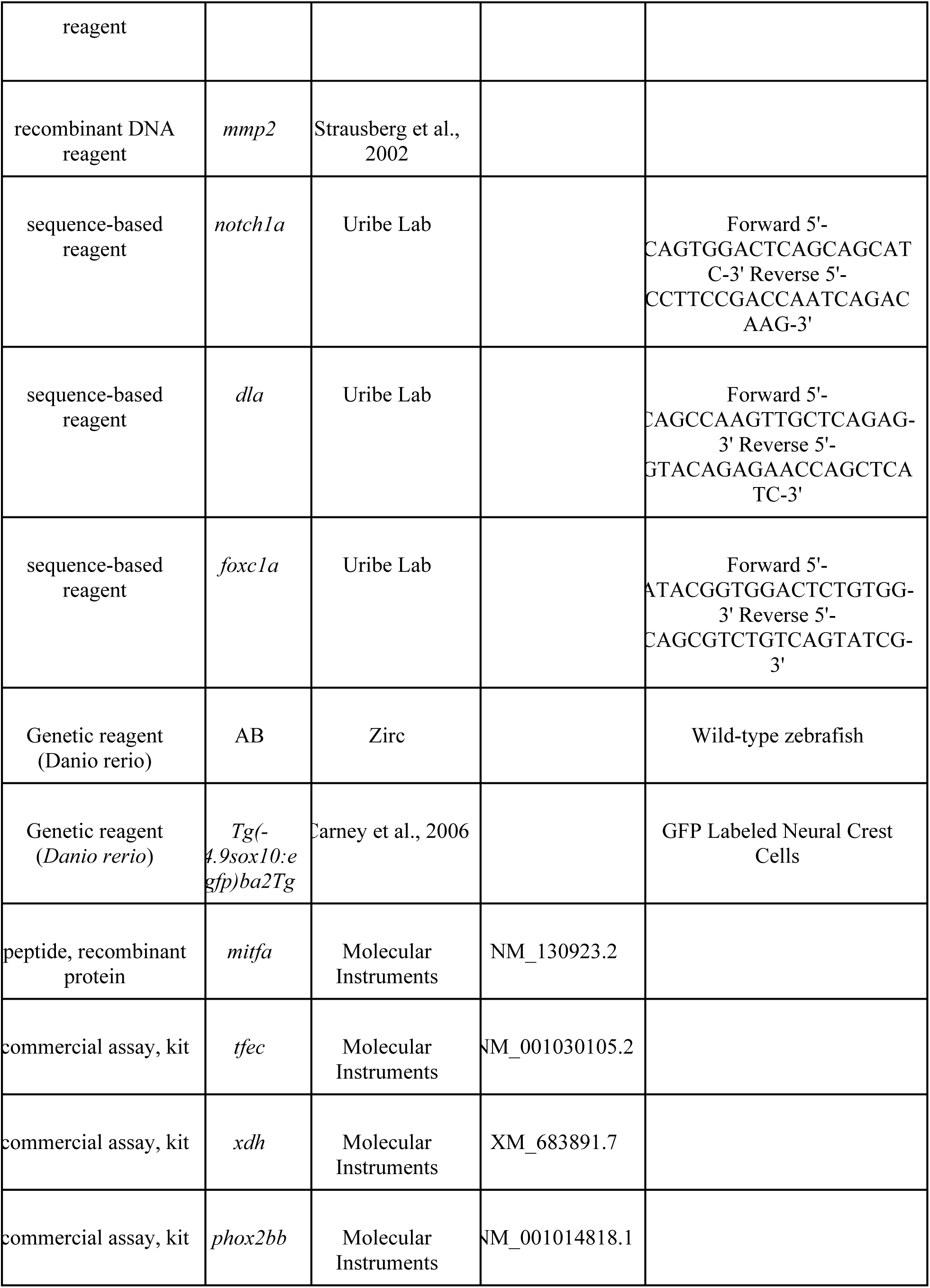
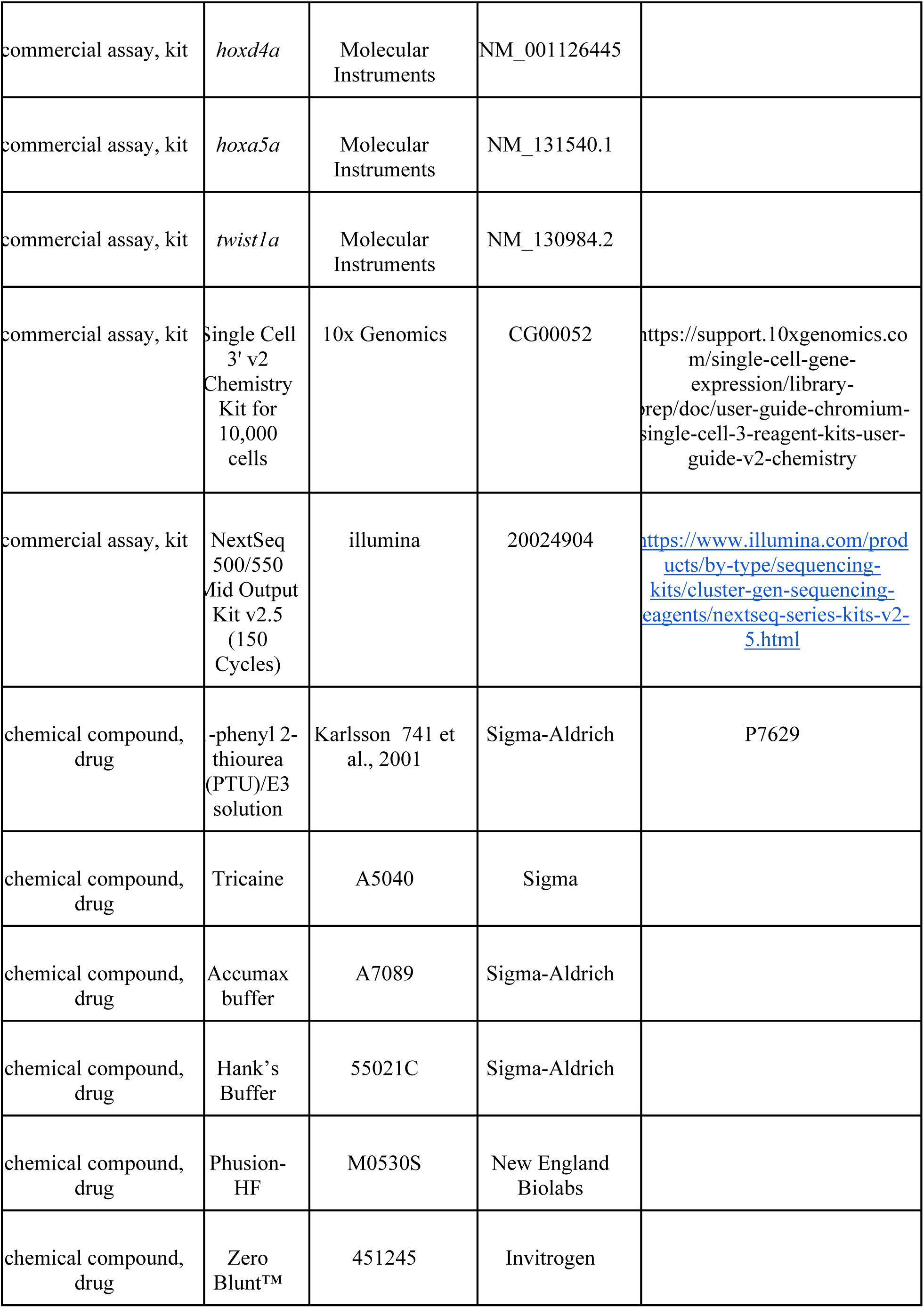
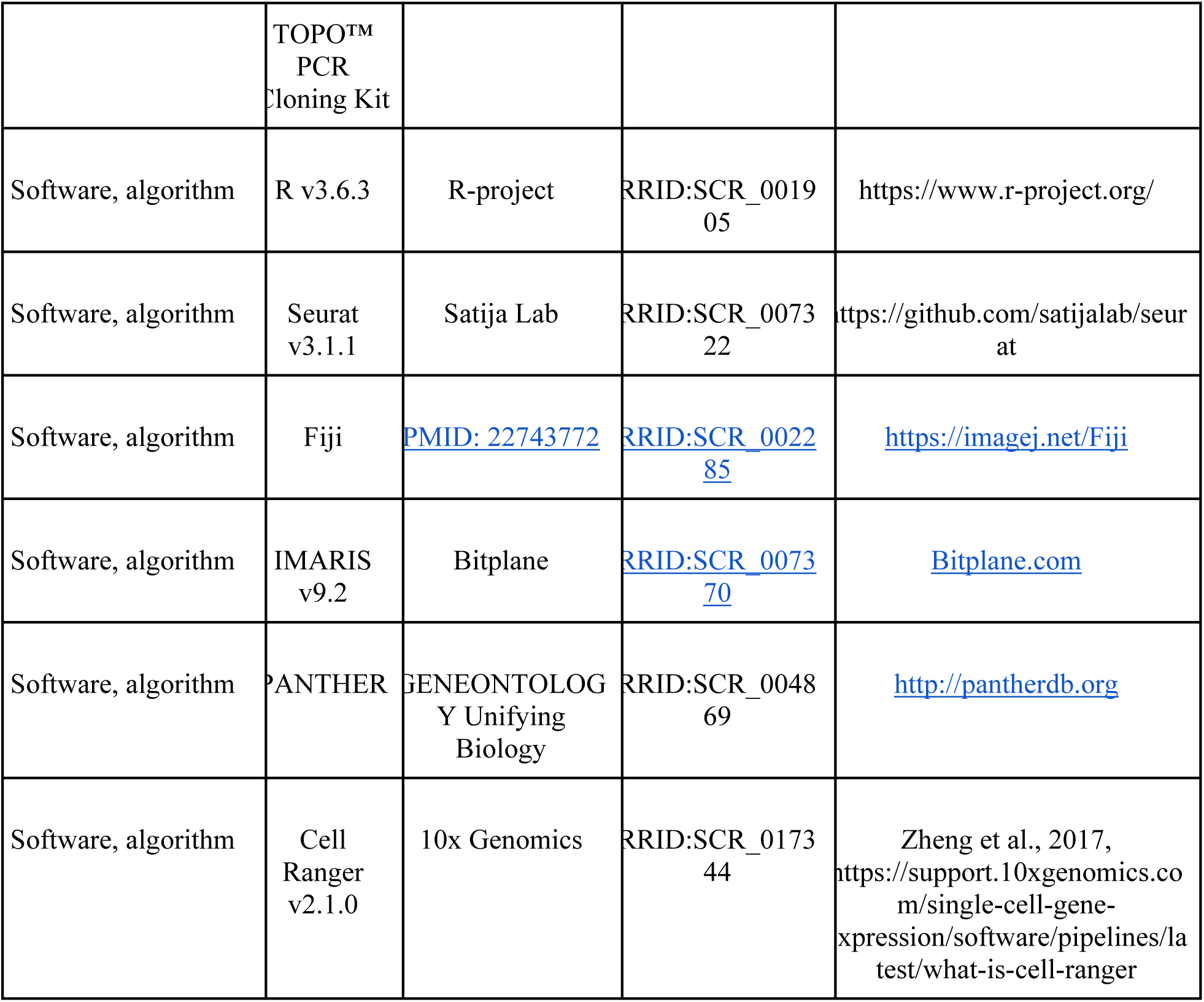

### Animal Husbandry, Care, & Synchronous Embryo Collection

Groups of at least 15 adult Tg(*-4.9sox10:*GFP)^ba2Tg^ (Carney et al., 2006) zebrafish (*Danio rerio*) males and 15 females from different tank stocks were bred to generate synchronously staged embryos across several clutches. All embryos were cultured in standard E3 media until 24 hours post fertilization (hpf), then transferred to .003% 1-phenyl 2-thiourea (PTU)/E3 solution (Karlsson et al., 2001), to arrest melanin formation and enable ease of GFP sorting. While it has been suggested that high concentration (.03%) PTU incubation prior to 22 hpf may cause organism-wide effects, .003% PTU application after 22 hpf has been shown to have no major effects on NCC survival or pigment cell formation, as described (Bohnsack et al., 2011). Embryos were manually sorted for GFP expression and synchronously staged at 24 hpf. Care was taken such that embryos which exhibited developmental delay or other defects were removed prior to collection. All work was performed under protocols approved by, and in accordance with, the Rice University Institutional Animal Care and Use Committee (IACUC).

### Isolation of Tissue & Preparation of Single Cell Suspension

100 embryos between 48-50 hpf and 100 larvae between 68-70 hpf were dechorionated manually and then transferred to 1X sterile filtered PBS, supplemented with 0.4% Tricane (Sigma, A5040) to anesthetize. Tissue anterior to the otic vesicle and tissue immediately posterior to the anal vent was manually removed using fine forceps in 48-50 hpf embryos, while tissue anterior to the otic vesicle was removed from 68-70 hpf larvae, as schematized in Figure 1. This was to capture as many posterior *sox10*:GFP^+^ cells in the later time point as possible. Remaining tissue segments were separated into nuclease-free tubes and kept on ice immediately following dissection. Dissections proceeded over the course of 1 hour. To serve as control for subsequent steps, similarly staged AB WT embryos were euthanized in tricaine and then transferred to sterile 1X PBS. All following steps were conducted rapidly in parallel to minimize damage to cells: Excess PBS was removed and tissue was digested in 37°C 1X Accumax buffer (Sigma-Aldrich, A7089) for 30-45 minutes to generate a single cell suspension for each sample. At 10 minute intervals, tissue was gently manually disrupted with a sterile pipette tip. As soon as the tissue was fully suspended, the cell solutions were then transferred to a fresh chilled sterile conical tube and diluted 1:5 in ice cold Hank’s Buffer (1x HBSS; 2.5 mg/mL BSA; 10μM pH8 HEPES) to arrest the digestion. Cells were concentrated by centrifugation at 200 rcf for 10 minutes at 4°C. Supernatant was discarded carefully and cell pellets were resuspended in Hank’s Buffer. Cell solution was passed through a 40 μm sterile cell strainer to remove any remaining undigested tissue and then centrifuged as above. Concentrated cells were resuspended in ice cold sterile 1X PBS and transferred to a tube suitable for FACS kept on ice. The 48-50 hpf and 68-70 hpf experiments were performed on completely separate dates and times using the above described procedures.

### Fluorescent Cell Sorting, & Single Cell Sequencing

Fluorescent Assisted Cell Sorting (FACS) was performed under the guidance of the Cytometry and Cell Sorting Core at Baylor College of Medicine (Houston, TX) using a BD FACSAria II (BD Biosciences). Zebrafish cells sorted via GFP fluorescence excited by a 488 nm laser, relying on an 85 μm nozzle for cell selection. Detection of GFP^+^ cells was calibrated against GFP^-^ cells collected from AB wildtype embryos, as well as GFP^+^ cells collected from the anterior portions of the *sox10:*GFP embryos. Optimal conditions for dissociated tissue inputs (number of embryos needed, etc.) and FACS gating was determined via pilot experiments prior to collection for subsequent scRNA-seq experiments. Sample preparation for scRNA-seq was performed by Advanced Technology Genomics Core (ATGC) at MD Anderson (Houston, TX). 4905 and 4669 FACS-isolated cells for the 48-50 and 68-70 hpf datasets were prepared on a 10X Genomics Chromium platform using 10X Single Cell 3’ V2 chemistry kit for 10,000 cells. cDNA libraries were amplified and prepared according to the 10X Genomics recommended protocol, with details provided in **Figure 1-figure supplement 1C.** A 150 cycle Mid-Output flow cell was used for sequencing on an Illumina NextSeq500. Sequencing was aligned at MD Anderson ATGC to the DanioGRCz10 version of the zebrafish genome using the 10X Genomics Cell Ranger software (v2.1.0) (Zheng et al., 2017). Gene reads per cell were stored in a matrix format for further analysis.

### Data Processing & Analysis

The 10x genomics sequencing data was then analyzed using Seurat (Satija et al., 2015, Stuart et al. 2019, Butler et al., 2018) v3.1.1 software package for R, v3.6.3 (R Core Team, 2020). The standard recommended workflow was followed for data processing. Briefly, for both the 48-50 hpf and 68-70 hpf datasets, cells which contained low (<200) or high (>2500) genes were removed from analysis. Gene expression was normalized using the NormilizeData command, opting for the LogNormalize method (Scale factor set at 10,000) and further centered using the ScaleData command. Variable features of the dataset were calculated with respect to groups of 2,000 genes at a time. Both datasets were evaluated considering the first 20 principle components (PC) as determined by the RunPCA command with a resolution of 1.2 for PCA, tSNE, and UMAP analyses. The appropriate PCs were selected based on a Jack Straw analysis with a significance of P < 0.01, as generated by the JackStraw command.

Clustering was performed using FindNeighbors and FindClusters in series. We identified 19 clusters in the 48-50 hpf dataset and 23 clusters in the 68-70 hpf dataset. Significant genes for each cluster were determined via a Wilcoxon Rank Sum test implemented by the FindAllMarkers command. From these expressed gene lists within each cluster, all cluster identities were manually curated via combinatorial expression analysis of published marker genes in the literature, zfin.org and/or bioinformatics GO term analysis via the Panther Database.

Generation of the merged atlas was performed via the FindIntegrationAnchors workflow provided in the Standard Workflow found on the Seurat Integration and Label Transfer vignette. Clustering was performed for the atlas based on the first 20 PCs, consistent with the original datasets. Subsets of the atlas discounted any spuriously sorted cells for clarity. All features plots represent expression values derived from the RNA assay. Sub-clustering of the enteric clusters was performed by sub-setting clusters 5 and 12 from the 68-70 hpf dataset and reinitializing the Seurat workflow, as described above. Clusters were identified based on the first 6 PCs. Detection of cell cycle phase was conducted following the Cell cycle and scoring vignette. Genes used for identification of cell cycle phases can be found in the supplementary table (Figure 1-figure supplement 3). Pairwise *hox* analysis was conducted using tools from the Seurat package in R by assessing the number of cells which had expression for *hox* gene pairs queried with a log2 fold change greater than 0. Dendrogram Cluster trees were generated using Seurat’s BuildClusterTree function.

### Whole mount in situ Hybridization

cDNAs for *foxc1a*, *notch1a*, and *dla* were amplified via high fidelity Phusion-HF PCR (NEB) from 48 hpf AB WT cDNA libraries using primers in the Key resources table. PCR products were cloned using the Zero Blunt™ TOPO™ PCR Cloning Kit (Invitrogen), as per manufacturer protocols, and sequenced validated. Plasmids encoding *phox2bb*, *sox10*, *mmp2* were generously sourced as listed in the key resources table. Antisense digoxigenin (DIG)- labeled riboprobes were produced from cDNA templates of each gene. AB wild type embryos were treated and stained to visualize expression as previously described in (Jowett and Lettice, 1994). Following *in situ* reactions, embryos were post-fixed in 4% Paraformaldehyde (PFA) and mounted in 75% Glycerol for imaging. A Nikon Ni-Eclipse Motorized Fluorescent upright compound microscope with a 4X objective was used in combination with a DS-Fi3 color camera. Images were exported via Nikon Elements Image Analysis software.

### Whole mount Hybridization Chain Reaction

HCR probes were purchased commercially (Molecular Instruments Inc., CA) and were targeted to specific genes based on their RefSeq ID (Key resources table). Whole mount HCR was performed according to the manufacturer’s instructions (v3.0, Choi et al., 2016, 2018) on *sox10:*GFP*^+^* or AB embryos previously fixed at the appropriate stage in 4% PFA.

### Confocal Imaging & Image Processing

Prior to imaging, embryos were embedded in 1% low melt agarose (Sigma) and were then imaged using an Olympus FV3000 Laser Scanning Confocal, with a UCPlanFLN 20×/0.70NA objective. Confocal images were acquired using lambda scanning to separate the Alexafluor 488/Alexafluor 514 or the Alexafluor 546/Alexafluor 594 channels. Final images were combined in the FlowView software and exported for analysis in either Fiji (Rueden et al., 2017; Schneider et al., 2012; Schindelin et al., 2012) or IMARIS image analysis software (Bitplane). Figures were prepared in Adobe Photoshop and Illustrator software programs, with some cartoons created via BioRender.com.

### Data Availability

The raw sequence read files and processed cellular barcode, gene, and matrix files produced by CellRanger are available in the National Center for Biotechnology Information’s (NCBI) Gene Expression Omnibus (GEO) database (https://www.ncbi.nlm.nih.gov/geo/), accession number: GSE152906. The atlas/associated processed Seurat objects are available on the University of California, Santa Cruz (UCSC) Cell Browser (https://zebrafish-neural-crest-atlas.cells.ucsc.edu/<u>). Code written for Seurat data analysis is available on GitHub (</u>https://github.com/UribeLabRice).

## ACKNOWLEDGEMENTS

Funding for this project was provided by Rice University, Cancer Prevention & Research Institute of Texas (CPRIT) Recruitment of First-Time Tenure Track Faculty Members (CPRIT-RR170062) and the NSF CAREER Award (1942019) awarded to R.A.U., a Houston Livestock Show & Rodeo Research Award to J.A.M. and P.A.B., and a SDB Choose Development! Fellowship award to J.L.W. We acknowledge the Cytometry and Cell Sorting Core at Baylor College of Medicine, which is funded from the CPRIT Core Facility Support Award (CPRIT-RP180672), the NIH (P30 CA125123 and S10 RR024574), and the expert assistance of Joel M. Sederstrom for assistance with flow cytometry. Single cell library preparation, Illumina sequencing, and Cell Ranger alignment was facilitated by Advanced Technology Genomics Core at MD Anderson Cancer Research Center funded by CA016672(ATGC). IMARIS image analysis was performed using Rice University’s Shared Equipment Authority (SEA) IMARIS workstation. We thank George Eisenhoffer and Oscar Ruiz (MD Anderson) for advice regarding flow cytometry and single-cell RNA-seq methodology. We thank Sarah Kucenas (University of Virginia) for helpful advice on glial populations. We thank Robert Naja and Robyn Fenty for technical assistance.

## Figure Legends

**Figure 1- figure supplement 1.**
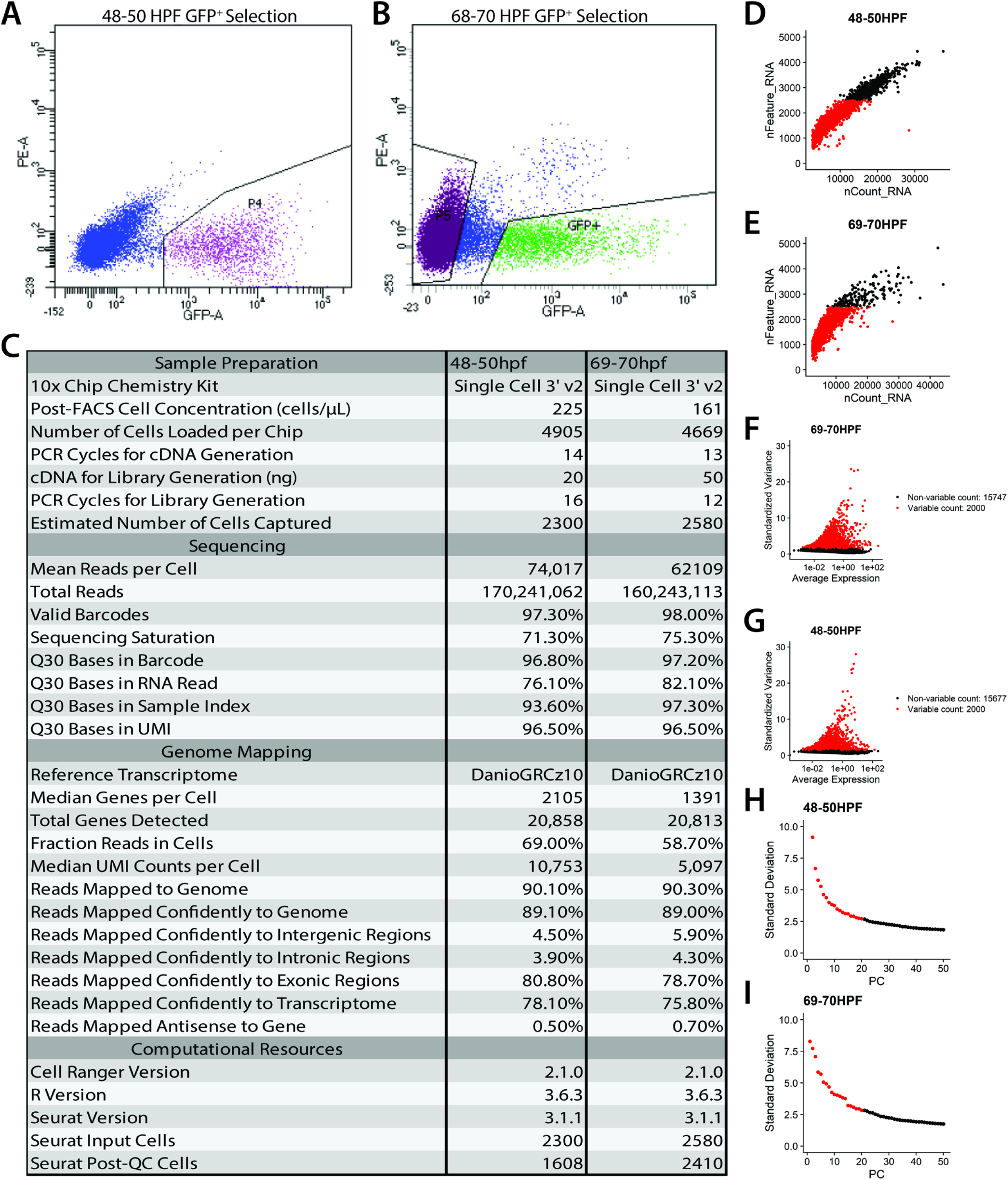
Statistics on generation and filtering of single cell transcriptomes at 48-50 hpf and 68-70 hpf. **(A,B)** Fluorescence activated cell sorting plots highlighting the GFP^+^ cell population sorted at 48-50 hpf **(A)** and 68-70 hpf **(B)**. **(C)** Table of general statistics pertaining to the sequencing and alignment of reads from the Cell Ranger pipeline. Additional metrics provided were derived from the Seurat R package as described in the Materials & Methods section. **(D,E)** Plots showing the feature selection for both the 48-50 hpf **(D)** and 68-70 hpf **(E)** datasets. Cells were selected such that they had fewer than 2500 features to reduce spuriously sorted cells. **(F,G)** Top 2000 most variably expressed genes were identified and used for further downstream identification of significant principal components, as described in the Materials & Methods section. **(H,I)** Most significant principle components (top 20 for both datasets) were selected to be used for subsequent cluster identification and cell embedding in tSNE and UMAP spaces.

**Figure 1-figure supplement 2.**
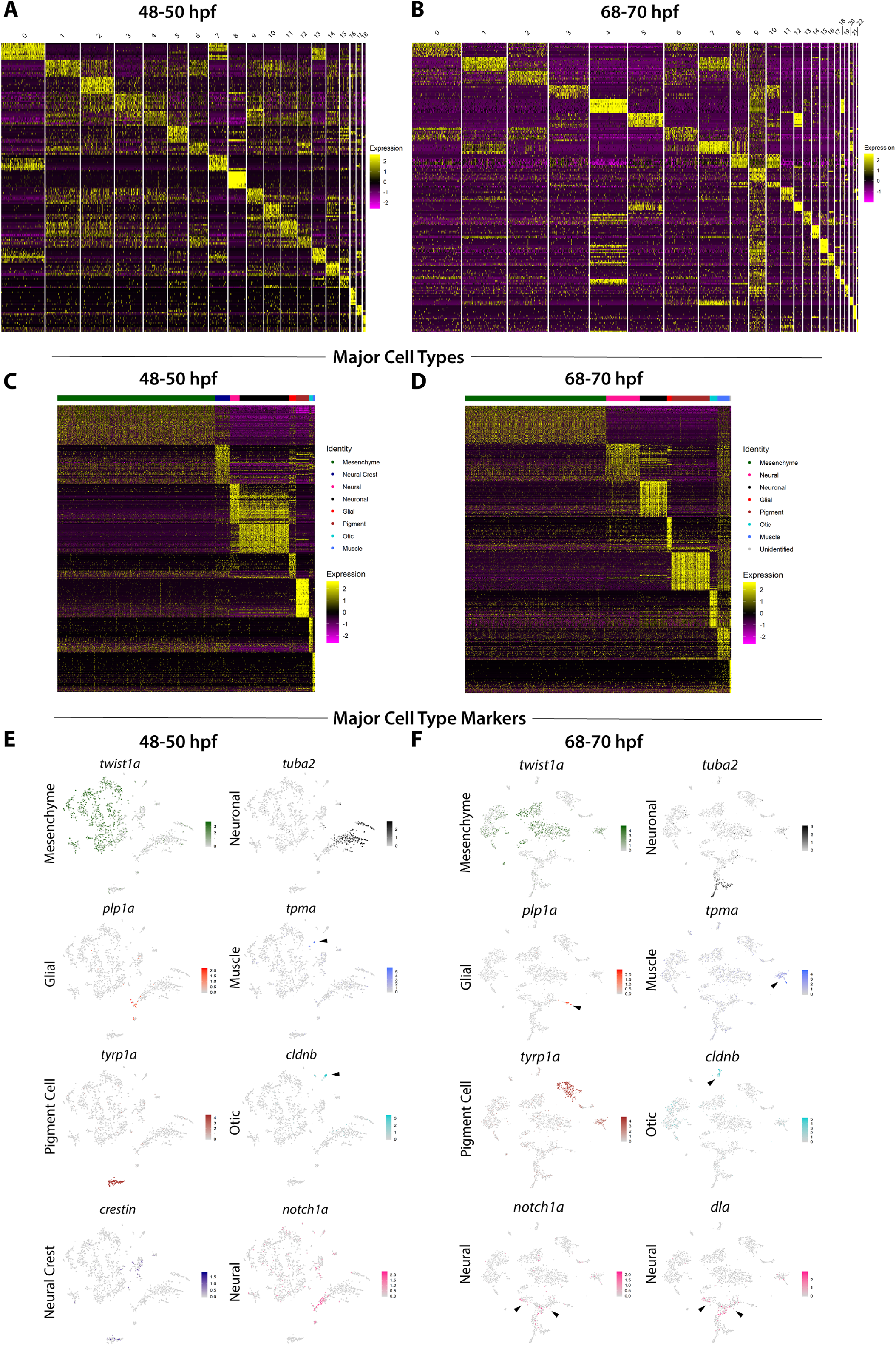
Major cell type annotations among *sox10*:GFP^+^ cells. **(A,B)** Heatmap summarizing the top 10 genes significantly expressed in each cluster, for 48-50 and 68-70 hpf, respectively. Relative expression levels within each cluster is summarized within the color key, where yellow to magenta color indicates high to low gene expression levels. **(C,D)** Heatmaps summarizing the top 30 genes significantly expressed among the major cell types identified among *sox10*:GFP^+^ cells, for 48-50 and 68-70 hpf, respectively. Relative expression levels within each major cell type cluster is summarized within the color key, where yellow to magenta color indicates high to low gene expression levels. **(E,F)** tSNE plots depicting the major cell type classification representative gene marker for each major cell type category, for 48-50 and 68-70 hpf, respectively. Relative expression levels are summarized within the color keys, where color intensity is proportional to expression level of each gene depicted.

**Figure 1-figure supplement 3.**
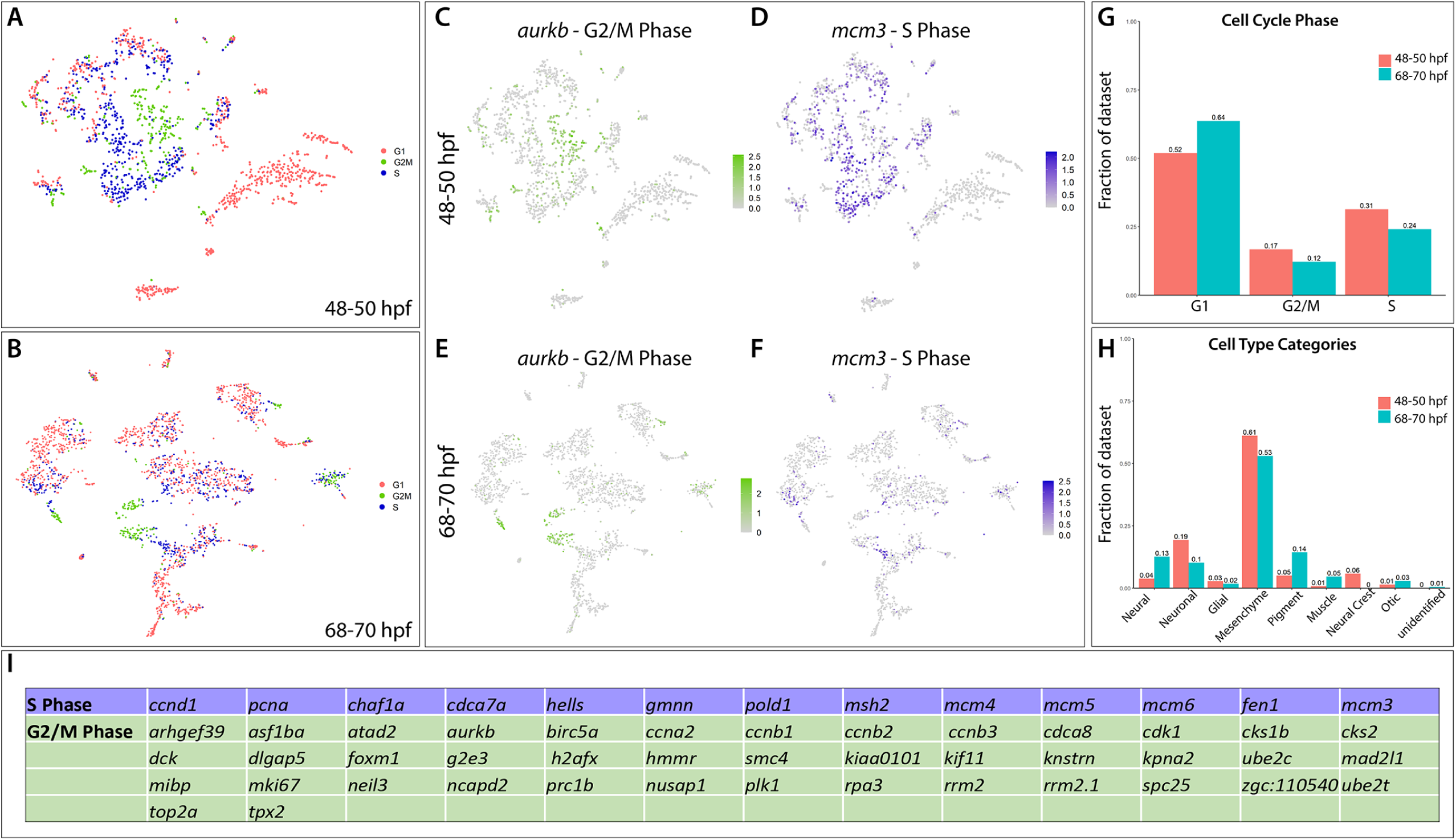
Major cell type categories and cell cycle distributions of the scRNA-seq datasets. **(A,B)** tSNE plots summarizing the G1, S and G2/M phase cell cycle phase occupancies of the cells in the 48-50 and 68-70 hpf time points, respectively. **(C,E)** A tSNE plot depicting the expression of *aurkb*, a G2/M phase marker, within the 48-50 and 68-70 hpf datasets, respectively. Relative expression levels are summarized within the color keys, where color intensity is proportional to expression level of each gene depicted. **(D,F)** A tSNE plot depicting the expression of *mcm3*, a S phase marker, within the 48-50 and 68-70 hpf datasets, respectively. Relative expression levels are summarized within the color keys, where color intensity is proportional to expression level of each gene depicted. **(G)** Bar graphs summarizing the cell cycle phase occupancies, as a fraction of cells within the total datasets for each time point. **(H)** Bar graphs summarizing the major cell type categories, as a fraction of cells within the total datasets for each time point. **(I)** Table summarizing the cell cycle genes used to demarcate cell cycle phase occupancy categories within the scRNA-seq datasets.

**Figure 1-figure supplement 4.**
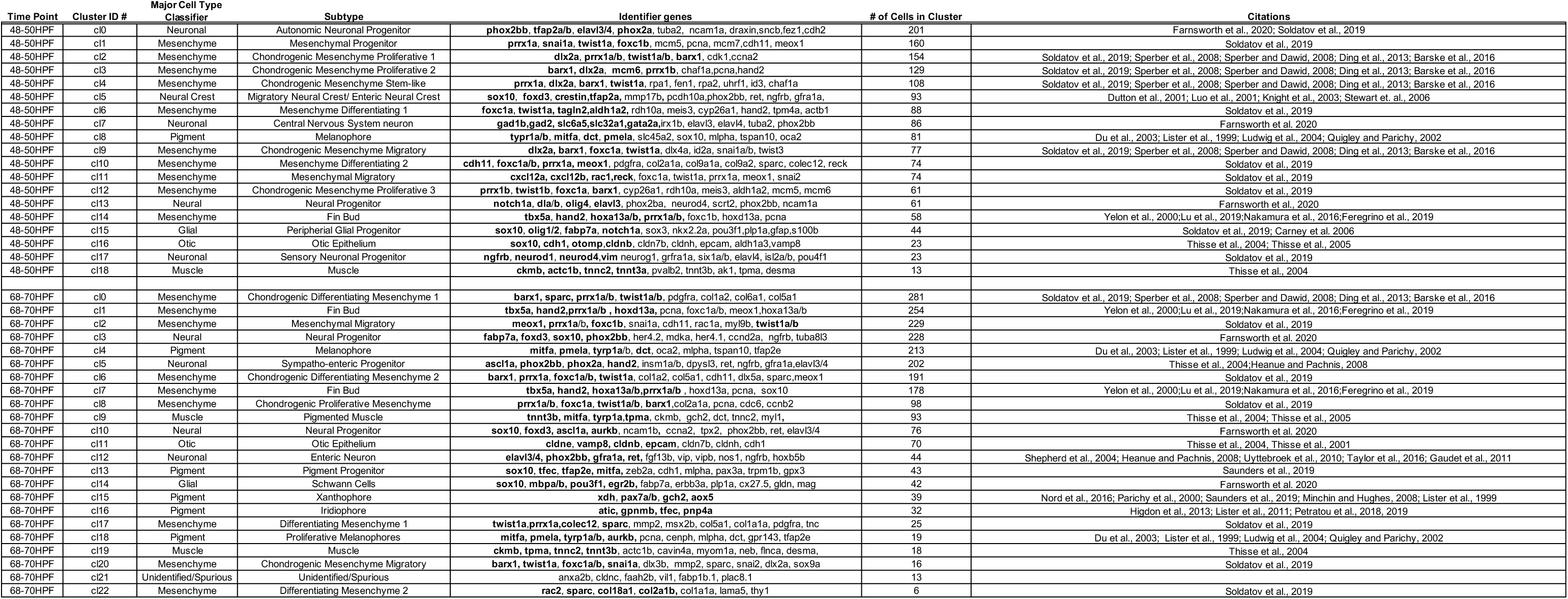
Table summarizing the top identity markers used for major cell type and subtype cellular classifications for each cluster at 48-50 and 68-70 hpf.

**Figure 1-figure supplement 5.**
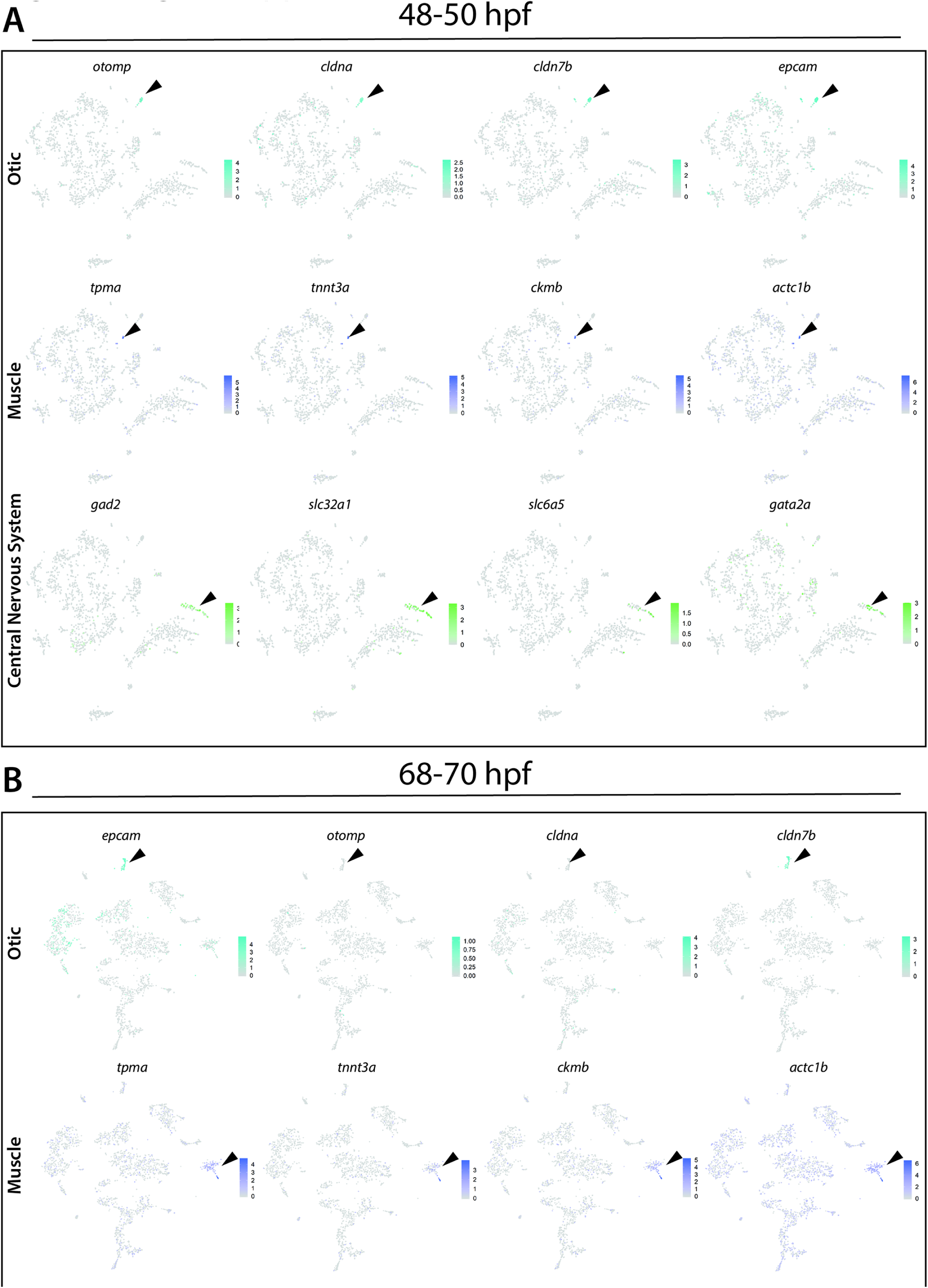
Identification of otic vesicle, muscle, and central nervous system (CNS) cellular populations **(A)** Panel of tSNE feature plots at 48-50 hpf that identify combinatorial expression of otic vesicle (*otomp, cldna, cldn7b and epcam),* muscle (*ckmb, actc1b, tnnt3a and tpma),* or CNS *(slc32a1, gad1b, slc6a5, gata2a)* markers. Cluster of interest denoted by black arrows. **(B)** Panel of tSNE feature plots at 68-70 hpf that identify combinatorial expression of otic vesicle markers *(otomp, cldna, cldn7b and epcam)* or muscle *(ckmb, actc1b, tnnt3a and tpma).* Cluster of interest denoted by black arrows.

**Figure 1-figure supplement 6.**
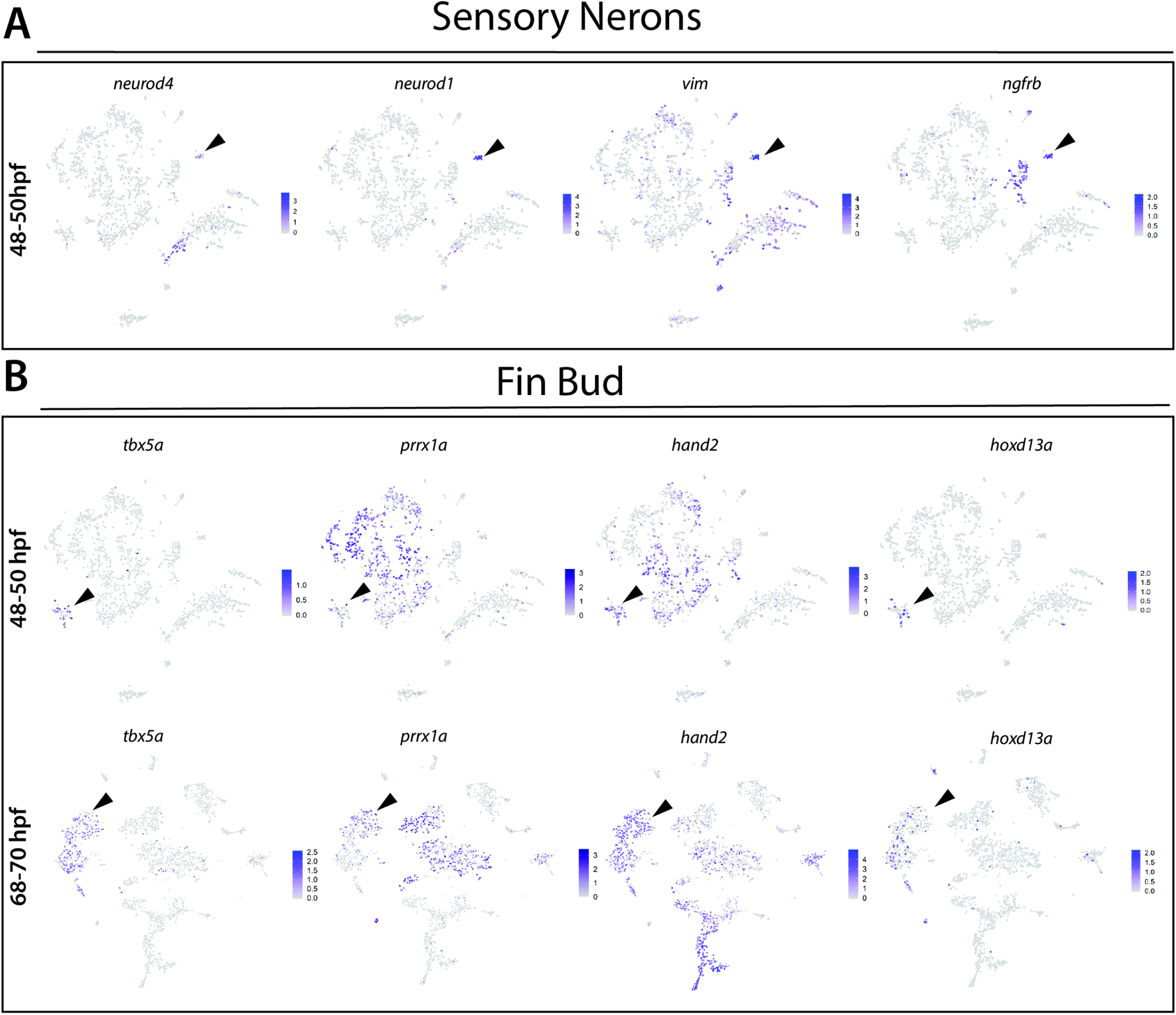
Identification of fin bud and sensory neuronal progenitor cellular populations **(A)** Panel of tSNE feature plots of sensory neuronal progenitors at 48-50 hpf that show combinatorial expression of *neurod4, neurod1, vim, and ngfrb,* Cluster of interest denoted by black arrows. **(B)** Panel of tSNE feature plots of fin bud makers at 48-50 hpf (top) and 68-70 hpf (bottom) that show combinatorial expression of *tbx5a, hand2, hoxd13a,* and *prrx1a.* Cluster of interest denoted by black arrows.

**Figure 1-source data 1. List of marker genes per cluster in the *sox10*:GFP scRNA-seq datasets.**

Tables reporting the Seurat output for genes for each cluster at the 48-50 hpf (Tab 1) and 68-70 hpf (Tab 2) time points, including p-values (<0.01), average log-fold change (≥.25), adjusted p-values (≤1.0), pct.1 summarizing proportion of cells expressing the individual gene in the cluster, pct.2 showing the proportion of cells expressing the individual gene in all other clusters in the dataset.

**Figure 2-source data 1. Melanophore populations, shared and unique genes at 68-70 hpf.**

Table summarizing the genes (“elements”) exclusively found in Cluster 4, exclusively found in Cluster 18, and genes found in both Cluster 4 and 18 at the 68-70 hpf time point scRNA-seq dataset.

**Figure 3-figure supplement 1.**
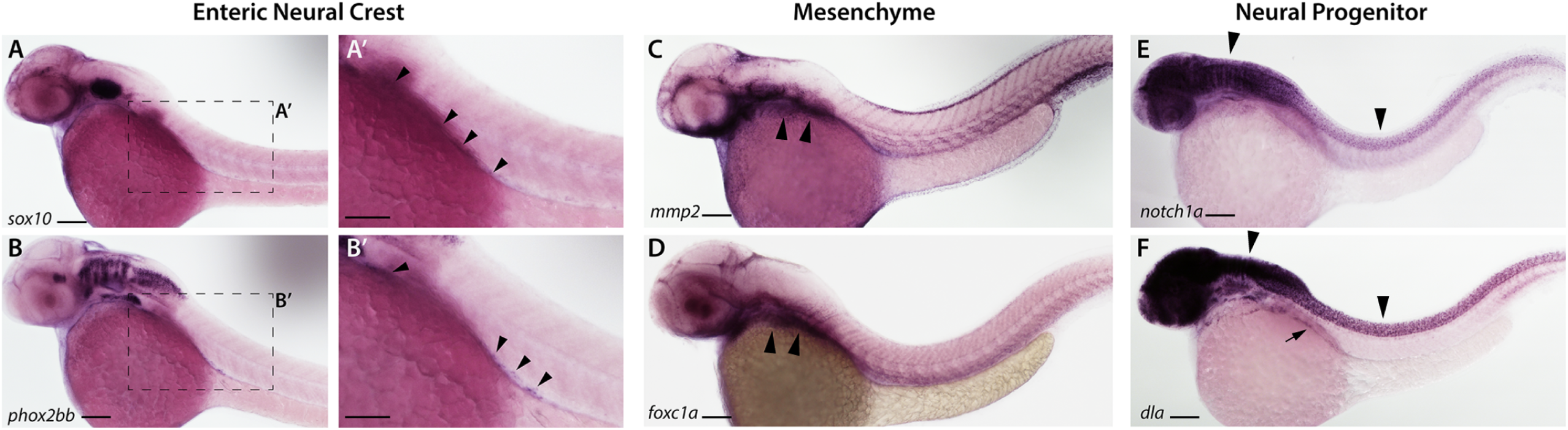
Whole mount *in situ* hybridization of select ENCC, mesenchyme and neural markers at 48-50 hpf. **(A,A’)** The marker *sox10* is shown along the vagal region and within ENCC along the foregut in (**A’**; highlighted via arrowheads). Scale bar in **A**: 60 μM, in **A’**: 40 μM. **(B, B’)** Expression of *phox2bb* is shown within the hindbrain-axial level of the embryo, as well as within ENCC within the foregut (**B’**; highlighted via arrowheads). Scale bar in **A**: 60 μM, in **A’**: 40 μM. **(C,D)** The mesenchyme markers *mmp2* (**C**, highlighted via arrowheads) and *foxc1a* (**D**; highlighted via arrowheads) are expressed within the posterior pharyngeal arches and the ventral mesenchyme. **(E,F)** The neural markers *notch1a* **(E)** and *dla* **(F)** are expressed within the hindbrain and spinal cord (arrowheads). **(F)** *dla* expression is seen in the ENCC (arrow). Scale bar in **C-F**: 60 μM

**Figure 5-figure supplement 1.**
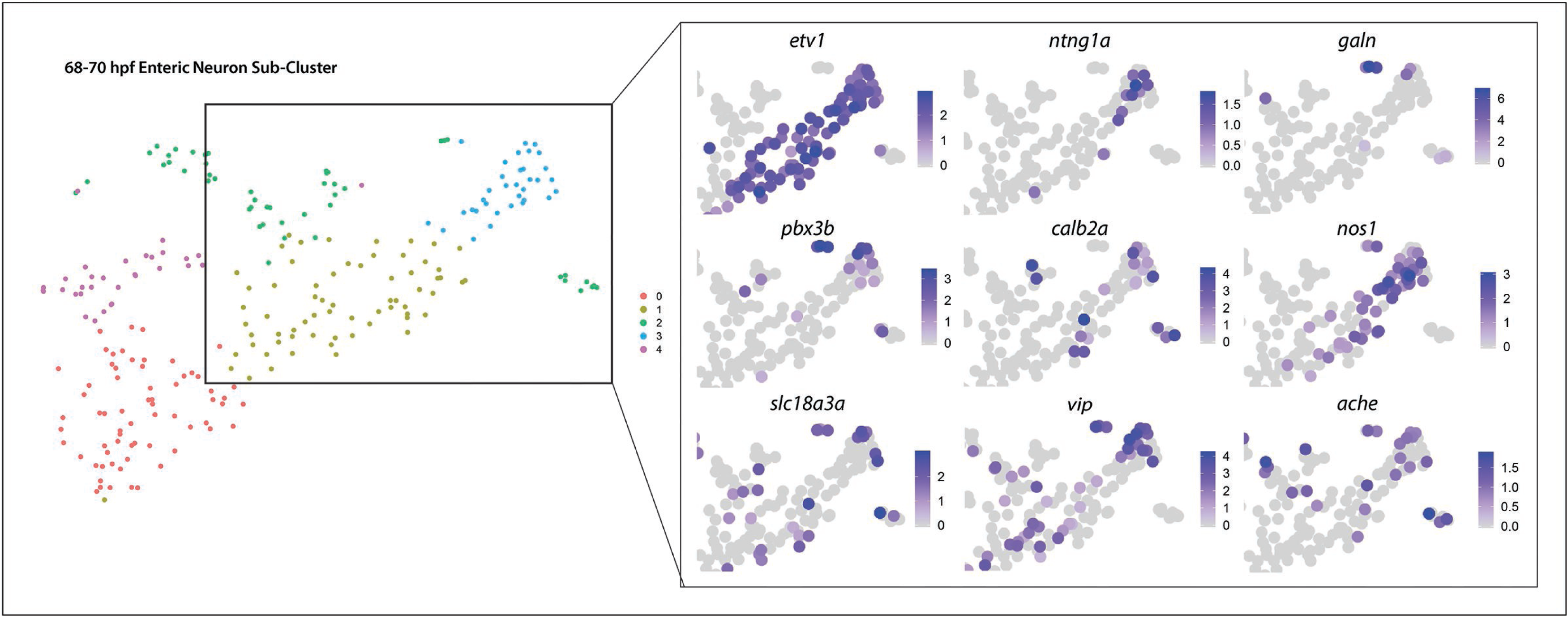
Enteric neuron subtype diversification gene expression patterns seen in enteric neuron Sub-Clusters related to Figure 5D,E Panel of tSNE feature plots magnified and cropped to focus on progressively differentiating enteric neurons (highlighted by *etv1* expression). Subtype diversification and IPAN emergence depicted via combinatorial gene expression (*etv1*, *ntng1aa*, *pbx3b*, *slc18a3a*, *calb2a*, and *ache*) localized to the distal tip of Sub-cluster 3. Inhibitory neuron markers, *nos1*, *vip*, and galanin (*galn*) were present within the pocket of diverging enteric subtypes.

**Figure 5-figure supplement 2.**
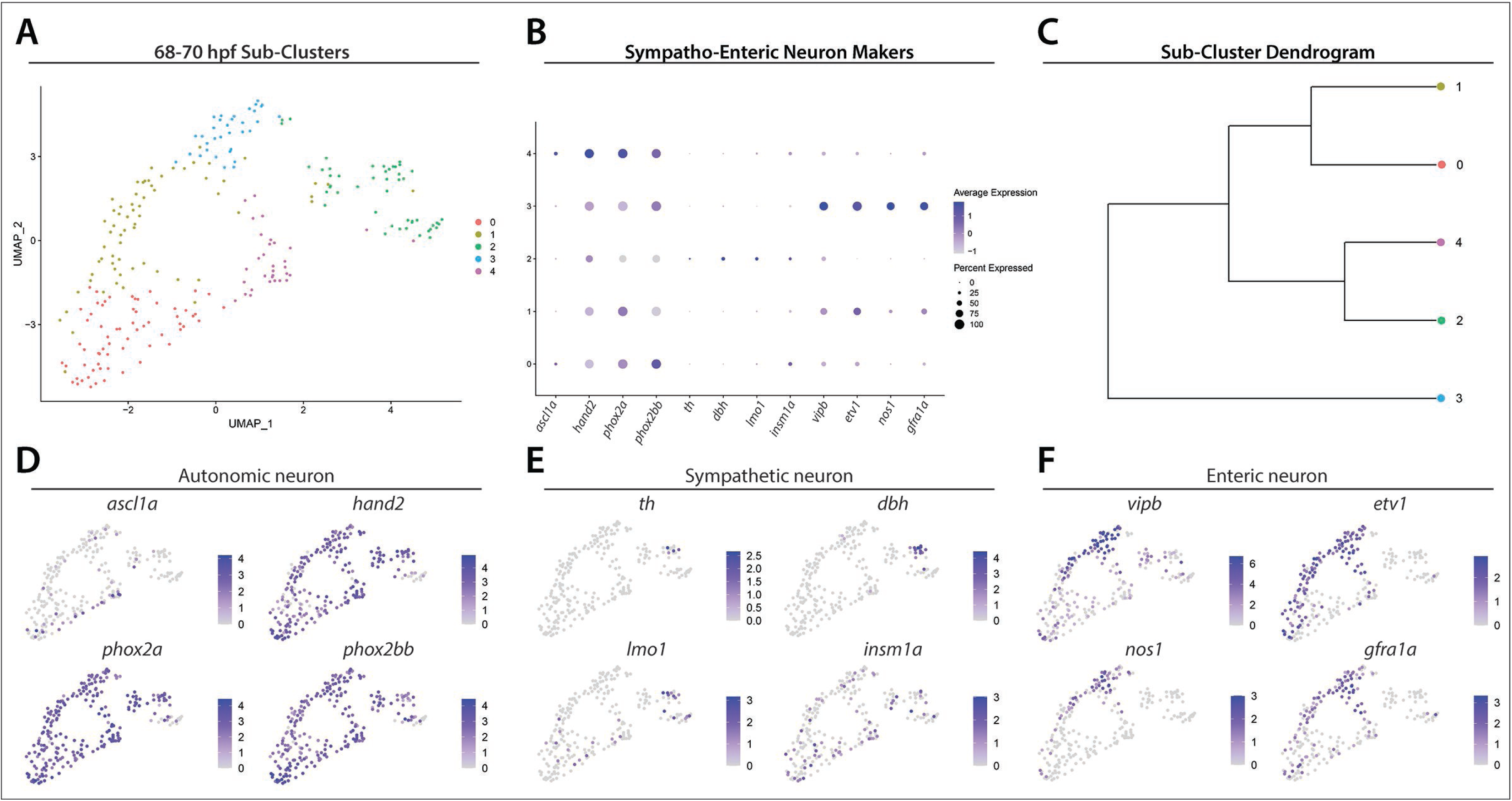
Enteric and sympathetic neuron markers distinguished among common autonomic neuron precursors. UMAP analysis of Sub-Clusters depicts enteric and sympathetic neurons delineating from a common pool of autonomic neurons. (A) UMAP plot generated following re-clustering of Clusters 5 and 12 from the 68-70 hpf data set. (B) Dot plot depicts expression levels of general autonomic neuron makers (*ascl1a, hand2, phox2a, phox2bb*), sympathetic neuron markers (*th, dbh, lmo1, insm1a*) and enteric neuron markers (*vipb, etv1, nos1, gfra1a*) within Sub-Clusters 0-4. (C) Dendrogram denotes similarity of Sub-Clusters based on average gene expression of each cell within the Sub-Clusters, which reveals transcriptomic distinction of enteric neuron Sub-Cluster 3. (**D-F**). Feature plots highlight the expression of autonomic, sympathetic and enteric neuron gene markers within UMAP Sub-Clusters.

**Figure 5-source data 1. List of marker genes per Sub-Cluster, following subset and re-clustering of enteric Cluster 5 and 12 at 68-70 hpf.**

Table reporting the Seurat output for genes for each Sub-Cluster (0-4), including p-values (<0.01), average log-fold change (.25), adjusted p-values (1.0), pct.1 summarizing proportion of cells expressing the individual gene in the sub-cluster, pct.2 showing the proportion of cells expressing the individual gene in all other sub-clusters in the sub-data set.

**Figure 5-source data 2. List of enriched pathways within enteric neuron Sub-Cluster 3 and genes present in specific opioid proenkephalin pathway identified following PANTHER Overrepresentation Test.**

Source Data table contains two sheets. Enriched Gene Pathways sheet lists the enriched pathways identified following the use of the statistical test, Fisher’s exact test that compares the number of genes in a given pathway within the reference *Danio rerio* list to the number of genes in a given pathway within the enteric neuron Sub-Cluster 3 list. This analysis produced 43 statistically overrepresented pathways. Column B: Number of genes within the given pathway present with *Danio rerio* reference list of 25888 genes. Column C: Number of genes within the given pathway present within Sub-Cluster 3 gene list. Column D: Number of pathway genes expected to be present within the Sub-Cluster 3 gene list based on the percentage of pathway genes present in the *Danio rerio* reference list. Column E: Denotes that more pathway genes were present in Sub-Cluster 3 gene list than expected. Column F: Fold enrichment of Sub-Cluster 3 pathway genes comparative to reference list. Column G: P-values calculated following Fisher’s exact test comparing expected number of pathway genes to number of genes pathway genes in Sub-Cluster 3. Column H: P-value following Benjamini-Hochberg false discovery rate (FDR) correction. Opioid proenkephalin pathway sheet lists genes associated with this pathway that are present within the enteric neuron Sub-Cluster 3 gene list, notably, opioid receptor *oprd1b*.

**Figure 6-figure supplement 1.**
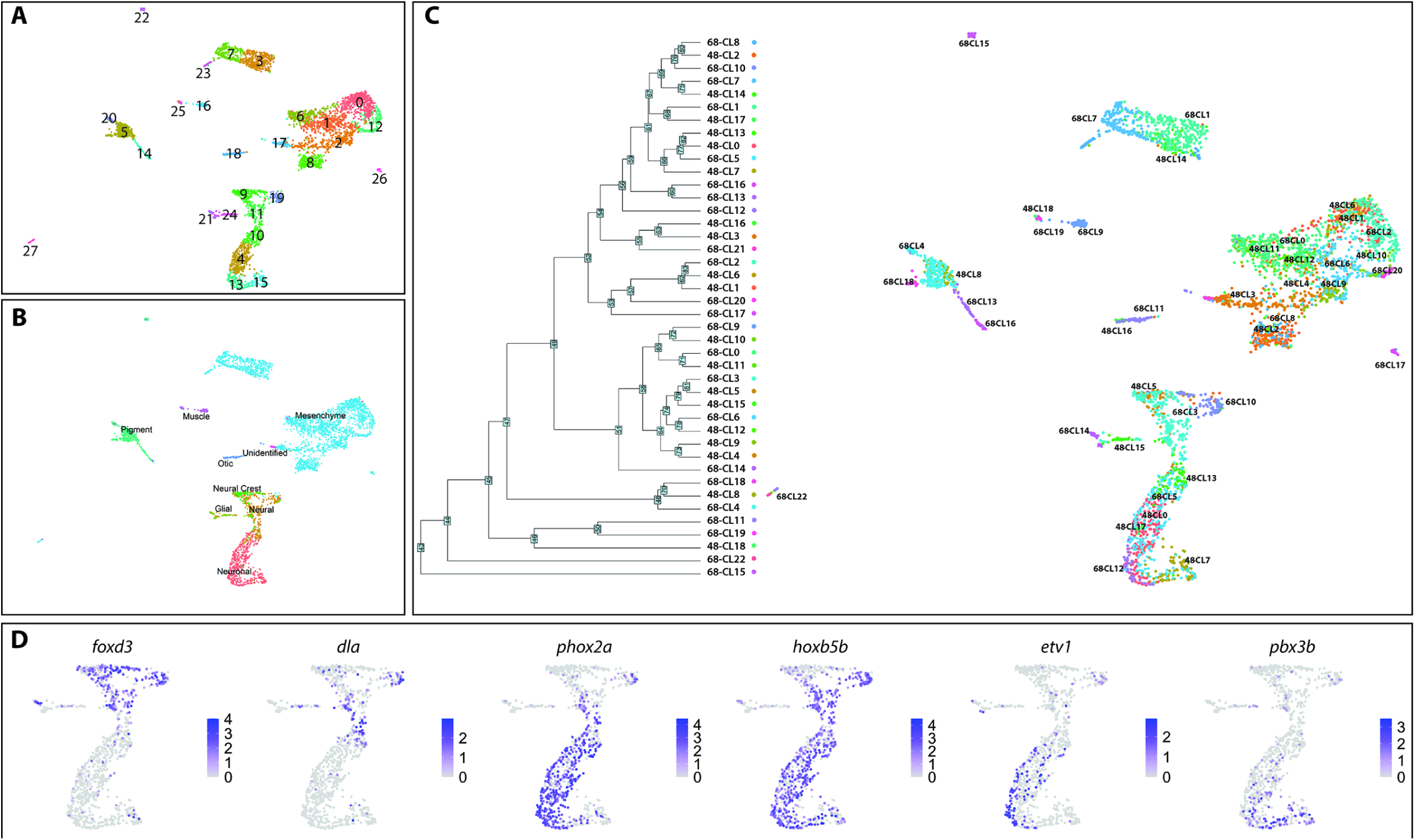
Annotated *sox10*:GFP^+^ atlas labeled by cell types. **(A)** UMAP labeling the 27 new clusters formed after the generation of the Atlas. **(B)** Following label transfer integration, major cell type classifications group together into distinct clusters. **(C)** High resolution visualization of both clustering of original cluster labels as well as their position within the UMAP. Cell categories segregate in the dendrogram largely as expected from the UMAP visualization. **(D)** Additional markers shown by UMAP for validation of the neural/neuronal clusters.

**Figure 6-figure supplement 2.**
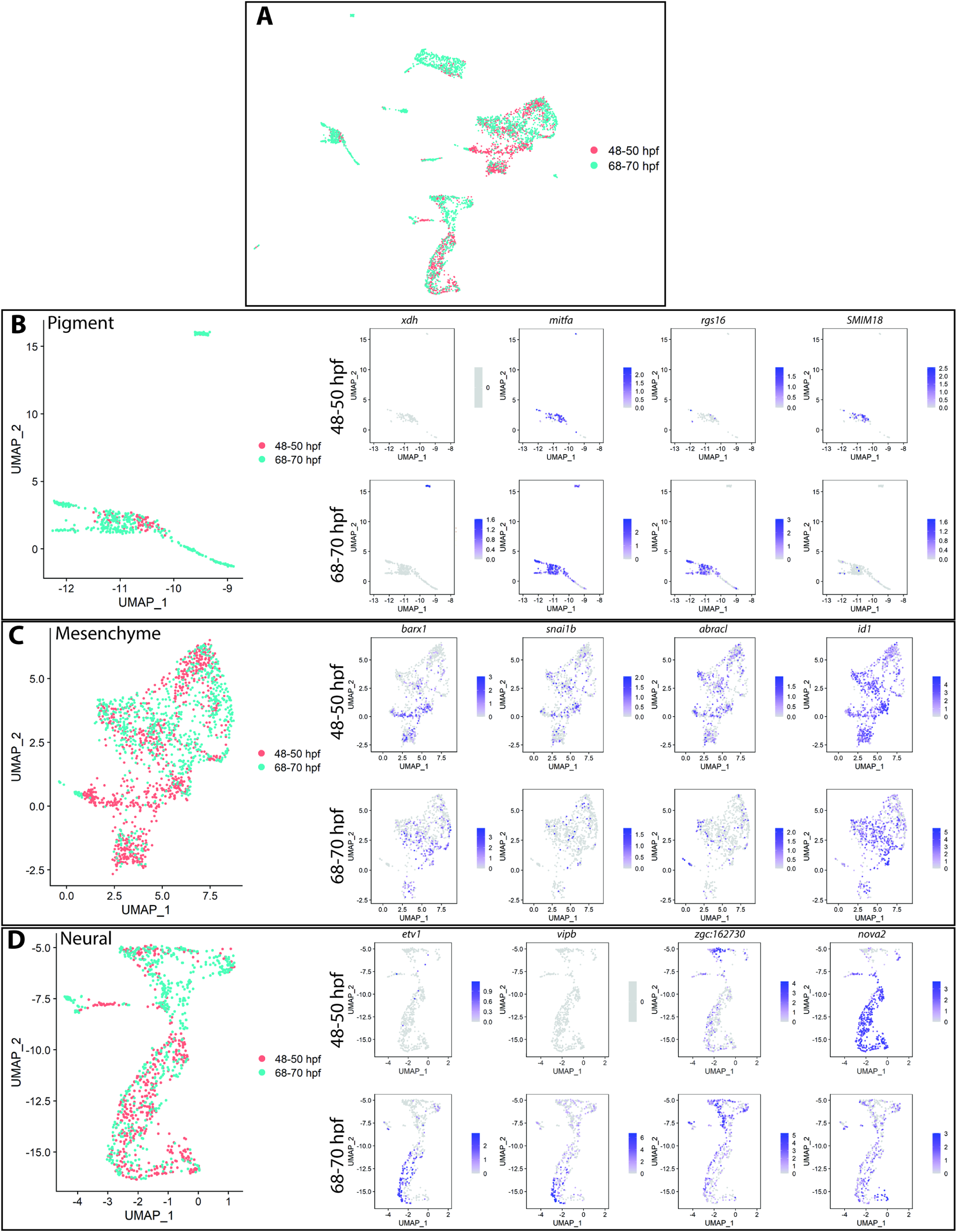
Differential expression among pigment, mesenchyme and neural/neuronal subsets of the *sox10*:GFP^+^ atlas **(A)** UMAP visualization of cells labeled by source identity (either 48-50 hpf or 68-70 hpf) following integration. All 48-50 hpf cells (pink) approximately map to a major cluster found at 68-70 hpf (aqua). **(B-D)** Subset of the pigment clusters **(B),** the mesenchyme clusters **(C)**, and the neural/neuronal clusters **(D)** highlighting several differentially expressed genes between the cells derived from each timepoint.

**Figure 6 -source data 1. List of marker genes per major cell type identity in the *sox10*:GFP^+^ merged atlas.**

Tables reporting the Seurat output for genes for each major cell type identity including p-values (<0.01), average log-fold change (≥.25), adjusted p-values (≤1.0), pct.1 summarizing proportion of cells expressing the individual gene in the cluster identity category, pct.2 showing the proportion of cells expressing the individual gene in all cluster identity categories in the atlas dataset.

**Figure 7 - figure supplement 1.**
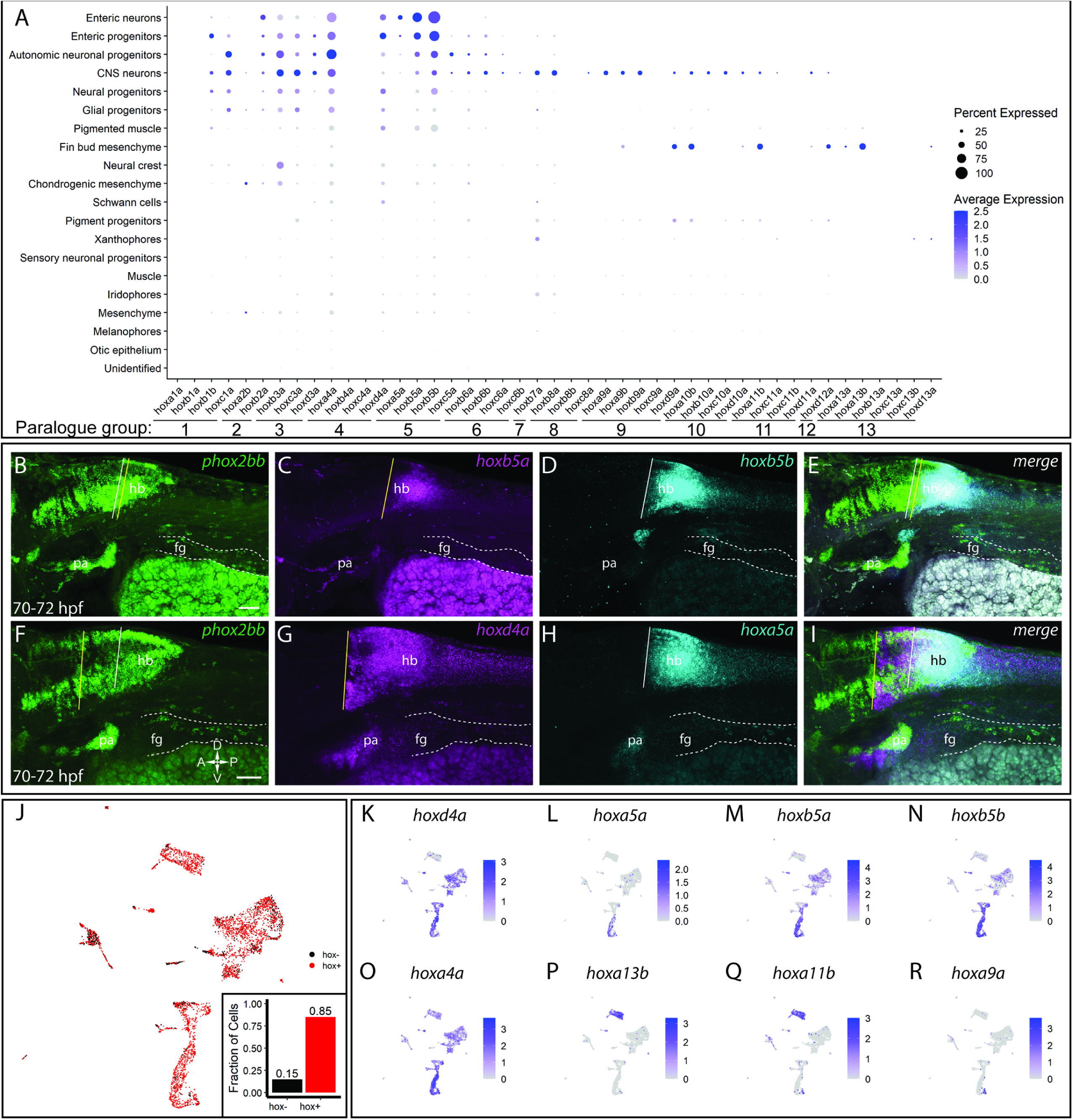
Comprehensive overview of *hox* expression profiles within the atlas. **(A)** Similar to **Figure 7A**, a comprehensive dot plot shows both the mean expression (color) as well as percent of cells per lineage (size) for zebrafish all hox genes. Discrete *hox* profiles discern specific cell types, which is particularly evident in the enteric neuronal cluster, as well as the fin bud mesenchyme. **(B-I)** Hindbrain expression patterns for enteric *hox* profiles. Autonomic neural marker *phox2bb* clearly marks the midbrain and hindbrain (hb) domains, as well as the pharyngeal arches (pa). Enterically fated cells can also clearly be seen along the tract of the gut (dash region), starting the foregut (fg). Anterior domains for *hoxb5a* **(C)**, *hoxb5b* **(D)**, *hoxd4a* **(G)**, and *hoxa5a* **(H)** are represented by a vertical line. Overlapping regions can be approximated by the merged images **(E, I)**. Scale in **(B, F)**: 50 μm; dorsoventral-rostrocaudal axis defined in **(F)**. **(J)** UMAP of the atlas demonstrating pervasive *hox* expression throughout the dataset. Expression was defined as having a log2 fold change > 0. Approximately 85% of cells contained expression for at least one *hox* gene. **(K-R)** Feature plots representing the expression of selected *hox* genes within the atlas UMAP. *hoxd4a* **(K),** *hoxa5a* **(L)**, *hoxb5a* **(M)**, *hoxb5b* **(N)** all localize in varying degrees to the enteric neural signature, though *hoxd4a* also marks the mesenchymal clusters, as seen **(A)**. *hoxa4a* **(O)** highlights its pervasive expression among the clusters. *hoxa13b* **(P)** and *hoxa11b* **(Q)** are key markers denoting the finbud mesenchyme. Alternative neural populations are further identified by their *hoxa9a* **(R)** expression profile, contrasting with the enteric populations.

## Citations

Ahn, D., and Ho, R.K. (2008). Tri-phasic expression of posterior Hox genes during development of pectoral fins in zebrafish: Implications for the evolution of vertebrate paired appendages. Dev. Biol. 322, 220–233.

Anderson, R.B., Stewart, A.L., and Young, H.M. (2006). Phenotypes of neural-crest-derived cells in vagal and sacral pathways. Cell Tissue Res. 323, 11–25.

Barlow, A.J. (1984). Neural Crest Cells in Enteric Nervous System Development and Disease. In Neural Crest Cells, (Elsevier Inc.), pp. 101–104.

Barsh, G.R., Isabella, A.J., and Moens, C.B. (2017). Vagus Motor Neuron Topographic Map Determined by Parallel Mechanisms of hox5 Expression and Time of Axon Initiation. Curr. Biol. 27, 3812–3825.

Barske, L., Askary, A., Zuniga, E., Balczerski, B., Bump, P., Nichols, J.T., and Crump, J.G. (2016). Competition between Jagged-Notch and Endothelin1 Signaling Selectively Restricts Cartilage Formation in the Zebrafish Upper Face. PLoS Genet. 12, e1005967.

Becht, E., McInnes, L., Healy, J., Dutertre, C.A., Kwok, I.W.H., Ng, L.G., Ginhoux, F., and Newell, E.W. (2019). Dimensionality reduction for visualizing single-cell data using UMAP. Nat. Biotechnol. 37, 38–47.

Bergner, A.J., Stamp, L.A., Gonsalvez, D.G., Allison, M.B., Olson, D.P., Myers, M.G., Anderson, C.R., and Young, H.M. (2014). Birthdating of myenteric neuron subtypes in the small intestine of the mouse. J. Comp. Neurol. 522, 514–527.

Bertrand, C., Chatonnet, A., Takke, C., Yan, Y.L., Postlethwait, J., Toutant, J.P., and Cousin, X. (2001). Zebrafish acetylcholinesterase is encoded by a single gene localized on linkage group 7. Gene structure and polymorphism; molecular forms and expression pattern during development. J. Biol. Chem. 276, 464–474.

Bohnsack B.L., Gallina D., Kahana A. (2011). Phenothiourea Sensitizes Zebrafish Cranial Neural Crest and Extraocular Muscle Development to Changes in Retinoic Acid and IGF Signaling. PLoS ONE 6: e22991

Bolande, R.P. (1997). Neurocristopathy: Its Growth and Development in 20 Years. Pediatr. Pathol. Lab. Med. 17, 1–25.

Brosens, E., Burns, A.J., Brooks, A.S., Matera, I., Borrego, S., Ceccherini, I., Tam, P.K., García-Barceló, M.M., Thapar, N., Benninga, M.A., et al. (2016). Genetics of enteric neuropathies. Dev. Biol. 417, 198–208.

Butler, A., Hoffman, P., Smibert, P., Papalexi, E., and Satija, R. (2018). Integrating single-cell transcriptomic data across different conditions, technologies, and species. Nat. Biotechnol. 36, 411–420.

Carney, T.J., Dutton, K.A., Greenhill, E., Delfino-Machin, M., Dufourcq, P., Blader, P., and Kelsh, R.N. (2006). A direct role for Sox10 in specification of neural crest-derived sensory neurons. Development 113, 4619–4630.

Cassar, S., Huang, X., and Cole, T. (2018). High-throughput Measurement of Gut Transit Time Using Larval Zebrafish. J. Vis. Exp. 140, e58497.

Cerdà, J., Conrad, M., Markl, J., Brand, M., and Herrmann, H. (1998). Zebrafish vimentin: Molecular characterisation, assembly properties and developmental expression. Eur. J. Cell Biol. 77, 175–187.

Choi, H.M.T., Calvert, C.R., Husain, N., Huss, D., Barsi, J.C., Deverman, B.E., Hunter, R.C., Kato, M., Lee, S.M., Abelin, A.C.T., et al. (2016). Mapping a multiplexed zoo of mRNA expression. Development 143, 3632–3637.

Choi, H.M.T., Schwarzkopf, M., Fornace, M.E., Acharya, A., Artavanis, G., Stegmaier, J., Cunha, A., and Pierce, N.A. (2018). Third-generation in situ hybridization chain reaction: multiplexed, quantitative, sensitive, versatile, robust. Development 145, dev165753.

Dash, S., and Trainor, P. (2020). The development, patterning and evolution of neural crest cell differentiation into cartilage and bone. Bone 137, 115409.

Delalande, J.M., Guyote, M.E., Smith, C.M., and Shepherd, I.T. (2008). Zebrafish sip1a and sip1b are essential for normal axial and neural patterning. Dev. Dyn. 237, 1060–1069.

Delfino-Machín, M., Madelaine, R., Busolin, G., Nikaido, M., Colanesi, S., Camargo-Sosa, K., Law, E.W.P., Toppo, S., Blader, P., Tiso, N., et al. (2017). Sox10 contributes to the balance of fate choice in dorsal root ganglion progenitors. PLoS One 12, e0172947.

DiCello, J.J., Carbone, S.E., Saito, A., Rajasekhar, P., Ceredig, R.A., Pham, V., Valant, C., Christopoulos, A., Veldhuis, N.A., Canals, M., et al. (2020). Mu and Delta Opioid Receptors Are Coexpressed and Functionally Interact in the Enteric Nervous System of the Mouse Colon. CMGH 9, 465–483.

Ding, H.L., Clouthier, D.E., and Artinger, K.B. (2013). Redundant roles of PRDM family members in zebrafish craniofacial development. Dev. Dyn. 242, 67–79.

Donica, C.L., Awwad, H.O., Thakker, D.R., and Standifer, K.M. (2013). Cellular mechanisms of nociceptin/orphanin FQ (N/OFQ) peptide (NOP) receptor regulation and heterologous regulation by N/OFQ. Mol. Pharmacol. 83, 907–918.

Le Douarin, N., and Kalcheim, C. (1999). The Neural Crest (Cambridge University Press).

Le Douarin, N.M., and Teillet, M.A.M. (1974). Experimental analysis of the migration and differentiation of neuroblasts of the autonomic nervous system and of neurectodermal mesenchymal derivatives, using a biological cell marking technique. Dev. Biol. 41, 162–184.

De Luca, A., and Coupart, L.M. (1996). Insights into Opioid Action in the Intestinal Tract (Elsevier Science Inc).

Du, J., Miller, A.J., Widlund, H.R., Horstmann, M.A., Ramaswamy, S., and Fisher, D.E. (2003). MLANA/MART1 and SILV/PMEL17/GP100 are transcriptionally regulated by MITF in melanocytes and melanoma. Am. J. Pathol. 163, 333–343.

Dutton, K.A., Pauliny, A., Lopes, S.S., Elworthy, S., Carney, T.J., Rauch, J., Geisler, R., Haffter, P., and Kelsh, R.N. (2001). Zebrafish Colourless Encodes sox10 and Specifies Non-Ectomesenchymal Neural Crest Fates. Development 128, 4113–4125.

Elworthy, S., Pinto, J.P., Pettifer, A., Cancela, M.L., and Kelsh, R.N. (2005). Phox2b function in the enteric nervous system is conserved in zebrafish and is sox10-dependent. Mech. Dev. 122, 659–669.

Epstein, M.L., Mikawa, T., Brown, A.M.C., and McFarlin, D.R. (1994). Mapping the origin of the avian enteric nervous system with a retroviral marker. Dev. Dyn. 201, 236–244.

Escot, S., Blavet, C., Faure, E., Zaffran, S., Duband, J.L., and Fournier-Thibault, C. (2016). Disruption of CXCR4 signaling in pharyngeal neural crest cells causes DiGeorge syndrome-like malformations. Development 143, 582–588.

Farnsworth, D.R., Saunders, L.M., and Miller, A.C. (2020). A single-cell transcriptome atlas for zebrafish development. Dev. Biol. 459, 100–108.

Feregrino, C., Sacher, F., Parnas, O., and Tschopp, P. (2019). A single-cell transcriptomic atlas of the developing chicken limb. BMC Genomics 20, 401.

Furness, J.B., Jones, C., Nurgali, K., and Clerc, N. (2004). Intrinsic primary afferent neurons and nerve circuits within the intestine. Prog. Neurobiol. 72, 143–164.

Gandhi, S., Ezin, M., and Bronner, M.E. (2020). Reprogramming Axial Level Identity to Rescue Neural-Crest-Related Congenital Heart Defects. Dev. Cell 53, 300–315.e4.

Ganz, J. (2018). Gut feelings: Studying enteric nervous system development, function, and disease in the zebrafish model system. Dev. Dyn. 247, 268–278.

Gaudet, P., Livstone, M.S., Lewis, S.E., and Thomas, P.D. (2011). Phylogenetic-based propagation of functional annotations within the Gene Ontology consortium. Brief. Bioinform. 12, 449–462.

Gou, Y., Guo, J., Maulding, K., and Riley, B.B. (2018). sox2 and sox3 cooperate to regulate otic/epibranchial placode induction in zebrafish. Dev. Biol. 435, 84–95.

Graham, A., Begbie, J., and McGonnell, I. (2004). Significance of the Cranial Neural Crest. Dev. Dyn. 229, 5–13.

Green, S.A., Simoes-costa, M., Bronner, M.E., and Engineering, B. (2016). Evolution of vertebrates: a view from the crest. Nature 520, 474–482.

Hall, B.K., and Hörstadius, S. (1988). The Neural Crest (London, New York, Tokyo, Toronto: Oxford University Press).

Hans, S., Irmscher, A., and Brand, M. (2013). Zebrafish Foxi1 provides a neuronal ground state during inner ear induction preceding the Dlx3b/4b-regulated sensory lineage. Development 140, 1936–1945.

Hao, M.M., and Young, H.M. (2009). Development of enteric neuron diversity. J. Cell. Mol. Med. 13, 1193–1210.

Harrison, C., Wabbersen, T., and Shepherd, I.T. (2014). In vivo visualization of the development of the enteric nervous system using a Tg(-8.3bphox2b: Kaede) transgenic zebrafish. Genesis 52, 985–990.

Heanue, T.A., and Pachnis, V. (2008). Ret isoform function and marker gene expression in the enteric nervous system is conserved across diverse vertebrate species. Mech. Dev. 125, 687– 699.

Heanue, T.A., Shepherd, I.T., and Burns, A.J. (2016). Enteric nervous system development in avian and zebrafish models. Dev. Biol. 417, 129–138.

Heffer, A., Marquart, G.D., Aquilina-Beck, A., Saleem, N., Burgess, H.A., and Dawid, I.B. (2017). Generation and characterization of Kctd15 mutations in zebrafish. PLoS One 12, e0189162.

Higdon, C.W., Mitra, R.D., and Johnson, S.L. (2013). Gene Expression Analysis of Zebrafish Melanocytes, Iridophores, and Retinal Pigmented Epithelium Reveals Indicators of Biological Function and Developmental Origin. PLoS One 8, e67801.

Hockman, D., Chong-Morrison, V., Green, S.A., Gavriouchkina, D., Candido-Ferreira, I., Ling, I.T.C., Williams, R.M., Amemiya, C.T., Smith, J.J., Bronner, M.E., et al. (2019). A genome-wide assessment of the ancestral neural crest gene regulatory network. Nat. Commun. 10, e4689.

Holmqvist, B., Ellingsen, B., Forsell, J., Zhdanova, I., and Alm, P. (2004). The early ontogeny of neuronal nitric oxide synthase systems in the zebrafish. J. Exp. Biol. 207, 923–935.

Holzer, P. (2004). Opioids and opioid receptors in the enteric nervous system: From a problem in opioid analgesia to a possible new prokinetic therapy in humans. Neurosci. Lett. 361, 192– 195.

Hong, E., Santhakumar, K., Akitake, C.A., Ahn, S.J., Thisse, C., Thisse, B., Wyart, C., Mangin, J.M., and Halpern, M.E. (2013). Cholinergic left-right asymmetry in the habenulo-interpeduncular pathway. Proc. Natl. Acad. Sci. U. S. A. 110, 21171–21176.

Hong, S.K., Tsang, M., and Dawid, I.B. (2008). The Mych gene is required for neural crest survival during zebrafish development. PLoS One 3, e2029.

Huang, V., Butler, A.A., and Lubin, F.D. (2019). Telencephalon transcriptome analysis of chronically stressed adult zebrafish. Sci. Rep. 9, 1379.

Hutchins, E.J., Kunttas, E., Piacentino, M.L., Howard, A.G.A., Bronner, M.E., and Uribe, R.A. (2018). Migration and diversification of the vagal neural crest. Dev. Biol. 444, S98–S109.

Janssens, E., Gaublomme, D., de Groef, L., Darras, V.M., Arckens, L., Delorme, N., Claes, F., van Hove, I., and Moons, L. (2013). Matrix Metalloproteinase 14 in the Zebrafish: An Eye on Retinal and Retinotectal Development. PLoS One 8, e52915.

Jarinova, O., Hatch, G., Poitras, L., Prudhomme, C., Grzyb, M., Aubin, J., Bérubé-Simard, F.-A., Jeannotte, L., and Ekker, M. (2008). Functional resolution of duplicated hoxb5 genes in teleosts. Development 135, 3543–3553.

Jowett, T., and Lettice, L. (1994). Whole-mount in situ hybridizations on zebrafish embryos using a mixture of digoxigenin- and fluorescein-labelled probes. Trends Genet. 10, 73–74.

Kague, E., Gallagher, M., Burke, S., Parsons, M., Franz-Odendaal, T., and Fisher, S. (2012). Skeletogenic Fate of Zebrafish Cranial and Trunk Neural Crest. PLoS One 7, e47394.

Kam, M.K.M., and Lui, V.C.H. (2015). Roles of Hoxb5 in the development of vagal and trunk neural crest cells. Dev. Growth Differ. 57, 158–168.

Kam, M.K.M., Cheung, M.C.H., Zhu, J.J., Cheng, W.W.C., Sat, E.W.Y., Tam, P.K.H., and Lui, V.C.H. (2014). Perturbation of Hoxb5 signaling in vagal and trunk neural crest cells causes apoptosis and neurocristopathies in mice. Cell Death Differ. 21, 278–289.

Karlsson, J., Von Hofsten, J., and Olsson, P.E. (2001). Generating transparent zebrafish: A refined method to improve detection of gene expression during embryonic development. Mar. Biotechnol. 3, 522–527.

Kelsh, R.N. (2004). Genetics and evolution of pigment patterns in fish. Pigment Cell Res. 17, 326–336.

Kelsh, R.N., and Eisen, J.S. (2000). The zebrafish colourless gene regulates development of non-ectomesenchymal neural crest derivatives. Development 127, 515–525.

Knight, R.D., Nair, S., Nelson, S.S., Afshar, A., Javidan, Y., Geisler, R., Rauch, G.J., and Schilling, T.F. (2003). Lockjaw encodes a zebrafish tfap2a required for early neural crest development. Development 130, 5755–5768.

Kuo, B.R., and Erickson, C.A. (2011). Vagal neural crest cell migratory behavior: A transition between the cranial and trunk crest. Dev. Dyn. 240, 2084–2100.

Kwak, J., Park, O.K., Jung, Y.J., Hwang, B.J., Kwon, S.H., and Kee, Y. (2013). Live image profiling of neural crest lineages in zebrafish transgenic lines. Mol. Cells 35, 255–260.

Lasrado, R., Boesmans, W., Kleinjung, J., Pin, C., Bell, D., Bhaw, L., McCallum, S., Zong, H., Luo, L., Clevers, H., et al. (2017). Lineage-dependent Spatial and Functional Organization of the Mammalian Enteric Nervous System. Science (80-.). 356, 722–726.

Lay, J., Carbone, S.E., Dicello, J.J., Bunnett, N.W., Canals, M., and Poole, D.P. (2016). Distribution and trafficking of the-opioid receptor in enteric neurons of the guinea pig. Am J Physiol Gastrointest Liver Phy-Siol 311, 252–266.

Leigh, N.R., Schupp, M.O., Li, K., Padmanabhan, V., Gastonguay, A., Wang, L., Chun, C.Z., Wilkinson, G.A., and Ramchandran, R. (2013). Mmp17b Is Essential for Proper Neural Crest Cell Migration In Vivo. PLoS One 8, e76484.

Le Lievre, C.S., and Le Douarin, N.M. (1975). Mesenchymal derivatives of the neural crest: analysis of chimaeric quail and chick embryos. J. Embryol. Exp. Morphol. 34, 125–154.

Ling, I.T.C., and Sauka-Spengler, T. (2019). Early chromatin shaping predetermines multipotent vagal neural crest into neural, neuronal and mesenchymal lineages. Nat. Cell Biol. 21, 1504–1517.

Lister, J.A. (2002). Development of pigment cells in the zebrafish embryo. Microsc. Res. Tech. 58, 435–441.

Lister, J.A., Robertson, C.P., Lepage, T., Johnson, S.L., and Raible, D.W. (1999). Nacre Encodes a Zebrafish Microphthalmia-Related Protein That Regulates Neural-Crest-Derived Pigment Cell Fate. Development 126, 3757–3767.

Lister, J.A., Lane, B.M., Nguyen, A., and Lunney, K. (2011). Embryonic expression of zebrafish MiT family genes Tfe3b, Tfeb, and Tfec. Dev. Dyn. 240, 2529–2538.

Lu, J.-K., Tsai, T.-C., Lee, H., Hsia, K., Lin, C.-H., and Lu, J.-H. (2019). Pectoral Fin Anomalies in tbx5a Knockdown Zebrafish Embryos Related to the Cascade Effect of N-Cadherin and Extracellular Matrix Formation. J. Dev. Biol. 7, 15.

Ludwig, A., Rehberg, S., and Wegner, M. (2004). Melanocyte-specific expression of dopachrome tautomerase is dependent on synergistic gene activation by the Sox10 and Mitf transcription factors. FEBS Lett. 556, 236–244.

Lumb, R., Buckberry, S., Secker, G., Lawrence, D., and Schwarz, Q. (2017). Transcriptome profiling reveals expression signatures of cranial neural crest cells arising from different axial levels. BMC Dev. Biol. 17, e5.

Luo, R., An, M., Arduini, B.L., and Henion, P.D. (2001). Specific pan-neural crest expression of zebrafish crestin throughout embryonic development. Dev. Dyn. 220, 169–174.

Martik, M.L., and Bronner, M.E. (2017). Regulatory Logic Underlying Diversification of the Neural Crest. Trends Genet. 33, 715–727.

Matini, P., Manneschi, L.I., Mayer, B., and Faussone-Pellegrini, M.S. (1995). Nitric oxide producing neurons in the human colon: an immunohistochemical and histoenzymatical study. Neurosci. Lett. 193, 17–20.

McGraw, H.F., Nechiporuk, A., and Raible, D.W. (2008). Zebrafish Dorsal Root Ganglia Neural Precursor Cells Adopt a Glial Fate in the Absence of neurogenin1. J. Neurosci. 28, 12558–12569.

Mcinnes, L., Healy, J., and Melville, J. (2018). UMAP: Uniform Manifold Approximation and Projection for Dimension Reduction. ArXiv 1802.03426v2.

Memic, F., Knoflach, V., Morarach, K., Sadler, R., Laranjeira, C., Hjerling-Leffler, J., Sundström, E., Pachnis, V., and Marklund, U. (2018). Transcription and Signaling Regulators in Developing Neuronal Subtypes of Mouse and Human Enteric Nervous System. Gastroenterology 154, 624–636.

Mi, H., Muruganujan, A., Huang, X., Ebert, D., Mills, C., Guo, X., and Thomas, P.D. (2019). Protocol Update for large-scale genome and gene function analysis with the PANTHER classification system (v.14.0). Nat. Protoc. 14, 703–721.

Minchin, J.E.N., and Hughes, S.M. (2008). Sequential actions of Pax3 and Pax7 drive xanthophore development in zebrafish neural crest. Dev. Biol. 317, 508–522.

Minoux, M., and Rijli, F.M. (2010). Molecular mechanisms of cranial neural crest cell migration and patterning in craniofacial development. Development 137, 2605–2621.

Morarach, K., Mikhailova, A., Knoflach, V., Memic, F., Kumar, R., Li, W., Ernfors, P., and Marklund, U. (2020). Diversification of molecularly defined myenteric neuron classes revealed by single cell RNA-sequencing. Nat Neurosci. https://doi.org/10.1038/s41593-020-00736-x

Nagy, N., and Goldstein, A.M. (2017). Enteric Nervous System Development: A Crest Cell’s Journey From Neural Tube to Colon. Semin. Cell Dev. Biol. 66, 94–106.

Nakamura, T., Gehrke, A.R., Lemberg, J., Szymaszek, J., and Shubin, N.H. (2016). Digits and fin rays share common developmental histories. Nature 537, 225–228.

Nord, H., Dennhag, N., Muck, J., and Von Hofsten, J. (2016). Pax7 is required for establishment of the xanthophore lineage in zebrafish embryos. Mol. Biol. Cell 27, 1853–1862.

Olden, T., Akhatar, T., Beckman, S.A., and Wallace, K.N. (2008). Differentiation of the Zebrafish Enteric Nervous System and Intestinal Smooth Muscle. Genesis 46, 484–498.

van Otterloo, E., Li, W., Garnett, A., Cattell, M., Medeiros, D.M., and Cornell, R.A. (2012). Novel Tfap2-mediated control of soxE expression facilitated the evolutionary emergence of the neural crest. Development 139, 720–730.

Parichy, D.M., Ransom, D.G., Paw, B., Zon, L.I., and Johnson, S.L. (2000). An Orthologue of the Kit-Related Gene Fms Is Required for Development of Neural Crest-Derived Xanthophores and a Subpopulation of Adult Melanocytes in the Zebrafish, Danio Rerio. Development 127, 3031–3044.

Parker, H.J., Pushel, I., and Krumlauf, R. (2018). Coupling the roles of Hox genes to regulatory networks patterning cranial neural crest. Dev. Biol. 444, S67–S78.

Parker, H.J., De Kumar, B., Green, S.A., Prummel, K.D., Hess, C., Kaufman, C.K., Mosimann, C., Wiedemann, L.M., Bronner, M.E., and Krumlauf, R. (2019). A Hox-TALE regulatory circuit for neural crest patterning is conserved across vertebrates. Nat. Commun. 10, 1182.

Petratou, K., Subkhankulova, T., Lister, J.A., Rocco, A., Schwetlick, H., and Kelsh, R.N. (2018). A Systems Biology Approach Uncovers the Core Gene Regulatory Network Governing Iridophore Fate Choice From the Neural Crest. PLOS Genet. 14, e1007402.

Petratou, K., Spencer, S.A., Kelsh, R.N., and Lister, J.A. (2019). The MITF paralog tfec is required in neural crest development for fate specification of the iridophore lineage from a multipotent pigment cell progenitor. BioRxiv 862011.

Philippidou, P., and Dasen, J.S.S. (2013). Hox Genes: Choreographers in Neural Development, Architects of Circuit Organization. Neuron 80, 12–34.

Poon, K.L., Richardson, M., Lam, C.S., Khoo, H.E., and Korzh, V. (2003). Expression pattern of neuronal nitric acid oxide synthase in embryonic zebrafish. Gene Expr. Patterns 3, 463–466.

Qu, Z.D., Thacker, M., Castelucci, P., Bagyánszki, M., Epstein, M.L., and Furness, J.B. (2008). Immunohistochemical analysis of neuron types in the mouse small intestine. Cell Tissue Res. 334, 147–161.

Quigley, I.K., and Parichy, D.M. (2002). Pigment pattern formation in zebrafish: A model for developmental genetics and the evolution of form. Microsc. Res. Tech. 58, 442–455.

R Core Team (2020). R. R A Lang. Environ. Stat. Comput. R Found. Stat. Comput.

Raffaeli, G., Cavallaro, G., Allegaert, K., Wildschut, E.D., Fumagalli, M., Agosti, M., Tibboel, D., and Mosca, F. (2017). Neonatal Abstinence Syndrome: Update on Diagnostic and Therapeutic Strategies. Pharmacother. J. Hum. Pharmacol. Drug Ther. 37, 814–823.

Rajan, S.G., Gallik, K.L., Monaghan, J.R., Uribe, R.A., Bronner, M.E., and Saxena, A. (2018). Tracking neural crest cell cycle progression in vivo. Genesis 56, e23214.

Rao, M., and Gershon, M.D. (2018). Enteric nervous system development: what could possibly go wrong? Nat. Rev. Neurosci. 19, 552–565.

Reedy, M. V., Faraco, C.D., and Erickson, C.A. (1998). Specification and migration of melanoblasts at the vagal level and in hyperpigmented silkie chickens. Dev. Dyn. 213, 476–485.

Rocha, M., Singh, N., Ahsan, K., Beiriger, A., and Prince, V.E. (2020). Neural crest development: insights from the zebrafish. Dev. Dyn. 249, 88–111.

Rodrigues, F.S.L.M., Doughton, G., Yang, B., and Kelsh, R.N. (2012). A novel transgenic line using the Cre-lox system to allow permanent lineage-labeling of the zebrafish neural crest. Genesis 50, 750–757.

Rueden, C.T., Schindelin, J., Hiner, M.C., Dezonia, B.E., Walter, A.E., Arena, E.T., and Eliceiri, K.W. (2017). ImageJ2: ImageJ for the next generation of scientific image data. BMC Bioinformatics 18, 529.

Roy-Carson, S., Natukunda, K., Chou, H., Pal, N., Farris, C., Schneider, S.Q., and Kuhlman, J.A. (2017). Defining the transcriptomic landscape of the developing enteric nervous system and its cellular environment. BMC Genomics 18, 290.

Ruzicka, L., Howe, D.G., Ramachandran, S., Toro, S., Van Slyke, C.E., Bradford, Y.M., Eagle, A., Fashena, D., Frazer, K., Kalita, P., et al. (2019). The Zebrafish Information Network: new support for non-coding genes, richer Gene Ontology annotations and the Alliance of Genome Resources. Nucleic Acids Res. 47, D867–873.

Satija, R., Farrell, J.A., Gennert, D., Schier, A.F., and Regev, A. (2015). Spatial reconstruction of single-cell gene expression data. Nat. Biotechnol. 33, 495–502.

Sauka-Spengler, T., and Bronner-Fraser, M. (2008). A gene regulatory network orchestrates neural crest formation. Nat. Rev. Mol. Cell Biol. 9, 577–568.

Saunders, L.M., Mishra, A.K., Aman, A.J., Lewis, V.M., Toomey, M.B., Packer, J.S., Qiu, X., McFaline-Figueroa, J.L., Corbo, J.C., Trapnell, C., et al. (2019). Thyroid hormone regulates distinct paths to maturation in pigment cell lineages. Elife 8, e45181.

Schindelin, J., Arganda-Carreras, I., Frise, E., Kaynig, V., Longair, M., Pietzsch, T., Preibisch, S., Rueden, C., Saalfeld, S., Schmid, B., et al. (2012). Fiji: an open-source platform for biological-image analysis. Nat. Methods 9, 676–682.

Schneider, C.A., Rasband, W.S., and Eliceiri, K.W. (2012). NIH Image to ImageJ: 25 years of image analysis. Nat. Methods 9, 671–675.

Shepherd, I.T., Pietsch, J., Elworthy, S., Kelsh, R.N., and Raible, D.W. (2004). Roles for GFRα1 receptors in zebrafish enteric nervous system development. Development 131, 241–249.

Simoes-Costa, M., and Bronner, M.E. (2016). Reprogramming of avian neural crest axial identity and cell fate. Science (80-.). 352, 1570–1573.

Simões-Costa, M., Tan-Cabugao, J., Antoshechkin, I., Sauka-Spengler, T., and Bronner, M.E. (2014). Transcriptome analysis reveals novel players in the cranial neural crest gene regulatory network. Genome Res. 24, 281–290.

Singleman, C., and Holtzman, N.G. (2014). Growth and maturation in the zebrafish, Danio Rerio: A staging tool for teaching and research. Zebrafish 11, 396–406.

Sobczak, M., Sałaga, M., Storr, M.A., and Fichna, J. (2014). Physiology, signaling, and pharmacology of opioid receptors and their ligands in the gastrointestinal tract: Current concepts and future perspectives. J. Gastroenterol. 49, 24–45.

Soldatov, R., Kaucka, M., Kastriti, M.E., Petersen, J., Chontorotzea, T., Englmaier, L., Akkuratova, N., Yang, Y., Häring, M., Dyachuk, V., et al. (2019). Spatio-temporal structure of cell fate decisions in murine neural crest. Science (80-.). 364, eaas9536.

Sperber, S.M., and Dawid, I.B. (2008). barx1 is necessary for ectomesenchyme proliferation and osteochondroprogenitor condensation in the zebrafish pharyngeal arches. Dev. Biol. 321, 101–110.

Sperber, S.M., Saxena, V., Hatch, G., and Ekker, M. (2008). Zebrafish dlx2a Contributes to Hindbrain Neural Crest Survival, Is Necessary for Differentiation of Sensory Ganglia and Functions With dlx1a in Maturation of the Arch Cartilage Elements. Dev. Biol. 314, 59–70.

Stewart, R.A., Arduini, B.L., Berghmans, S., George, R.E., Kanki, J.P., Henion, P.D., and Look, A.T. (2006). Zebrafish foxd3 Is Selectively Required for Neural Crest Specification, Migration and Survival. Dev. Biol. 292, 174–188.

Strausberg, R.L., Feingold, E.A., Grouse, L.H., Derge, J.G., Klausner, R.D., Collins, F.S., Wagner, L., Shenmen, C.M., Schuler, G.D., Altschul, S.F., et al. (2002). Generation and initial analysis of more than 15,000 full-length human and mouse cDNA sequences. Proc. Natl. Acad. Sci. U. S. A. 99, 16899–16903.

Stuart, T., Butler, A., Hoffman, P., Hafemeister, C., Papalexi, E., Mauck, W.M., Hao, Y., Stoeckius, M., Smibert, P., and Satija, R. (2019). Comprehensive Integration of Single-Cell Data. Cell 177, 1888–1902.e21.

Tambalo, M., Mitter, R., and Wilkinson, D.G. (2020). A single cell transcriptome atlas of the developing zebrafish hindbrain. Dev. 147, dev184143.

Taylor, C.R., Montagne, W.A., Eisen, J.S., and Ganz, J. (2016). Molecular fingerprinting delineates progenitor populations in the developing zebrafish enteric nervous system. Dev. Dyn. 245, 1081–1096.

Theodore, L.N., Hagedorn, E.J., Cortes, M., Natsuhara, K., Liu, S.Y., Perlin, J.R., Yang, S., Daily, M.L., Zon, L.I., and North, T.E. (2017). Distinct Roles for Matrix Metalloproteinases 2 and 9 in Embryonic Hematopoietic Stem Cell Emergence, Migration, and Niche Colonization. Stem Cell Reports 8, 1226–1241.

Theveneau, E., and Mayor, R. (2012). Neural crest delamination and migration: From epithelium-to-mesenchyme transition to collective cell migration. Dev. Biol. 366, 34–54.

Thisse, B., and Thisse, C. (2004). Fast Release Clones: A High Throughput Expression Analysis. ZFIN Direct Data Submission. (http://zfin.org).

Thisse, C., and Thisse, B. (2005). High Throughput Expression Analysis of ZF-Models Consortium Clones. ZFIN Direct Data Submission. http://zfin.org.

Thisse, B., Pflumio, S., Fürthauer, M., Loppin, B., Heyer, V., Degrave, A., Woehl, R., Lux, A., Steffan, T., Charbonnier, X.., et al. (2001). Expression of the zebrafish genome during embryogenesis. ZFIN Direct Data Submission. (http://zfin.org).

Uribe, R.A., and Bronner, M.E. (2015). Meis3 is required for neural crest invasion of the gut during zebrafish enteric nervous system development. Mol. Biol. Cell 26, 3728–3740.

Uyttebroek, L., Shepherd, I.T., Harrisson, F., Hubens, G., Blust, R., Timmermans, J.P., and van Nassauw, L. (2010). Neurochemical coding of enteric neurons in adult and embryonic zebrafish (Danio rerio). J. Comp. Neurol. 518, 4419–4438.

Vega-Lopez, G.A., Cerrizuela, S., and Aybar, M.J. (2017). Trunk neural crest cells: formation, migration and beyond. Int. J. Dev. Biol. 61, 5–15.

Wagner, D.E., Weinreb, C., Collins, Z.M., Briggs, J.A., Megason, S.G., and Klein, A.M. (2018). Single-cell mapping of gene expression landscapes and lineage in the zebrafish embryo. Science (80-.). 360, 981–987.

Wang, H.H., Chen, H.S., Li, H.B., Zhang, H., Mei, L.Y., He, C.F., Wang, X.W., Men, M.C., Jiang, L., Liao, X. Bin, et al. (2014). Identification and functional analysis of a novel mutation in the SOX10 gene associated with Waardenburg syndrome type IV. Gene 538, 36–41.

Wang, W. Der, Melville, D.B., Montero-Balaguer, M., Hatzopoulos, A.K., and Knapik, E.W. (2011). Tfap2a and Foxd3 regulate early steps in the development of the neural crest progenitor population. Dev. Biol. 360, 173–185.

Williams, A.L., and Bohnsack, B.L. (2015). Neural crest derivatives in ocular development: Discerning the eye of the storm. Birth Defects Res. Part C Embryo Today Rev. 105, 87–95.

Williams, R.M., Candido-Ferreira, I., Repapi, E., Gavriouchkina, D., Senanayake, U., Ling, I.T.C., Telenius, J., Taylor, S., Hughes, J., and Sauka-Spengler, T. (2019). Reconstruction of the Global Neural Crest Gene Regulatory Network In Vivo. Dev. Cell 51, 522–576.e7.

Wilson, Y.M., Richards, K.L., Ford-Perriss, M.L., Panthier, J.J., and Murphy, M. (2004). Neural crest cell lineage segregation in the mouse neural tube. Development 131, 6153–6162.

Wood, J.D., and Galligan, J.J. (2004). Function of opioids in the enteric nervous system. Neurogastroenterol. Motil. 16, 17–28.

Yelon, D., Brauch, T., Halpern, M.E., Ruvisnsky, I., Ho, R.K., Silver, L.M., and Stainier, D.Y.R. (2000). The bHLH transcription factor Hand2 plays parallel roles in zebrafish heart and pectoral fin development. Development 127, 2573–2582.

Yntema, C.L., and Hammond, W.S. (1954). The origin of intrinsic ganglia of trunk viscera from vagal neural crest in the chick embryo. J. Comp. Neurol. 101, 515–541.

Zheng, G.X.Y., Terry, J.M., Belgrader, P., Ryvkin, P., Bent, Z.W., Wilson, R., Ziraldo, S.B., Wheeler, T.D., Mcdermott, G.P., Zhu, J., et al. (2017). Massively parallel digital transcriptional profiling of single cells. Nat. Commun. 8, 14049.

Zoli, M. (2000). Distribution of Cholinergic Neurons in the Mammalian Brain with Special Reference to their Relationship with Neuronal Nicotinic Acetylcholine Receptors. In Neuronal Nicotinic Receptors. Handbook of Experimental Pharmacology, F. Clementi, D. Fornasari, and C. Gotti, eds. (Berlin, Heidelberg: Springer), pp. 13–30.

